# Predicting response to cytotoxic chemotherapy

**DOI:** 10.1101/2023.01.28.525988

**Authors:** Joe Sneath Thompson, Laura Madrid, Barbara Hernando, Carolin M. Sauer, Maria Vias, Maria Escobar-Rey, Wing-Kit Leung, Jamie Huckstep, Magdalena Sekowska, Karen Hosking, Mercedes Jimenez-Linan, Marika A. V. Reinius, Harry Dobson, Dilrini De Silva, Ángel Fernández-Sanromán, Deborah Sanders, Filipe Correia Martins, Miguel Quintela-Fandino, Florian Markowetz, Jason Yip, James D Brenton, Anna M Piskorz, Geoff Macintyre

## Abstract

Cytotoxic chemotherapies have been a crucial part of cancer treatment for over 40 years. While their primary target is cancer cells, they can also harm normal cells, resulting in dose-limiting toxicity. Most chemotherapies were approved before the advent of precision biomarkers, as such, many patients experience severe toxic side effects without any benefit. To address this challenge, we have developed three precision biomarkers to predict response to platins, taxanes, and anthracyclines. Based on chromosomal instability (CIN) signatures, these biomarkers can be computed from a single genomic test. For platins and taxanes, we used CIN signatures related to impaired homologous recombination, while for anthracyclines, we discovered a CIN signature representing micronuclei induction which predicts resistance. In a clinical study involving 41 high-grade serous ovarian cancers, patients predicted to be sensitive by these biomarkers showed significantly prolonged progression-free survival. To further validate the effectiveness of the taxane and anthracycline predictors, we conducted a retrospective randomised control study involving 182 ovarian and 219 breast cancer patients. Patients predicted as resistant showed increased risk of time to treatment failure compared to standard of care, hazard ratios of 1.73 (95%CI=0.98-3.07) for taxane in ovarian, 3.67 (95%CI=2.12-6.34) for taxane in breast, and 1.93 (95%CI=1.22-3.04) for doxorubicin in ovarian. We also found that liquid biopsies can be used to make these predictions in up to 30% of ovarian cancer patients. Our findings highlight the clinical value of CIN signatures in predicting treatment response to various chemotherapies across multiple different types of cancer. The ability to quantify multiple CIN signature biomarkers using a single genomic test offers a unified approach to guide treatment decisions for cytotoxic chemotherapies. Ultimately, this has the potential to transform the current one-size-fits-all chemotherapy approach into a more precise and tailored form of medicine.

## Introduction

Cytotoxic chemotherapies exploit the defective properties of a cancer cell, such as impaired DNA repair mechanisms, to preferentially drive cancer cells to programmed cell death^1^. However, most chemotherapies also have detrimental effects on healthy cells, potentially causing severe side effects despite administration alongside modern day supportive care^2^. Many of these agents were approved for clinical use before the adoption of therapy selection biomarkers, which is in contrast to new targeted therapies that increasingly require the presence of a companion diagnostic test to determine whether a patient is eligible for treatment^3^. Therefore, a large number of patients treated with cytotoxics will experience extreme side effects with no benefit from the therapy. Enabling precision use of these agents could allow patients resistant to a therapy to avoid unnecessary side effects. This may also enable them to receive an alternative therapy, ultimately improving overall health outcomes. Furthermore, precision use of cytotoxics can reduce healthcare costs by lowering expenditure on ineffective cancer therapies and additional medical interventions for treatment complications.

We have recently developed a new type of biomarker that has the potential to predict response to multiple cytotoxic chemotherapies. This technology encodes the genome-wide imprints of distinct types of chromosomal instability (CIN) operating in a tumour genome using CIN signatures^4,5^. These signatures represent the activity of different defective pathways in a tumour that cause characteristic patterns of DNA copy number change. Initially prototyped in ovarian cancer^5^ and then extended pan-cancer^4^, we have previously demonstrated their potential to predict response to platinum-based chemotherapy in patients and taxane response *in vitro*^*4*^. As the full spectrum of signatures can be quantified in a tumour using a single genomic test, we hypothesise that CIN signatures might be used to predict response to multiple cytotoxic chemotherapies at diagnosis.

In this study we present a single, integrated approach to predict response to three of the most commonly used types of chemotherapies: platins, taxanes and anthracyclines, using both tumour tissue samples and cfDNA samples obtained from liquid biopsies.

## Results

### Identifying an anthracycline response predictor

Like many other genotoxic chemotherapies, anthracyclines can cause DNA damage resulting in extrachromosomal DNA (ecDNA) encapsulated in micronuclei^6^. When micronuclei rupture and release their contents into the cytoplasm, it can trigger the activation of cGAS-STING signalling, resulting in proinflammatory signalling through type I interferon^7^. It has also been established that such immune system activation is crucial for the success of anthracycline treatments^8^. However, how tumours resist anthracycline treatment is less well known.

Tumours exposed to chronic cGAS-STING activation have been shown to undergo a switch to noncanonical NF-KB signalling, ultimately promoting metastasis and immune evasion^9^. Therefore, it is possible that tumours resistant to anthracyclines may tolerate the ongoing formation of micronuclei via this switching mechanism. This switching mechanism is seen as an important bottleneck during tumour evolution^7^ and may represent a vital distinction between anthracycline sensitive and resistant tumours. As the amplified DNA commonly found in micronuclei can be incorporated back into the genome as homogeneously staining regions (HSRs)^10^, it may be possible to identify tumours which have survived this evolutionary bottleneck from their genomes. As ovarian CIN signature 6 and pan-cancer signatures CX8, CX9 and CX13 represent focal amplifications linked to ecDNA^45^, we hypothesised these could be used to identify micronuclei tolerant and thus anthracycline resistant tumours.

To test if the presence of these signatures was associated with any modulation in micronuclei formation rates, we treated a panel of four ovarian cancer cell lines with low dose doxorubicin and observed micronuclei induction rates using fluorescent imaging (**Figure 1a)**. We sequenced the genomes of the cell lines prior to treatment, computed ovarian CIN signatures and also estimated the expected number of induced micronuclei using a model of micronuclei induction and inheritance (see Methods). Cell lines with high ovarian CIN signature 6 showed fewer than expected micronuclei, whereas the cell line with no signature 6 showed the expected number of micronuclei (**Figure 1b**). This suggests cells with signature 6 have a reduction in DNA damage and potential genome stabilisation, likely mediated by the switch to noncanonical NF-KB signalling which in turn can activate homologous recombination^11^.

**Figure 1.**
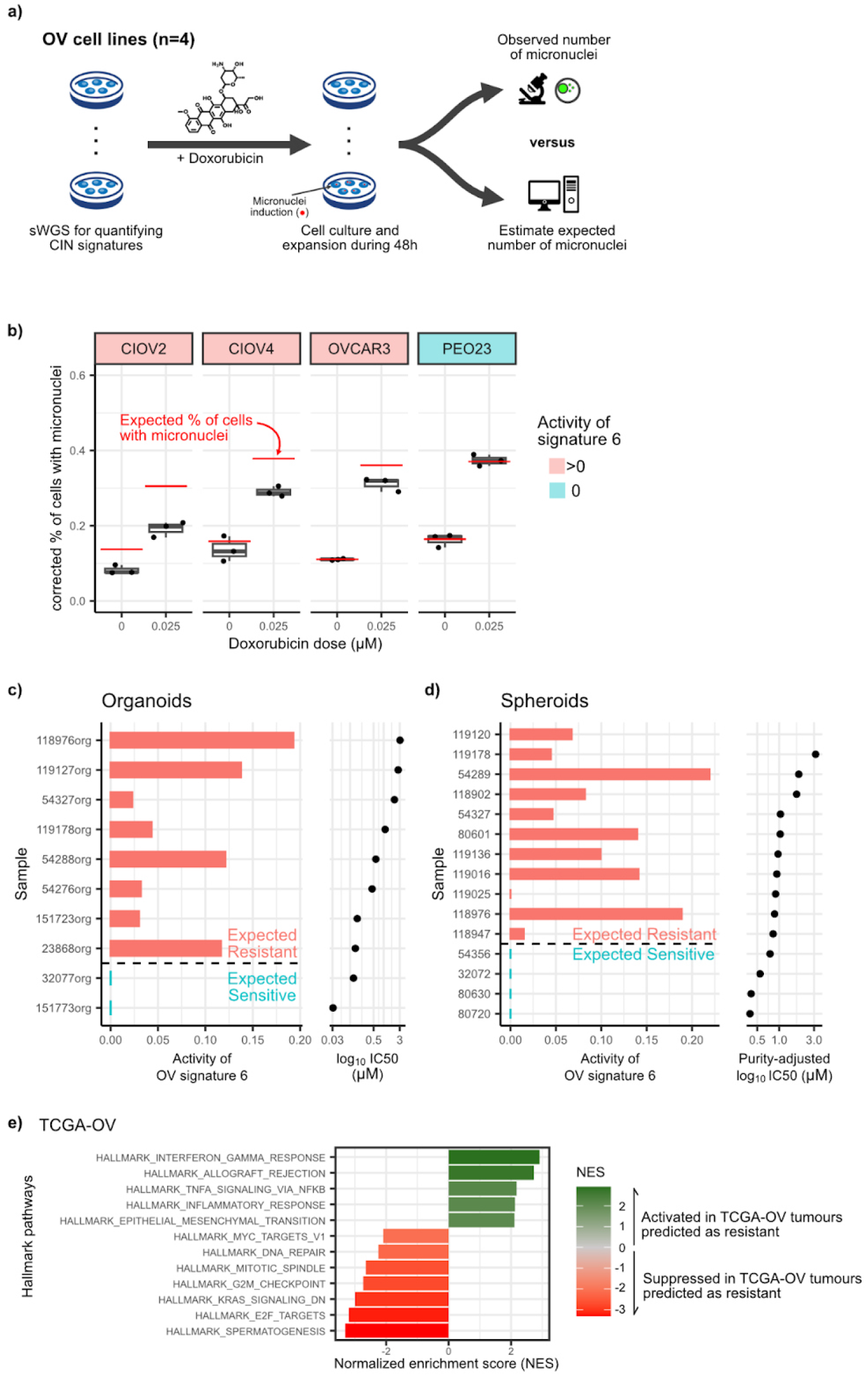
Predicting doxorubicin response using copy number signatures linked to extrachromosomal DNA. **a)** Overview of experimental design for exploring the presence of ovarian copy number signature 6 and micronuclei induction. **b**) Evaluation of micronuclei induction under doxorubicin treatment. Boxplots show the observed frequency of cells with micronuclei (y-axis) in the presence of 0 and 0.025pM of doxorubicin. Red line indicates the expected frequency of cells with micronuclei according to growth rates and micronuclei persistence across cell divisions. **c**) Ovarian copy number signature 6 activities (x-axis) across 10 patient derived organoids (y-axis). Organoids are colour coded based on their expected sensitivity (blue) or resistance (red) to *in vitro* doxorubicin treatment based on donor patient response to first-line treatment. **d**) Ovarian copy number signature 6 activities (x-axis) across 15 patient derived spheroids harvested from ascitic fluid (y-axis). Spheroids are colour coded based on their expected sensitivity (blue) or resistance (red) to *in vitro* doxorubicin treatment. **e**) Gene set enrichment analysis results showing HALLMARK gene sets that are highly enriched in TCGA-OV tumours predicted as resistant compared with those predicted as sensitive to doxorubicin. Only significant results after FDR correction are shown (q-value < 0.05)

We next sought to determine whether ovarian CIN signature 6 was directly predictive of doxorubicin treatment response *in vitro. We* treated 10 ovarian cancer organoids with doxorubicin, measured response via IC50 and observed signature 6 activity computed from low-pass WGS of the organoids prior to treatment (**Figure 1c**). The two most sensitive organoids had >0 signature 6 activity whereas the remainder had >0 activity. To further refine this observation, we attempted to estimate the fraction of the organoids that could be considered sensitive to doxorubicin based on the observed sensitivity to first-line platinum treatment in the donor patients, as patients resistant to platinum chemotherapy are expected to have an 18% response rate to doxorubicin monotherapy^12-18^, whereas sensitive patients a 28% response rate^19^. Using the observed platinum sensitivity rates (>6 months recurrence free, summarised in **Supplementary Table 3**), we estimated that 2 of the 10 organoids could be considered sensitive to doxorubicin and 8 resistant (Methods, **Figure 1c)**. Thus a threshold of >0 in signature 6 activity accurately predicted the sensitivity status of the organoids to doxorubicin treatment (p=0.022, permutation test).

To validate this doxorubicin response predictor we used an alternative, 3D *in vitro* treatment system^6^. We harvested tumour spheroids from 15 patient’s ascitic fluid and performed short term culture, treating them with doxorubicin and observing response via IC50. Similar to the organoids, the expected number of sensitive spheroids was estimated using clinical response data from the donor patients, with 4 spheroids considered sensitive and 11 resistant (**Figure 1d**). In this cohort, all sensitive spheroids showed 0 signature 6 activity, indicating 100% sensitivity to identify doxorubicin responders. However, 1 of the resistant spheroids also showed 0 signature 6 activity, suggesting 90% specificity for identifying responsive spheroids (p=0.004, permutation test). We also used our pan-cancer CIN signatures to identify responsive cases in both organoids and spheroids (CX8, CX9 or CX13 > 0.01 activity) and found an overall sensitivity of 100% and specificity of 79% (**Extended Data Figure 1)**.

Finally, we sought evidence that tumours predicted to be resistant to doxorubicin treatment had undergone the switch from cGAS-STING to non-canonical NF-KB signalling. Ovarian patient samples from the TOGA predicted as resistant showed enrichment of interferon and NF-KB signalling and epithelial-to-mesenchymal transition (EMT) (**Figure 1e)**. Together, these data support a model of anthracycline resistance that is mediated by suppressed micronuclei induction and chronic cGAS-STING activation leading to non-canonical NF-KB signalling mediated immune suppression and genome stabilisation. Importantly, these tumours which have acquired resistance can be identified using CIN signatures representing focal amplifications.

### Predicting chemotherapy response in ovarian cancer

Current ovarian cancer treatment relies heavily on cytotoxic chemotherapy with standard of care typically being a combination of carboplatin/paclitaxel and primary or interval debulking surgery (first-line of treatment), and combinations of carboplatin, paclitaxel, and doxorubicin at second and subsequent lines of treatment. The recent introduction of PARP inhibitors as a maintenance therapy following first-line treatment has improved patient outcomes^20^, however, first and second-line treatments remain unchanged. Being able to predict doxorubicin response as well as platinum and paclitaxel from a single CIN signature test therefore has the potential to enable precision medicine for all standard of care therapies used in the treatment of high-grade serous ovarian cancer (HGSOC).

In our previous work, we showed that CIN signatures CX3 and CX5 were linked with response to taxane treatment *in vitro* across 297 cancer cell lines^4^. In order to adapt these signatures for prediction in patients, we fitted a linear model and observed that CX5 explained more of the treatment response variance than CX3 (see Methods for further details, **Supplementary Table 2)**, suggesting CX5 may be a superior predictor of taxane response. To convert the CX5 signature activity to a binary response predictor we opted for a threshold approach, whereby the threshold is optimised on a training cohort, or when a cohort is not available it is selected at the point representing the prior probability of response across a relevant patient cohort. For example, as 20-40% of metastatic breast cancer patients are expected to respond to taxane treatment^21^, a threshold at 30% of CX5 activity across the cohort will likely effectively separate sensitive and resistant patients. Also included in our previous work was a retrospective analysis of ovarian and esophageal cancers showing the potential of CX2 and CX3 to predict response to platinum treatment^4^. One limitation of this analysis was that overall survival was used as a surrogate for response. Here, we aimed to assess performance using the more direct measure of progression-free survival (PFS).

To assess the feasibility of an integrated platinum/taxane/anthracycline response predictor, we assembled a retrospective real-world study cohort using patients enrolled in the OV04 clinical study. OV04 is an ongoing observational study that records patient clinical data and collects patient material for the purpose of biomarker and scientific discovery. We identified 41 patients with available tumour samples where we could assess predictive performance after first-line platinum treatment, second-line doxorubicin treatment and second-line paclitaxel treatment (**Extended Data Figure 2)**. We performed a comprehensive annotation of the clinical data from the OV04 cohort to accurately calculate PFS (see Methods). This enabled us to directly link CIN signatures to the response of the three mainstay treatments for ovarian cancers.

Classification of the 41 patients into platinum-sensitive and platinum-resistant groups yielded a significant difference in PFS intervals, taking into account both age at diagnosis and tumour stage as covariates (hazard ratio=0.379, 95% CI=0.180-0.799, p=0.011, **Figure 2a)**.

**Figure 2.**
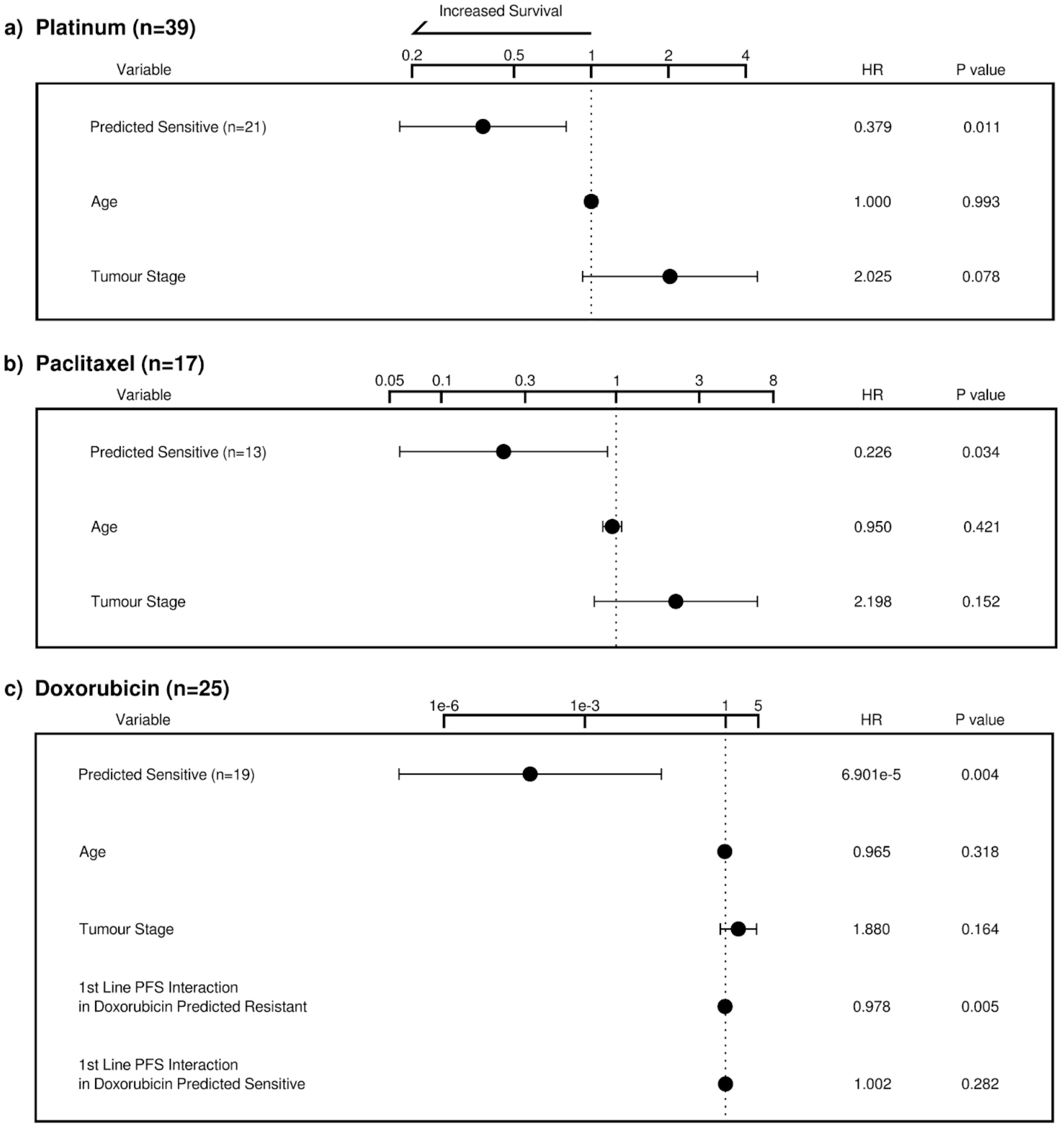
Performance assessment of platinum, paclitaxel and doxorubicin response prediction in ovarian cancer using progression-free survival. **a)** Cox proportional hazards model results for predicting platinum sensitivity accounting for age at diagnosis and tumour stage. **b**) Cox proportional hazards model results for predicting paclitaxel sensitivity accounting for age at diagnosis and tumour stage. **c**) Cox proportional hazards model results for predicting doxorubicin resistance accounting for age at diagnosis, tumour stage, and the interaction between the response prediction and PFS from 1st line platinum treatment.

Classification of the 17 out of 41 patients that received paclitaxel treatment at second-line into sensitive and resistant groups also showed significant differences in PFS after correcting for age at diagnosis and tumour stage (hazard ratio=0.226, 95% CI=0.057-0.892, p=0.034, Figure 2b). Classification of the 25 out of 41 patients that received doxorubicin treatment at second-line into sensitive and resistant groups showed significant differences in PFS after controlling for first-line platinum PFS, age at diagnosis and tumour stage (hazard ratio=6.901×10^−5^, 95%CI=1.108×10^− 7^-0.043, p=0.004, **Figure 2c**).

While these results are highly encouraging, there is a limitation: the predictions were made across a cohort of patients that all received the therapy. Therefore, there is a possibility that the classification may be prognostic, rather than predictive. To truly assess predictive performance of the biomarkers, a randomised controlled study design would be required^22^.

### Predicting chemotherapy response pan-cancer

To address this challenge we annotated time to treatment failure (TTF) intervals for cancer-specific cohorts of the Cancer Genome Atlas (TCGA) study and selected tumour types with a sufficient number of samples to enable a retrospective randomised controlled study design (**Figure 3a**). Briefly, patients within predicted resistant and predicted sensitive groups were retrospectively allocated to the experimental arm (treated with single-agent chemotherapy) or to the control arm (treated with standards-of-care therapies). There were sufficient patient numbers to perform this analysis for ovarian and breast patients treated with taxanes, and ovarian patients treated with doxorubicin (**Extended Data Figure 3**).

**Figure 3.**
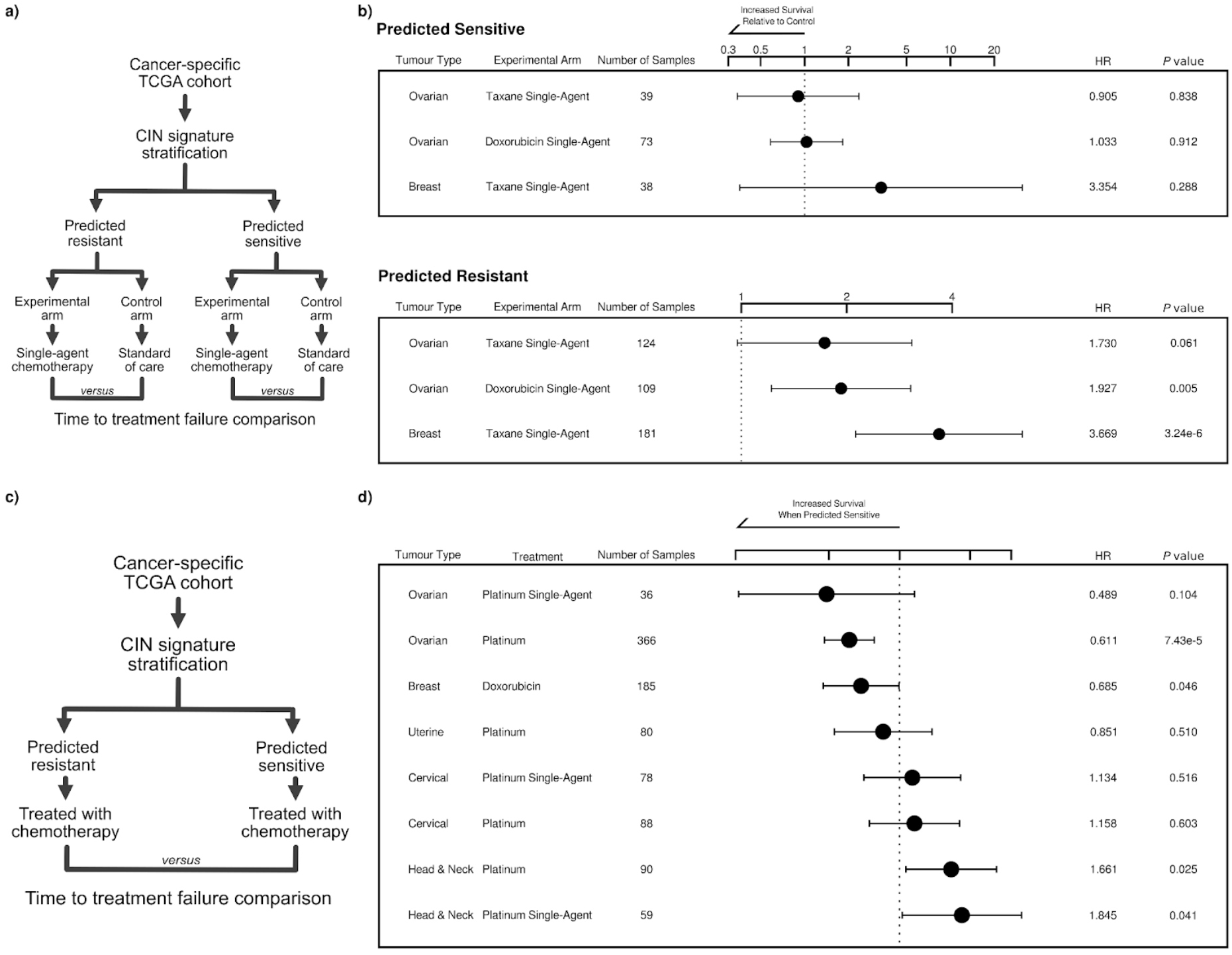
Performance assessment of response prediction in TCGA cohorts using time to treatment failure. **a)** Stratification diagram showing the design of the retrospective randomised control study. **b**) Cox proportional hazards model results for the ovarian and breast TCGA cohorts using the trial design from a). These models account for age at diagnosis, tumour stage and, in the case of breast cancer, the cancer subtype. **c**) Stratification diagram showing the design of the chemotherapy-only survival analysis. **d**) Cox proportional hazards model results for the TCGA cohorts using the trial design from c). Where possible, the models account for age at diagnosis, tumour stage, and cancer subtype.

Within the predicted sensitive groups, no difference was observed in TTF between patients receiving single agent treatment with taxanes or doxorubicin compared to standard-of-care treatment (**Figure 3b)**. However, for patients predicted to be resistant, the use of single-agent chemotherapy resulted in significantly shorter TTF compared to the standard-of-care treatment (ovarian taxane: hazard ratio=1.730, 95% CI=0.975-3.069, p=0.061; ovarian doxorubicin: hazard ratio=1.927, 95% Cl=1.220-3.044, p=0.005; breast taxane: hazard ratio=3.669, 95% Cl=2.122-6.342, p=3.24×10^− 6^; **Figure 3b**). This demonstrates the potential clinical utility of these predictors as able to identify patients who will not respond to taxanes or doxorubicin. Unfortunately, as there was no standard-of-care alternative to platinum treatment for ovarian patients first-line, it was not possible to apply the retrospective randomised control study design to assess platinum predictor performance. For this same reason it was not possible to perform this analysis for platinum prediction in the cervical or head and neck cohorts.

To broaden the pan-cancer assessment of our technology, we conducted additional survival analysis comparing TTF between patients predicted as sensitive and resistant to a specific chemotherapy (**Figure 3c)**. In this case, we were able to include additional cancer-specific TCGA cohorts which did not have sufficient sample size for a randomised controlled trial design (**Extended Data Figure 4**). Classification of the ovarian patients treated with platinum according to the CX2/CX3 activities yielded a significant difference in TTF (single-agent platinum: hazard ratio=0.489 95% CI=0.206-1.159, p=0.104; platinum with co-therapies: hazard ratio=0.611, 95% CI=0.478-0.779, p=7.43×10^−5^; **Figure 3d**). We also observed a significant separation in TTF between breast patients predicted as resistant and sensitive to doxorubicin (hazard ratio=0.685, 95% CI=0.473-0.994, p=0.046, **Figure 3d**). However, our clinical classifier failed to predict platinum response for the cervical, head and neck, and uterine TCGA cohorts, possibly due to limited cohort sizes, the use of TTF as a surrogate for PFS, or unaccounted aspects of the biology of these cancer types in the Cox model.

### Predicting response using liquid biopsy

Finally, we assessed the feasibility of applying our classifiers to liquid biopsies collected at the time of diagnosis from patients enrolled in the OV04 cohort. Plasma samples were used to extract cell-free DNA (cfDNA), which underwent low-pass whole genome sequencing. DNA copy number profiles were generated and samples were categorised based on their ctDNA fraction as either low or high. Out of the 41 patients in the cohort, 29 had plasma samples available. Among these samples, 9 (representing 31% of the cohort) were considered as samples with high circulating tumour DNA (ctDNA) fraction **(Figure 4a)**. The samples with high ctDNA fraction were subjected to CIN signature analysis, similar to solid tumours samples. The remaining plasma samples had insufficient overall tumour DNA to assess chromosomal instability using the currently available CIN signature methods.

**Figure 4.**
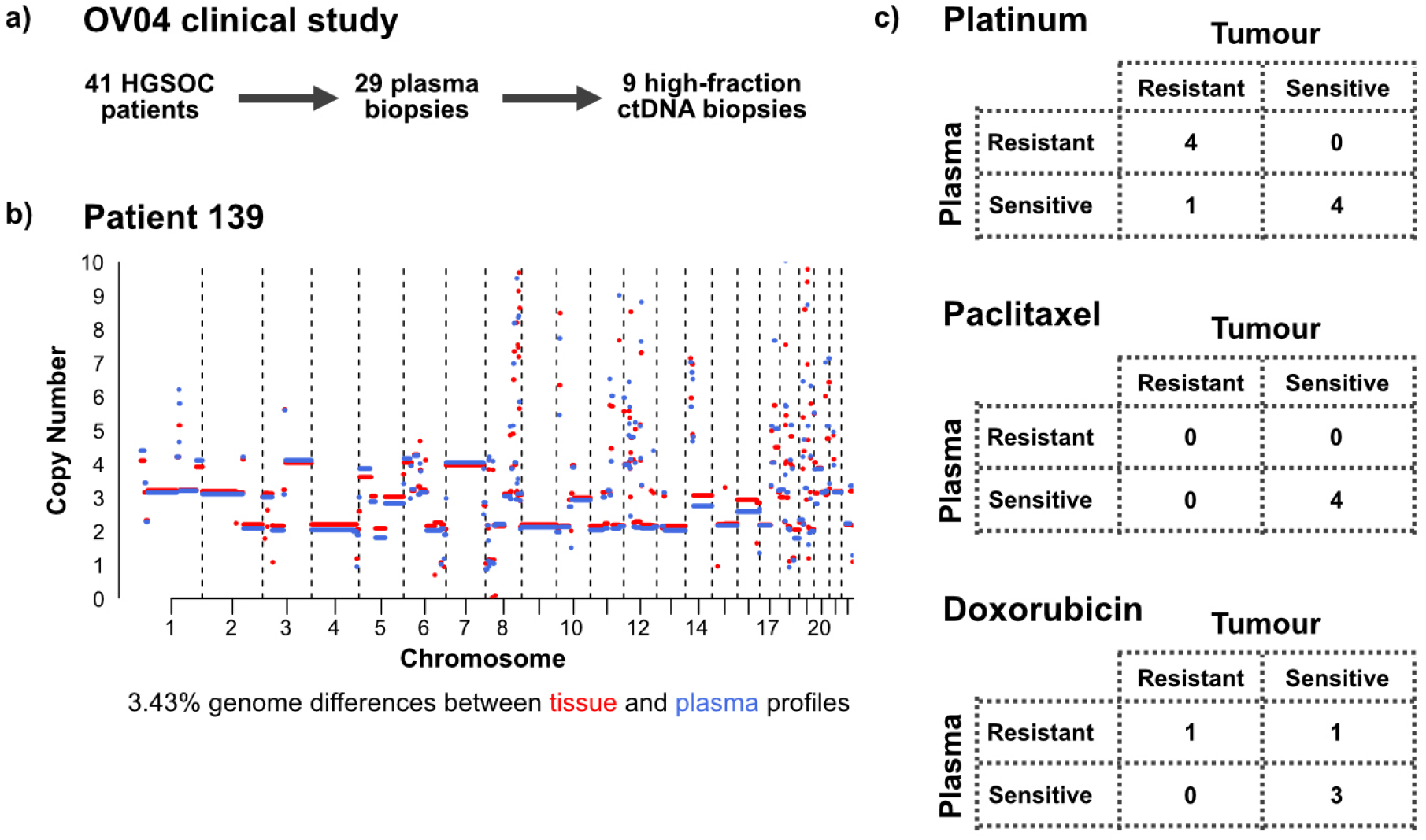
Comparison of response prediction using tissue or liquid biopsy. **a)** Diagram summarising the number of patients with high-quality ctDNA for applying our signature-based predictor. **b**) Matched tumour tissue sample and plasma sample from Patient 139 showing concordant copy number profiles. **c**) Contingency tables showing the number of patients predicted as sensitive or resistant to platinum, paclitaxel and doxorubicin according to their signature activities in tumour tissue and plasma biopsies.

In general, a strong agreement was observed between plasma and tumour tissue pairs in terms of copy number profiles (median percentage of genome-wide copy number differences was 21%; **Figure 4b)**, and activity levels of CIN signatures (median cosine similarity for pan-cancer signatures was 0.88, median cosine similarity for ovarian signatures was 0.81; **Extended Data Figure 5**).

In relation to platinum response prediction, there was agreement between predictions obtained from the tumour tissue sample and the plasma sample in 87.5% (7 out of 8) of the patients (**Figure 4c**). Patient 713 was the only case where the platinum prediction differed between the tissue and blood samples. This patient was predicted to be resistant based on the tumour tissue sample collected at diagnosis, and sensitive based on the plasma sample collected at diagnosis. By using the plasma prediction instead of the tissue prediction, the p-value of the log-rank survival test improved from 0.0187 to 0.0158, while maintaining the same hazard ratio of 0.4. This finding suggests that the plasma-based platinum response prediction might surpass the predictive accuracy of the tumour tissue sample (**Extended Data Figure 6)**.

Among the 17 patients included in the paclitaxel cohort, 23.5% (4 patients) had available high ctDNA fraction samples for comparison. All patients exhibited concordance between the paclitaxel response predictions derived from the tumour tissue and the plasma sample (**Figure 4c)**. Regarding the doxorubicin prediction cohort, 17.2% (5 out of 29 patients) had matching high ctDNA fraction samples available for comparison. Among this subgroup, there was an agreement of 80% (4 out of 5 patients) between the doxorubicin response predictions obtained from the tumour tissue and the plasma sample (Figure 4c). In this case, only patient 788 differed in doxorubicin prediction between the tissue and blood samples. In the tissue sample the patient was predicted to be sensitive, but the blood sample returned a resistant prediction. By using the plasma prediction instead of the tissue prediction, the p-value of the log-rank survival test increased from 0.004 to 0.0143, with the hazard ratio also increasing from 6.901×10^−5^ to 0.013.

These results suggest that for approximately 17-31% of patients, our cytotoxic chemotherapy response predictor may be applied using a simple blood test, without the need for a tumour biopsy or a surgical specimen.

## Discussion

In this study, we tested the potential for CIN signature biomarkers to predict response to multiple chemotherapies including platins, taxanes and anthracyclines.

Biomarker-driven stratification would be a major improvement in the clinic, because these cytotoxic chemotherapies are currently used in multiple cancers as one-size-fits-all treatments. Identifying patients who will not respond to chemotherapy could allow patients to avoid the toxic side effects of treatments that are unlikely to benefit them and physicians could more quickly stratify non-responding patients to other approved therapy choices. In the absence of suitable approved alternatives, this may also have utility for identifying clinical trial patient populations with an unmet medical need. Such an intervention also has the potential to save healthcare payers from unnecessary healthcare costs that do not necessarily result in patient benefit, and in fact several government initiatives have been established to encourage such stratified medicine approaches^23-26^. This current study has also shown that it may be feasible to build such a test on both tumour tissue biopsies as well as liquid biopsies of patient ascites or blood samples, which are more easily accessed in the clinic, potentially reducing future barriers to clinical adoption.

Previous approaches attempting to predict response to cytotoxic chemotherapy have largely focussed on platins and use cell culture or gene expression assays^27,28^. However, these tests have failed to reach widespread adoption in the clinic. The OncotypeDX recurrence score is the only test to have been widely adopted, however is not a direct test for chemotherapy response, rather it determines whether a patient will be adequately managed with hormone therapies alone (compared to a combination with chemotherapy)^29^. The ability to predict response to multiple chemotherapies from a single genomic assay is a novel offering. Furthermore, as genomic-based assays are already widely used in the clinic, there is an increased probability of adoption in the future.

Our analysis was conducted on patient tumours that were collected prior to the approval of PARP inhibitors as a maintenance therapy, which has since been implemented in the clinic for ovarian cancer. It is currently unclear what effect PARP inhibitor maintenance therapy may have on the prediction of paclitaxel, doxorubicin and platinum response. Within ovarian cancer, future work is needed to increase the sample numbers for our analysis and further validate the robustness of our predictions on clinical trial quality data, which could be obtained as part of a prospective observational trial. This work would also need to assess the potential impact of PARP inhibitor maintenance therapy in predicting response to subsequent second-line therapies.

Our analysis encompassed three common chemotherapies, platins, taxanes and anthracyclines, however there are other chemotherapies which could be considered. In order to transition this work into patient benefit, further work is necessary to understand the regulatory pathway of a clinical decision support test for already approved chemotherapies. One of the main challenges that will need to be overcome is the heterogeneity of genomic testing in a clinical environment. Currently, different hospital systems employ different genomic assays including gene panel sequencing and whole genome sequencing. Thus, the use of CIN signature biomarkers for chemotherapy response prediction will need to be enabled across a variety of sequencing technologies.

This study introduces novel genomic analysis techniques for patient stratification for multiple medicines that were not originally developed as targeted therapies. Our study demonstrates the potential to extract predictive insights from both tumour tissue samples and ctDNA samples, which may increase accessibility and utility in a clinical environment. CIN signature analysis can be applied widely across cancer types^4^ and thus our results have the potential for broad future implications for patient stratification and precision medicine in cancer.

## Data availability

Raw data will be made available at time of publication via the European Genome Phenome Archive. Information on the data sources used can be found in **Supplementary Table 5**. All data required for reproducing these analyses has been deposited in figshare (the link can be found in the code repository).

## Code availability

The source code for reproducing analyses and figures will be available at publication. Information on the R packages used in our analysis can be found in **Supplementary Table 4**.

## Acknowledgements

We would like to acknowledge the support of Tailor Bio, Illumina Accelerator, The University of Cambridge, Cambridge University Hospitals NHS foundation trust and Cancer Research UK. Parts of this work were funded by CRUK core grant C14303/A17197, A19274 (FM lab) and the UK Research and Innovation’s (UKRI) Innovate UK Data to Early Diagnosis challenge. J.ST, B.H., M.E., A.FS and G.M. are hosted by the Centro Nacional de Investigaciones Oncológicas (CNIO), which is supported by the Instituto de Salud Carlos III and recognized as a ‘Severo Ochoa’ Centre of Excellence (ref. CEX2019-000891-S) by the Spanish Ministry of Science and Innovation (MCIN/AEI/10.13039/501100011033). J.T., M.E., B.H., A.FS, and G.M. were also supported by a Spanish Ministry of Science and Innovation grant PID2019-111356RA-I00 (MCIN/AEI/ 10.13039/501100011033). A.F.-S. and J.ST received the support of a fellowship from La Caixa Foundation (ID 100010434; LCF/BQ/DR21/11880009 and LCF/BQ/DI22/11940038, respectively). B.H’s postdoctoral research contract was supported by philanthropists via the ‘Amigos/as del CNIO’ Programme.

## Competing interests

G.M., A.P. J.B., J.Y. and F.M are co-founders, directors and shareholders of Tailor Bio Ltd. J.ST and L.M are shareholders of Tailor Bio Ltd. G.M., F.M., A.P. and J.D.B are inventors on a patent on using copy number signatures to predict response to doxorubicin treatment in ovarian cancer (PCT/EP2021/065058). G.M., B.H. and F.M. are inventors on a patent on a method for identifying pan-cancer copy number signatures (PCT/EP2022/077473).

## Extended Data Figures

**Extended Data Figure 1.**
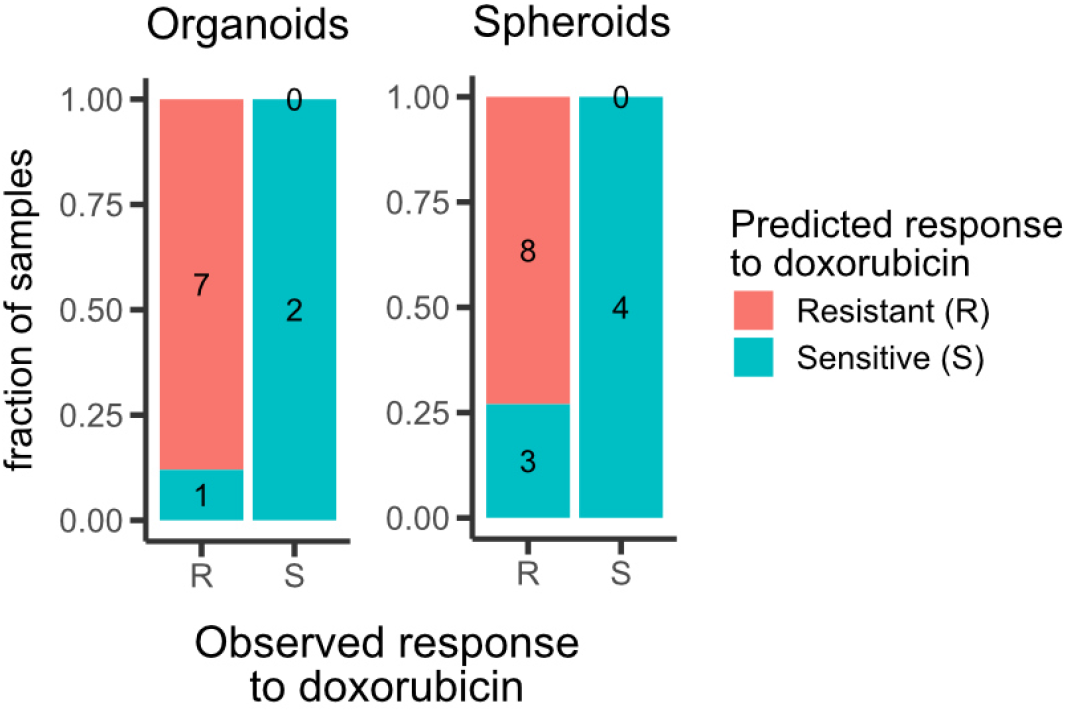
Prefoemance assessment of doxorubicin reponse prediction by using pan-cancer CIN signatures in organoids and spheroids. Bar plots showing the agreement between the observed and the predicted response of organoids and spheroids to doxorubicin. Samples with CX8/CX9/CX13 activities higher than 0.01 were classified as resistant.

**Extended Data Figure 2.**
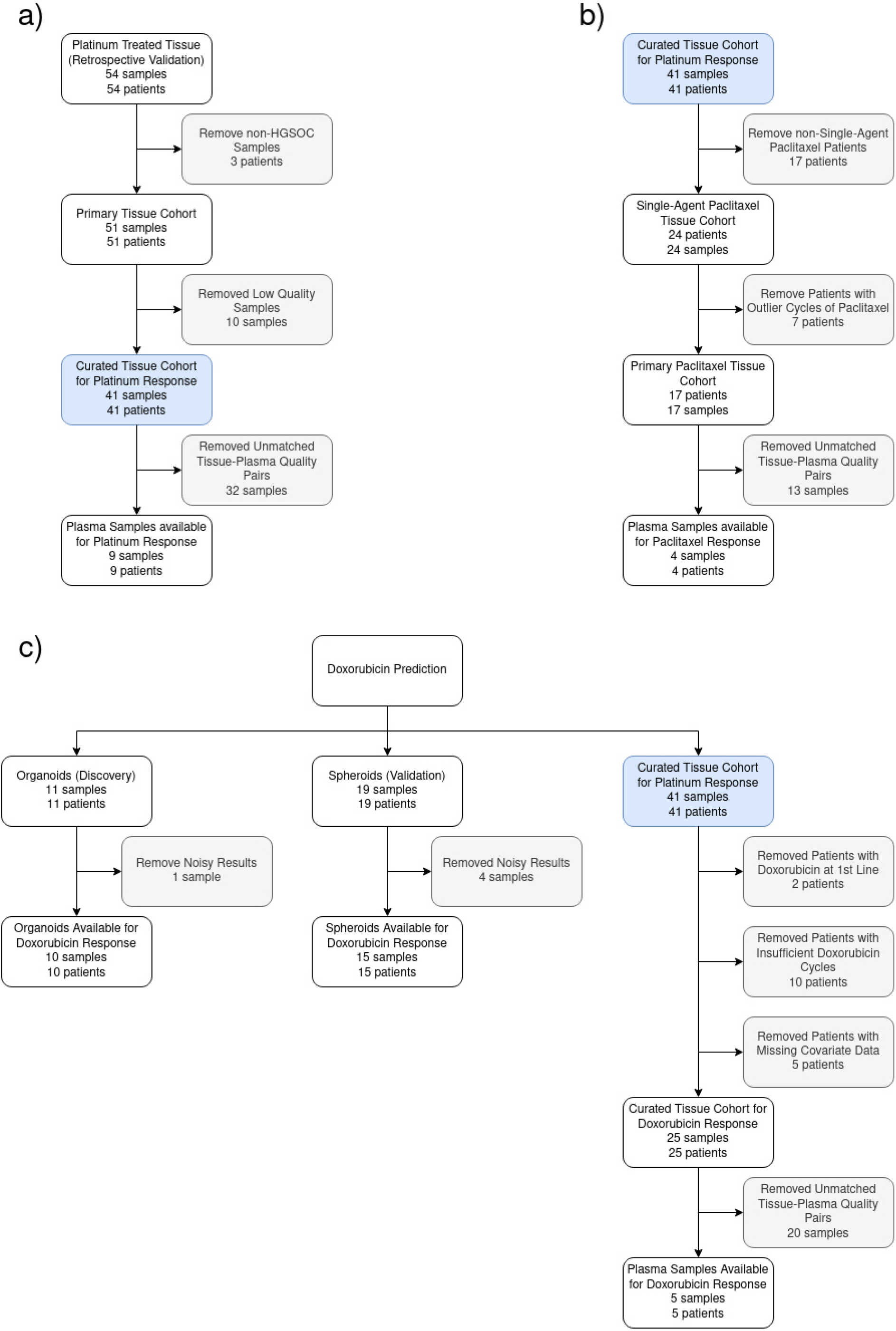
REMARK diagram. **a)** Flow diagram summarising the quality control filtering of samples and patients to get the curated cohort for the retrospective validation of platinum response prediction. **b**) Flow diagram detailing samples and patients filtered to get the curated cohort for the retrospective validation of paclitaxel response prediction. **c**) Flow diagram detailing organoids and spheroids filtered for the phases of biomarker discovery and validation for doxorubicin response, as well as samples and patients filtered for the retrospective doxorubicin response prediction cohort.

**Extended Data Figure 3.**
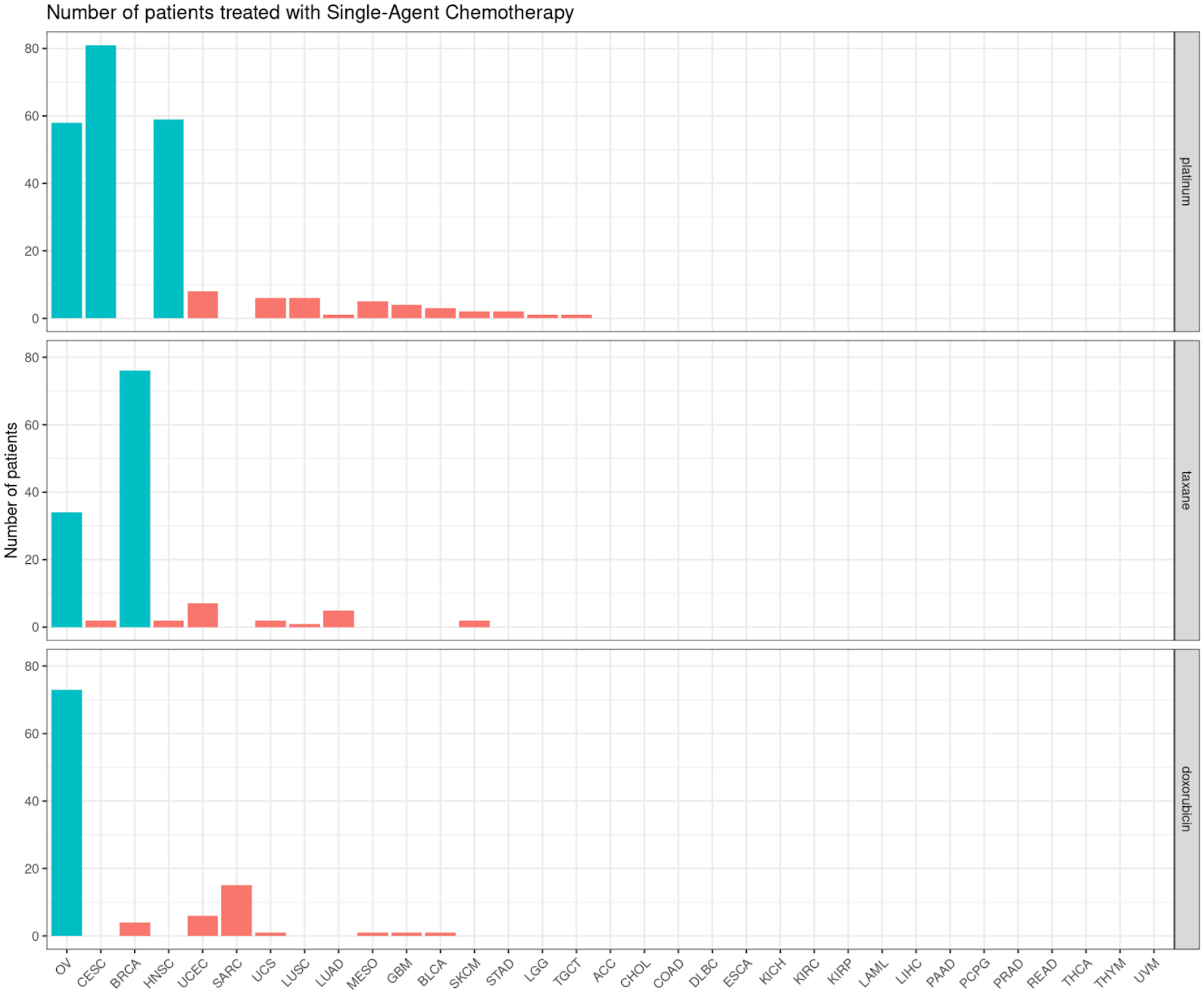
Number of available patients treated with single-agent platinum, taxanes and doxorubicin in each cancer-specific TCGA cohort. Bar plots are coloured according to the cutoff threshold, with blue indicating that a cohort has at least 30 patients available. Patients included in this plot have both treatment response data available and detectable chromosomal instability (>20 CNAs).

**Extended Data Figure 4.**
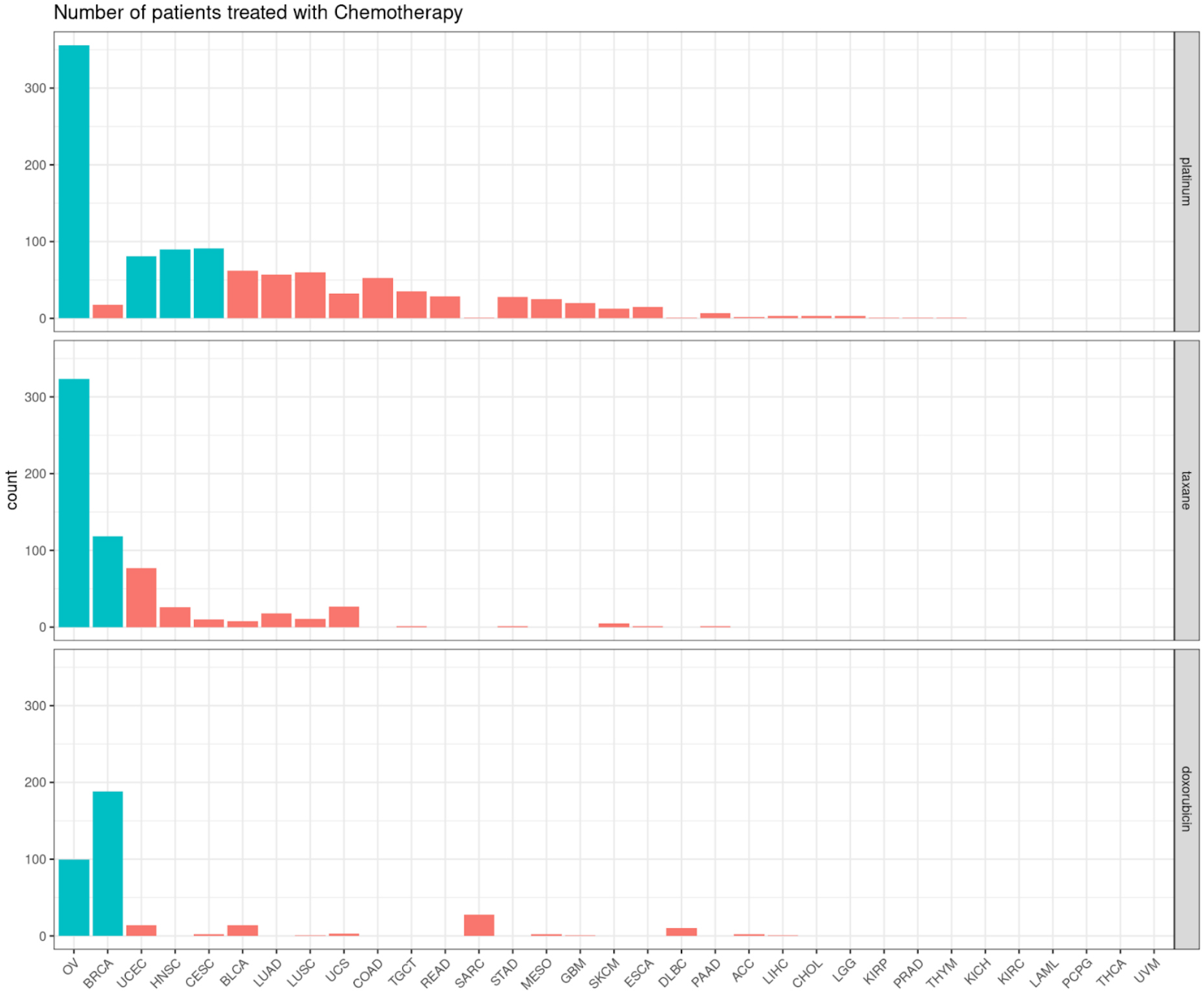
Number of available patients treated with platinum, taxanes and doxorubicin in each cancer-specific TCGA cohort. Bar plots are coloured according to the cutoff threshold, with blue indicating that a cohort has at least 80 patients available. Chemotherapy could be administered as a single-agent or in combination with other therapies. Patients included in this plot have both treatment response data available and detectable chromosomal instability (>20 CNAs).

**Extended Data Figure 5.**
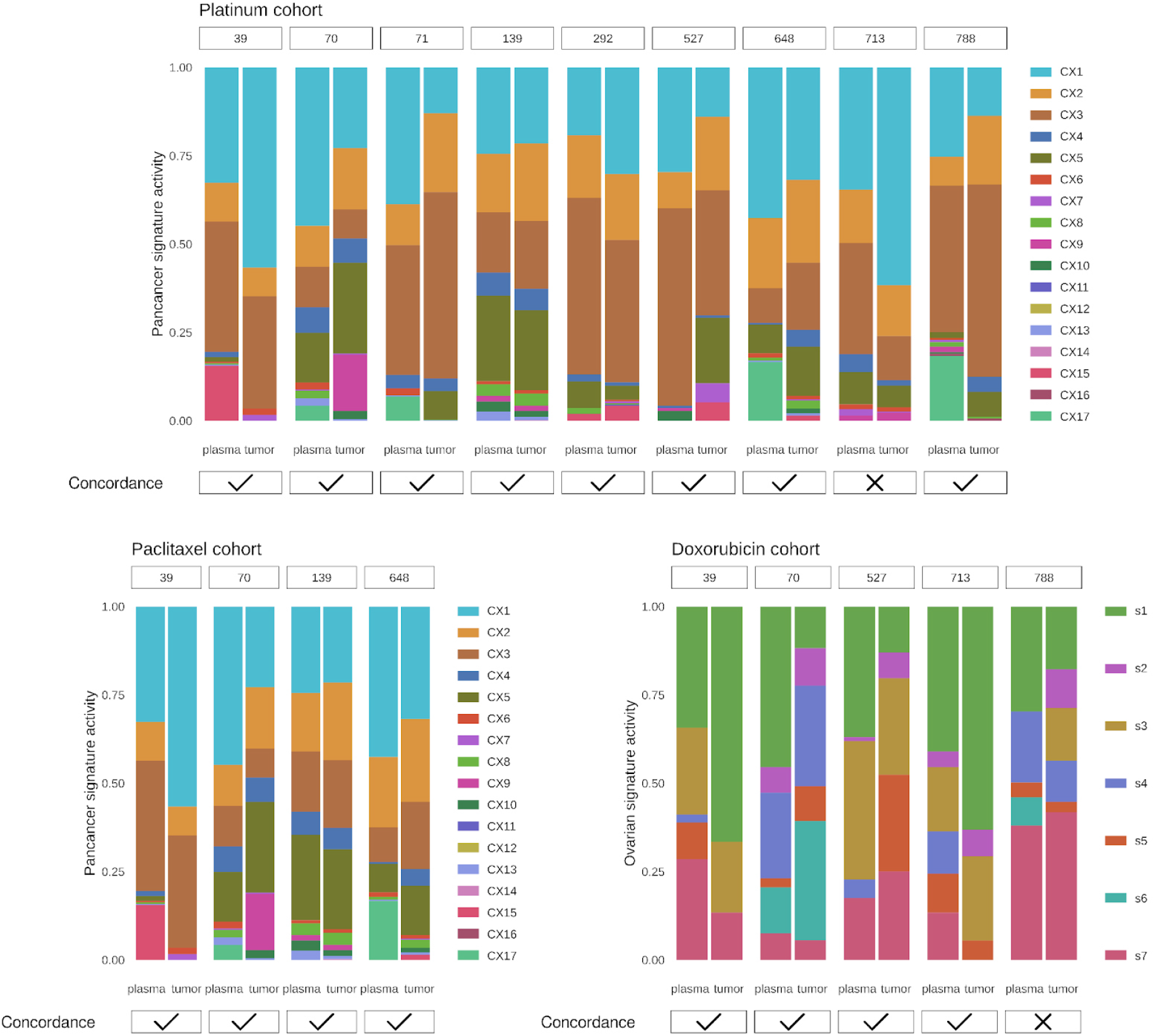
Comparison of signature-based response prediction using tissue or liquid biopsy. Bar plots show activity levels of pan-cancer and ovarian CIN signatures quantified in tissue and plasma samples from matching patients. CIN signatures were applied for predicting response to platinum, paclitaxel and doxorubicin.

**Extended Data Figure 6.**
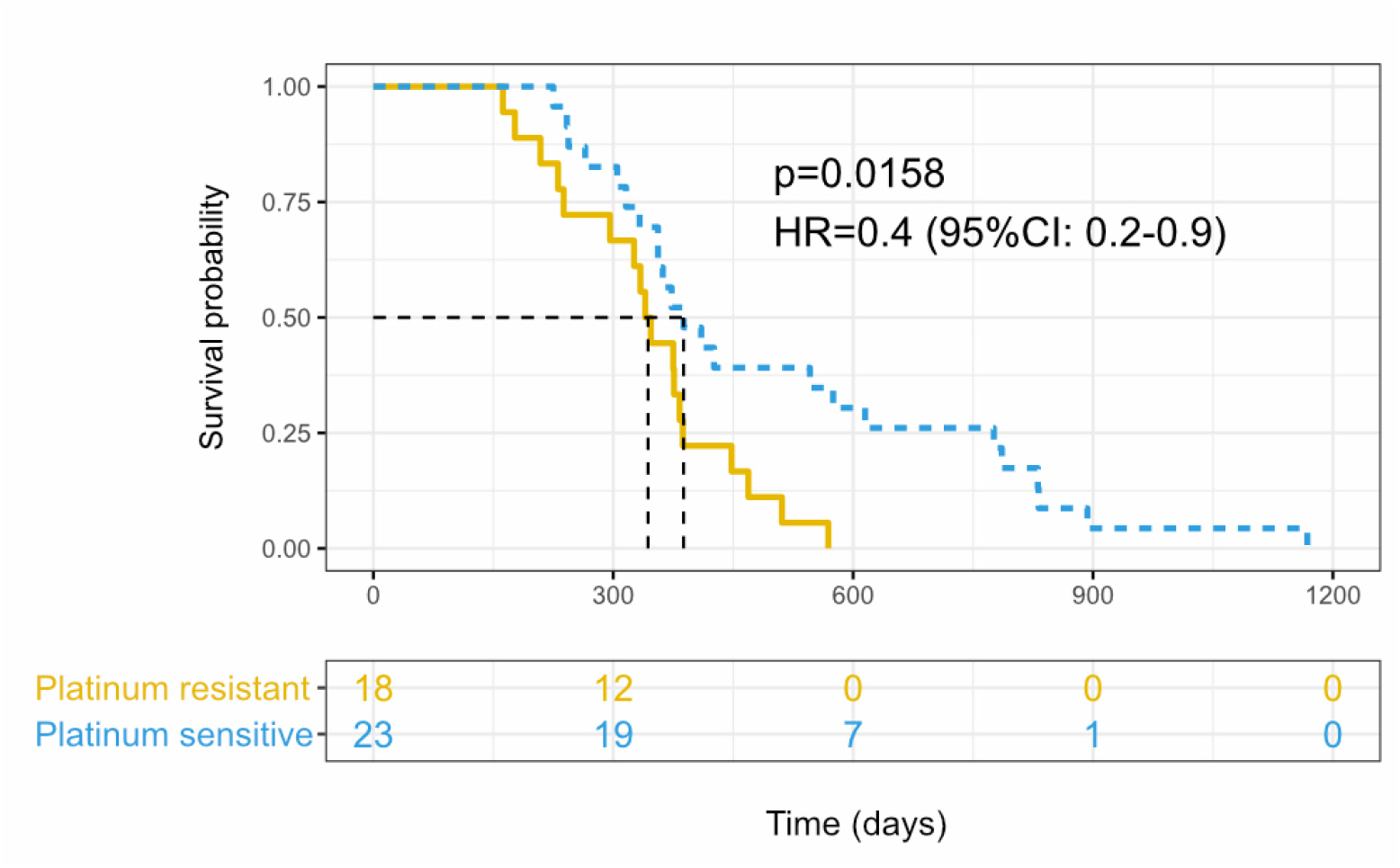
Performance assessment of platinum-based chemotherapy response prediction from tumour tissue samples using plasma prediction labels and progression-free survival across 41 ovarian cancer patients. Kaplan Meier curves for patients predicted to be sensitive (blue) and resistant (yellow). Survival analysis was not corrected by covariates. P-values estimated by the log-rank test.

## Notes

### Summary of Updates

The study has been expanded to include analysis of the mechanism of resistance to doxorubicin treatment and a retrospective randomised control study assessment of predictive performance of biomarkers predicting response to anthracycline and taxane treatment.

