## Supplementary material for "Predicting response to cytotoxic chemotherapy": Methods

Sample Cohorts 2
- Ovarian cancer cell lines 2
- High-grade serous ovarian cancer human samples 2
- The Cancer Genome Atlas (TCGA) collection 3

Sample processing 3
- Tissue samples 3
- Plasma samples 4
- Organoids 4
- Spheroids 4
- Cell lines 4

Doxorubicin treatment assays 5
- Treatment of cell lines 5
- Treatment of spheroids and organoids 5

Micronuclei counting 6

Estimating expected number of micronuclei 6

Shallow whole genome sequencing 7
- Tissue samples, organoids, spheroids and cell lines 7
- Plasma samples 7

Read alignment 7

Absolute copy-number fitting 7

Sample quality control and filtering 8
- Organoids and spheroids 8
- Tissue 8
- Plasma 9

Copy number signature computation 9
- Ovarian signatures 9
- Pan-cancer signatures 9
- Signature-specific thresholding 10

Biomarker discovery for doxorubicin response 10
- Predictive performance assessment in vitro 11

Gene enrichment analysis 11

Signature-based clinical classifiers 11
- Clinical classifier for platinum response prediction 11
- Clinical classifier for paclitaxel response prediction 11
- Clinical classifier for doxorubicin response prediction 12
  - Ovarian signatures 12
  - Pancancer signatures 12

Survival period calculation 13
- Progression-free survival (PFS) 13
  - PFS for platinum treatment 13
  - PFS for doxorubicin treatment 13
  - PFS for paclitaxel treatment 14

Time to Treatment Failure (TTF) 14
Survival analysis 14
Retrospective randomised control study 14
Retrospective study 15
Annotation of breast tumour subtypes 16
Concordance between tissue and plasma pairs 16
References 16

### Sample Cohorts

#### Ovarian cancer cell lines

Four ovarian cancer cell lines CIOV1, CIOV2, CIOV4, CIOV6[1] were derived in-house, OVCAR3 was obtained from the American Type Culture Collection, and PEO23 was a kind gift from Langdon lab.

#### High-grade serous ovarian cancer human samples

Clinical data and samples for patients with high-grade serous ovarian cancer (HGSOC) were collected as part of the prospective Cambridge Translational Cancer Research Ovarian Study 04 (CTCROV04) approved by the Institutional Ethics Committee (REC07/Q0106/63). Patients provided written, informed consent for participation in this study and for the use of their donated tissue for the laboratory studies carried out in this work. All sample cohorts in this analysis were obtained from patients in this study, including primary ascites spheroids, tumour tissue, plasma samples and tumour tissue derived organoids and cell lines.

OV04 is an ongoing observational study that records patient clinical data and collects patient material for the purpose of biomarker and scientific discovery. To compile a real-world cohort for this study, we assessed patients enrolled in OV04 from 2010-2019 and identified 54 patients with available tumour samples who received platinum-based chemotherapy as a first-line therapy and/or doxorubicin and platinum as a subsequent treatment following approved standards of care. These patients underwent histopathological review, with 3 patients discarded for not showing signs of HGSOC pathology. Samples collected from these patients totaled 51 tumour samples and 29 cell free DNA (cfDNA) samples extracted from plasma. Following sample processing and quality control, a final cohort of 41 ovarian cancer patients was identified as suitable for retrospective assessment of treatment response prediction. Among them, 9 patients had high-quality plasma samples to assess prediction performance in liquid biopsies (see **Extended Data Figure 2a** for sample workflow, **Supplementary Table 1** for clinical characteristics of the cohort, and **Supplementary Figures 1-41** for treatment history of each patient).

Apart from these 41 patients, additional patient samples from the OV04 study were used to harvest spheroids from ascitic fluid and derive organoids for the purpose of CIN signature biomarker discovery for doxorubicin prediction. A total of 10 organoids and 15 spheroids were available for downstream analyses (**Extended Data Figure 2b**). Further characterisation of these collections of patient-derived spheroids and organoids can be found in parallel studies[1,2].

For the paclitaxel response subset of this cohort, the 41 platinum-treated ovarian patients were filtered for patients that had been treated with single-agent paclitaxel after 1st line treatment. Patients were then further filtered to remove paclitaxel treatment lines with abnormally high (>25) or abnormally low (<3) cycles of paclitaxel. This resulted in a group of 17 patients, of which 4 also had plasma samples with a sufficiently high ctDNA fraction (**Extended Data Figure 2b**).

For the retrospective validation of our doxorubicin response biomarker, the clinical histories of the 41 patient platinum treated cohort were further inspected to identify those also given doxorubicin as a subsequent treatment. Patients with 3 or fewer cycles of doxorubicin were excluded as they had not received sufficient treatment to reliably impact clinical outcomes. Additionally, patients who received doxorubicin in combination with platinum-based chemotherapy at first-line were removed, resulting in 29 out of the 41 patients suitable for the retrospective validation of doxorubicin prediction. Among them, plasma samples with high ctDNA fraction were available for 5 patients (**Extended Data Figure 2b**).

#### The Cancer Genome Atlas (TCGA) collection

In our previous work, we evaluated the extent, diversity and origin of chromosomal instability across 6,335 high-quality tumours with detectable CIN (>20 copy number alterations) representing 33 cancer types from the TCGA collection[3]. Here, we used CIN signature activities quantified in our previous work to predict response to chemotherapies. To do so, we downloaded clinical data from TCGA cohorts from the data portal of the Genomic Data Commons (GDC). Curated clinical data was available for the HGSOC subset of the TCGA ovarian cohort[4].

### Sample processing

#### Tissue samples

Formalin-fixed, paraffin-embedded (FFPE) tissue blocks were cut as 8μm sections and tumour-enriched regions were recovered by macrodissection based on regions marked on an adjacent haematoxylin-and-eosin-stained section by the study pathologist. DNA was extracted from 3-10 sections at 8μm thickness using QIAmp DNA Micro kit (Qiagen) with the following modification to the original protocol: an additional incubation step with Buffer ATL at 95 ◦C for 15 minutes was introduced before adding proteinase K. The paraffin was removed using a xylene/ethanol method. DNA extraction from fresh frozen tumour tissues and spheroids fraction was performed using Allprep DNA/RNA tissue kit (Qiagen) following manufacturer's instructions.

#### Plasma samples

We focused on selected plasma timepoints collected before the primary line of chemotherapy treatment and before doxorubicin treatment (usually before 2nd or 3rd line of therapy). DNA was extracted from 2 or 4 mL of plasma using the QIAamp circulating nucleic acid kit (Qiagen) or QIAsymphony (Qiagen) according to the manufacturer's instructions.

#### Organoids

Samples were obtained from patients via surgical resection, ward drains or surgical washings. Solid tumours were assessed by a pathologist and only tumour samples with ≥ 50% tumour cellularity were selected for organoid model derivation. Organoids were derived as previously described[1].

Organoid culture medium was refreshed every 2 days. To passage the organoids, the domes were scraped and collected in a falcon tube, TrypLE (Invitrogen) was added and they were incubated at 37 °C for approximately 10 min. The suspension was centrifuged at 800g for 2 min and the cell pellet was resuspended in 7.5 mg/ml BME-2 supplemented with complete media and plated as 20 µl droplets in a 6-well plate. After allowing the BME-2 to polymerize, complete media was added and cells incubated at 37 °C. DNA was extracted from cell pellets using the Qiagen Allprep DNA/RNA extraction kit according to manufacturer instructions.

#### Spheroids

Ascitic fluid was collected from patients between 100ml-2L volume. The fluid was initially gently centrifuged at 800xg for 5 minutes and the majority of the supernatant was removed. The sample was filtered using autoclaved muslin cloth, the flow trough was then filtered again using a cell strainer at 40µm. Spheroids from the strainer were then recovered by a 10ml wash with PBS and centrifuged at 1500rpm for 5 min. The spheroid fraction was divided in two: a cell pellet for DNA extraction and resuspension of cells in filtered acellular ascitic supernatant and 8% DMSO for the drug screen. Spheroids were thawed and put in media overnight to fully recover before dispensing for the drug screen. DNA extraction was performed using the Qiagen Allprep DNA/RNA extraction kit according to manufacturer instructions.

#### Cell lines

All cell lines were maintained in DMEM/F12 or RPMI1640 plus 10% of foetal calf serum (FCS). Cell line identities were confirmed by STR profiling. Cells were regularly screened for mycoplasma using a MycoAlert Mycoplasma Detection Kit (Lonza). Sulforhodamine B (SRB) colorimetric assay was used for quantifying cell numbers and cell proliferation in culture. DNA extraction from cell pellets of approximately 2x10^6 million cells per sample was performed using Qiagen Allprep DNA/RNA extraction kit following manufacturer recommendations.

### Doxorubicin treatment assays

#### Treatment of cell lines

Cancer cell lines were initially exposed to four doxorubicin concentrations (0, 0.1, 0.5 and 1μM) for 5 days. Since cell growth rates significantly decayed after one day of exposure to all concentrations tested (**Supplementary Figure 42**), we then performed cell viability assays by treating cells with 0.025μM and 0.05μM for 48 hours. Two cell lines showed 100% confluence after 48 hours of being exposed to low doses of doxorubicin and therefore were discarded from further analyses due to signal saturation (**Supplementary Figure 43**). All the remaining cell lines (OVCAR3, PEO23, CIOV2 and CIOV4) showed stable growth rates in the presence of 0.025μM of doxorubicin, and were therefore used for evaluating micronuclei formation and tolerance. For doxorubicin treatment and micronuclei formation experiments, optical 94-well plates (CellCarrier-96 Ultra, Perkin Elmer) were coated with sterile Poly-L-Lysine (P4832, Sigma), and cells were seeded using previously estimated seeding densities until 50% confluency was reached. Subsequently, cells were treated with 0.025, 0.05 and 0.1μM of doxorubicin or control for 5 days. All treatment experiments were performed in triplicates.

#### Treatment of spheroids and organoids

For organoid and spheroid samples doxorubicin treatment was administered following the method in Martins *et al*[2]. Briefly, an 8-point half-log dilution series of doxorubicin starting at 30μM was dispensed into 384 well plates using an Echo® 550 acoustic liquid handler instrument (Labcyte) and kept at -20°C until used. Organoid plates were spun down and 50 μl of suspension added per well using a Multidrop™ Combi Reagent Dispenser (Thermo-Fisher). Following 5 days of drug incubation cell viability was assayed using 30μl of CellTiter-Glo® (Promega). Screens were performed in technical triplicate. An untreated control was used to normalise response values to be equivalent to the percentage of viable cells remaining. To assist with dose response curve fitting, dummy values were added for each sample below and above the minimum and maximum dose ranges at 100% viable cells with a dose of 1e-3μM and 0% viable cells at a dose of 300μM. A 4 parameter log-logistic model was used to fit dose response curves, which includes the IC50 parameter used here. Fitting was performed using the *drm* function in the *drc* package in R[5]. For spheroid samples, the IC50 values were scaled by the inverse of the tumour purity to account for increased cell viability due to normal cell contamination.

We then determined an IC50 threshold which divided the samples into those considered sensitive or resistant based on the clinical characteristics of the patients. As our *in vitro* drug screen is analogous to patients being treated with doxorubicin as a monotherapy following first line treatment with platinum based chemotherapy, we estimated the expected number of sensitive samples based on response observed in clinical trials. Patients resistant to platinum chemotherapy are expected to have an 18% response rate to doxorubicin monotherapy[6–12], and sensitive patients a 28% response rate[13]. Patients who had relapsed disease less than 6 months after first-line platinum based chemotherapy were considered resistant and greater than 6 months sensitive (**Supplementary Table 3**). These data allowed us to estimate the expected number of doxorubicin sensitive organoids to be approximately

2 (5 x 0.28 + 5 x 0.18 = 2.3) and the sensitive spheroids to be 4 (8 x 0.28 + 7 x 0.18 = 3.5). Samples were ranked based on their IC50 and a threshold set yielding the expected number of sensitive samples in each case.

### Micronuclei counting

Micronuclei counts were estimated using fluorescent imaging. In brief, following doxorubicin treatment, cells were fixed in 100% ice-cold methanol for 5 mins at -20°C, washed three times with 1xPBS and permeabilised with 1% TritonX100 + 0.5% NP40 in 1xPBS for 5 mins at room temperature, before blocking for 1 hour with 5% BSA in 1xPBS at room temperature. To allow reliable micronuclei identification at single cell level, cells were stained with Hoechst (1μg/ml), phospho-histo H3 (pHH3; 1μg/ml), and cytokeratin 7 conjugated to fluorophore 488 (1μg/ml). Primary non-conjugated antibodies were diluted in blocking buffer, added to the cell lines and incubated overnight at 4°C. Cells were washed three times with 0.1% Tween 20 in PBS, and secondary antibody (Alexa Fluor 555, 1μg/ml) diluted in blocking buffer was added to the cells for 1 hour at 37°C. Hoechst 33342 was diluted to 1 μg/ml in MilliQ water, added to the cells and incubated for 15-30 mins at room temperature. Cells were washed and stored in PBS.

Stained cell lines were imaged in filtered 1xPBS as mounting medium (200 μl/well) with a confocal 40x 1.1NA water objective using the Operetta CLS™ high-content analysis system. Images from ten independent non-overlapping imaging fields were acquired from each well (three wells per cell line and treatment group), with Z-stack images being collected at a step size of 0.5 μm across all imaging channels.

Image analysis was performed using the Harmony 4.9 software. Images were reconstructed as maximum projections using basic brightfield correction, and analysed by performing nuclei segmentation, removing border object to only include whole cells, identifying cytoplasm and regions of interest using cytokeratin stains and performing spot counting and micronuclei identification using the Harmony 4.9 Micronucleus Analysis RMS module. Mitotic cells identified via pHH3 staining were excluded from micronuclei counting analysis as condensed chromosomes were commonly misidentified as micronuclei. In addition pHH3 stains were used to estimate the mitotic index of the treated and stained cell lines.
The fraction of cells with micronuclei in cell lines after being exposed to doxorubicin was computed by taking into account the number of mitotic cells which have doubled their genetic material. pHH3 score was used as a marker for cells undergoing mitosis. We then compared the observed and the expected fraction of micronucleated cells in each cell line in order to evaluate if the cell lines with ovarian signature 6 have less fraction of micronucleated cells than expected as a sign of doxorubicin resistance.

### Estimating expected number of micronuclei

To perform statistical analyses, we then modelled the expected fraction of cells with micronuclei across divisions in culture under the presence or absence of doxorubicin. The model considered the fraction of dividing cells in each cell line after 48 hours in culture with or without 0.025μM of doxorubicin ($f$), the fraction of induced micronuclei persisting across cell divisions ($p$=75%[14]), and the fraction of cells hitted by doxorubicin ($d$=60%[15]) or experiencing damage under normal culture conditions ($d$=25%[15]).

This approach for estimating the fraction of micronucleated cells was based on a previous study that monitored micronucleus-containing cells through time-lapse imaging[14]. This study observed an increase in the number of micronuclei per cell after the first division, indicating that some cells contained more than one micronuclei, ideally ensuring that each daughter cell possessed at least one micronucleus. However, subsequent generations exhibited a decrease in the number of micronuclei per cell, suggesting that some cells lost or reintegrated their micronuclei. This phenomenon of micronuclei persistence occurred in approximately 75% of cases.

Furthermore, we also considered cell viability in the model. Micronuclei persisted over time and generations in non-dividing cells, while the number of micronucleated cells was expected to reduce by 50% with each cell division. In the first division, where there was no degradation, reincorporation, or extrusion of micronuclei, the estimated rate of cells taking up doxorubicin (and consequently being damaged by the drug) was approximately 60%[15]. Under control conditions, the rate of cells accumulating damage was approximately 25%[15]. Consequently, the formation of micronuclei would only be induced in this fraction of cells.

Taking altogether, we estimated the number of micronucleated cells after the first division ($MN_t$) as follows:

$$MNt = (1 - f) \times MNi + d \times f \times \left(\frac{MNi}{2} + MNi + \frac{1-MNi}{2}\right)$$

where $MN_i$ is the number of micronuclei observed at time 0 in culture. In case the fraction of dividing cells ($f$) was higher than 1, we corrected the number of micronucleated cells across subsequent divisions as follows:

$$MNt = (1 - f) \times MNt + p \times f \times \frac{MNt}{2}$$

The fraction of dividing cells was corrected across divisions by subtracting 1.

### Shallow whole genome sequencing

#### Tissue samples, organoids, spheroids and cell lines

Whole-genome sequence libraries were prepared from 50ng DNA using Illumina DNA prep (Illumina) and SMARTer Thruplex DNA-Seq (Takara) reagents following the manufacturer's protocol. Library quality and quantity were assessed with D5000 on 4200 Tapestation and Qubit BR dsDNA assay according to the supplier's recommendations. Libraries were then pooled together in equal ratios and sequenced using PE-50 mode on NovaSeq S2 flow cell aiming for 10 million reads per sample.

#### Plasma samples

10ul of extracted circulating nucleic acids was taken as an input for whole genome library preparation using ThruPLEX DNA-Seq (Takara) library prep kit with the following modifications: no DNA shearing was performed, 14 PCR cycles was applied, library purification using Ampure beads (Beckman Coulter) was performed separately for each sample, elution was performed using 20ul of Tris EDTA buffer. Generated libraries were quantified using the Fragment Analyser NGS kit (Agilent Technologies) diluted to 10nmol/l and pooled in the same proportions. All libraries were sequenced using NovaSeq S2 (Illumina) using PE-150bp mode to achieve at least 10 million reads per sample.

### Read alignment

Reads were aligned as single-end against the human genome assembly GRCh37 using BWA-MEM v0.7.17[16], following which duplicate reads were identified and marked using the MarkDuplicates tool in the GATK v4.1.8.1[6] toolsuite.

### Absolute copy-number fitting

After alignment, relative copy number was computed using QDNAseqmod[17] with a bin size of 30kb. Absolute tumour copy number (the number of chromosome copies of each DNA segment in the tumour cells in a sample) was computed for every bin across each sample. Each segmented relative copy number bin estimate *j* was transformed from relative copy number (*rCN)* to absolute copy number (*aCN*) as follows:

$$aCN_j = \frac{1}{purity} \cdot \left(\frac{rCN_j}{d} - 2 \times (1 - purity)\right) - 2$$

where *purity* is the fraction of tumour cells in the sample, and *d* is a constant proportional to the read depth, which is computed from the mean relative copy number of the sample, *r*, and the average absolute copy number of the tumour cells in the sample, *ploidy*:

$$d = \frac{r}{(ploidy \times purity + 2 \times (1 - purity))} .$$

Both *purity* and *ploidy* were unobserved in the data and were estimated using a grid search of purities ranging from [0.05,1] in 0.01 increments and ploidies ranging from [1.8,8] in increments of 0.1, minimising the following mean squared error:

$$e_{purity,ploidy} = \frac{1}{J} \cdot \sum_{j=1}^{J} (aCN - round(aCN))^2$$

Cell lines and organoid samples were assumed 100% pure so purity was fixed to 1 and a search was only performed across ploidy states. Purity/ploidy values were excluded from consideration if they resulted in a fit which showed greater than 10 megabases of the genome with homozygous loss. For tissue samples, an additional filter was used, removing fits which did not show at least one genomic segment at every integer copy number state from 1 to *ploidy*.

### Sample quality control and filtering

A summary of filtering criteria and number of samples retained for further analysis can be found in **Extended Data Figure 2**.

#### Organoids and spheroids

*Drug screen:* Samples showing greater than 20% standard deviation across more than 3 dose concentrations were removed from downstream analysis.

#### Tissue

*Copy number (tissue):* Tissue samples with absolute copy number profiles were evaluated using a manual curation where samples were rated following a 1-3 star system, following the criteria from our previous work[18]. This star rating system is designed to evaluate the copy number fits for their ability to produce reliable CIN signature activities. The description of the different star ratings is as follows:

- 1-star: samples show some observed copy number change, but not enough to provide a reliable CIN signature activity. They can be used in some analysis if the size of the cohort outweighs the lower reliability of the signatures.
- 2-star: samples have clear copy number fits, but can be underpowered, or are too noisy to be 3 star.
- 3-star: samples have ideal copy number fits, minimal noise, and are not underpowered.

Following absolute copy number fitting, samples were rated using this star system. Overall only 10 samples were identified as 1-star and were subsequently excluded from downstream analysis. These 1-star samples showed noisy copy number profiles and were considered likely to have incorrect segments and missing calls. In general, only 2 and 3-star copy number profiles are usable in downstream analysis, resulting in 41 out of 51 primary samples (80.39%) being usable for signature analysis. Overall this is in line with typical sequencing success rates for archival material[19].

#### Plasma

*ctDNA Frequency:* Samples were curated according to their copy number profiles, where samples with low levels of circulating tumour DNA displayed insufficient copy number alterations. These samples were excluded from the analysis as the estimation of the copy number signatures is not accurate. This resulted in 9 out of 29 (31%) being usable for signature analysis.

### Copy number signature computation

#### Ovarian signatures

Copy number features were calculated from the absolute copy number profiles as detailed in Macintyre et al[18]. Briefly, absolute copy number profiles were summarised by calculating, for each sample, the genome-wide distribution of six features associated with copy number (CN) events:

- Segment size: the length of each genome segment;
- Breakpoint count per 10MB: the number of genome breaks appearing in 10MB sliding windows across the genome;
- Change-point copy-number: the absolute difference in CN between adjacent segments across the genome;
- Segment copy-number: the observed absolute copy-number state of each segment;
- Breakpoint count per chromosome arm: the number of breaks occurring per chromosome arm;
- Length of segments with oscillating copy number: a traversal of the genome counting the number of contiguous CN segments alternating between two copy-number states, rounded to the nearest integer copy-number state.

For each sample, a sum of posterior probability vector was calculated using the feature component definitions as outlined in Macintyre *et al.*[18] and the predict function in the *flexmix* package in R[20].

The sum of posterior probability vectors were used to compute signature activities using the *LCD* function found in the *YAPSA*[21] package in R, and the signature definition matrix reported in Macintyre *et al*[18]. The *LCD* function in this package can calculate activities for known signatures, given a predefined signature weight matrix.

#### Pan-cancer signatures

Pan-cancer copy number signatures were calculated in a similar manner to the ovarian copy number features but with the removal of segment copy number as a feature. As with the ovarian signatures, a sum of posterior probability vector was created using the pancancer feature component definitions. This probability vector was then used to calculate the signature activities using linear combination decomposition with the signature definition matrix for the 17 pancancer signatures developed in Drews *et al*[3].

#### Signature-specific thresholding

Signature-specific thresholds were determined for both the pan-cancer and ovarian signatures within the retrospective clinical cohort following methods established in Drews *et al*[3]. A Monte Carlo simulation was performed where noise was added to measurements of the copy number features, after which the difference in the resulting signature activities were investigated. This simulation was run 1000 times over the cohort. Samples were then considered to be adequately similar if cosine similarity between the original and noisy signature activities was higher than 0.85. For those samples where the original activity was zero and the perturbed activity was non-zero, a single Gaussian distribution was fitted to the nonzero activities using the function *Mclust* from the R package *mclust*[22]. The 95th percentile was chosen as the cutoff point, giving an activity level for each signature, below which it is not possible to reliably distinguish between a truly non-zero activity and a zero activity perturbed by sources of noise. For every sample where either pan-cancer and/or ovarian signature activities were computed, the activities were set to 0 if they fell under the signature-specific thresholds.

### Biomarker discovery for doxorubicin response

In our endeavour to use CIN signatures as biomarkers for predicting doxorubicin response, we initially aimed at using genomic and drug response data obtained from the 297 cancer cell lines that were used to identify CX5 as a biomarker for paclitaxel prediction (see section "Clinical classifier for paclitaxel response prediction"). However, we encountered limitations in the range of AUC values (0.58 ± 0.08) for doxorubicin response across the cell lines, which impeded the possibility of achieving accurate correlations. Furthermore, there was a lack of sufficient cell lines exhibiting AUC values close to 0, and therefore resistant to this treatment.

For that reason, we designed *in vitro* experiments to identify and validate a signature-base doxorubicin response predictor. Cancer cell lines, organoid and spheroid samples were treated with doxorubicin *in vitro*, the response was measured using IC50, and the activity of CIN signatures associated with focal amplifications (ovarian signature 6 and/or pan-cancer signatures CX8, CX9 and CX13) was computed from sWGS of the samples prior to treatment. In addition, to identify the biological mechanism driving doxorubicin resistance, we compared the observed and expected fraction of micronucleated cells in culture under doxorubicin treatment (see section "Estimating expected number of micronuclei"); and explored transcriptomic differences between ovarian cancer patients included in the TCGA predicted as resistant and sensitive to doxorubicin according to our signature-based classifier (see section "Gene enrichment analysis").

#### Predictive performance assessment *in vitro*

For the training (organoid) and validation (spheroids) analyses, performance was assessed using specificity and sensitivity for predicting resistance to doxorubicin. Sensitivity was considered as the proportion of sensitive samples correctly identified and specificity as the proportion of resistant samples correctly identified. In order to assess the significance of the observed sensitivity a permutation test was performed. All possible resistant/sensitive sample labels were computed, a predictive threshold determined that would yield 100% specificity given the sensitive sample labels, and then sensitivity recorded. The resulting distribution of all observed sensitivity values was used to determine the probability of observing a sensitivity greater than or equal to the observed sensitivity.

### Gene enrichment analysis

To further explore the mechanism driving micronuclei tolerance in resistant tumours, we performed a differential expression analysis between ovarian TCGA tumours predicted as resistant or sensitive to doxorubicin based on our clinical classifier by using the *DESeq2* package in R[23]. Genes with low overall expression counts (sum counts across all samples < 10) were excluded. Genes that exhibited significant expression differences across the two groups were ranked based on their significance and fold changes to facilitate. For this analysis, we selected "h.all.v7.1.symbols.gmt" as the reference set, the number of permutations was set to 10,000, and a minimum of 15 genes was required for each gene set. Gene sets with an FDR-adjusted p-value below 0.05 were considered for further analyses, and the Normalised Enrichment Score (NES) was used to indicate the level of enrichment within the gene sets.

### Signature-based clinical classifiers

#### Clinical classifier for platinum response prediction

Platinum response prediction was performed using a classifier based on the pan-cancer CIN signatures CX3 and CX2, developed and described in Drews *et al*[3]. For each thresholded signature, the activity levels were centred to the mean and variance activity scaled to allow for comparison between the different signatures. The centred and scaled CX3 and CX2 activity levels were then compared. If CX2 activity was greater, then the sample was predicted to be platinum resistant, by contrast, if CX3 activity was greater, then the sample was predicted to be platinum sensitive.

#### Clinical classifier for paclitaxel response prediction

By correlating signature activities, gene essentiality scores, and sensitivity to drug perturbation, we previously identified CX3 and CX5 signatures were significantly correlated with response to paclitaxel[3]. To build a clinical classifier, a linear model for predicting paclitaxel response including both signatures was initially generated, observing that CX3 and CX5 had an independent effect on the area under dose response curve (AUC) value for paclitaxel. Given that CX5 presented a higher correlation coefficient than CX3, we finally selected CX5 for building a binary classifier to predict patients as sensitive or resistant to paclitaxel (**Supplementary Table 2**).

To apply CX5 signature as biomarker for predicting paclitaxel response in ovarian cancer, we estimated the optimal threshold to maximise prediction performance in the 17 patients enrolled in the OV04 clinical study treated with paclitaxel. This optimal threshold was applied also to classify TCGA-OV patients. Patients were predicted as sensitive if CX5 activity was greater than the optimal threshold, otherwise as resistant.

In the case of the TCGA-BRCA cohort, we did not have a reference cohort with curated clinical data for setting the classification threshold to maximise response prediction.

Therefore, we selected the optimal threshold to classify 30% of TCGA-BRCA patients as sensitive to paclitaxel, which is the expected response rate reported in the literature[24].

### Clinical classifier for doxorubicin response prediction

#### Ovarian signatures

Doxorubicin response prediction was performed using the ovarian signature 6, developed and described in Macintyre *et al*[18]. Signature 6 has previously been implicated in mutational processes representing focal amplification as a result of the formation of micronuclei[18]. A binary classifier was applied to the thresholded signature 6. Samples with 0 signature 6 activity were considered doxorubicin sensitive, and samples with non-zero activity were predicted to be doxorubicin resistant.

The process of selecting signature 6 as the optimal biomarker for doxorubicin response prediction, along with the optimal threshold, involved genomic and drug response data obtained from organoids (for training) and spheroid samples (for testing). Briefly, we determined a predictive threshold that achieved 100% specificity when considering the sensitive sample labels in the training cohort. Subsequently, we applied this selected threshold to the spheroid data to verify that it effectively distinguished between resistant and sensitive samples. We conducted this analysis for all seven ovarian signatures and concluded that ovarian signature 6 captures the highest prediction signal.

#### Pancancer signatures

Ovarian signatures were specifically tailored for dissecting CIN in ovarian cancer samples sequenced via sWGS[18]. For that reason, we opted to employ pan-cancer signatures that represented focal amplifications associated with extrachromosomal DNA (ecDNA) (CX8, CX9, and CX13) as biomarkers for predicting the response to doxorubicin in the TCGA cancer-specific cohorts.

In order to identify the most effective threshold for optimising the prediction of doxorubicin response, we used drug response data obtained from organoids and spheroids. Using the organoids data, we conducted a grid search, varying the activities from 0.005 to 0.015 in increments of 0.001, exploring all possible combinations. We then selected combinations that achieved both 100% specificity and the highest sensitivity. Subsequently, we applied these selected activity combinations to the spheroids data to determine the optimal combination for predicting doxorubicin response. Samples with an activity level greater than 0.01 in CX8, CX9, or CX13 were classified as doxorubicin resistant, while samples falling below this threshold were predicted to be sensitive to doxorubicin.

### Survival period calculation

#### Progression-free survival (PFS)

##### PFS for platinum treatment

PFS was calculated following the CA125 definitions of progression in 1st line therapy agreed by the Gynecologic Cancer InterGroup (GCIG) in November 2005 (https://gcigtrials.org/system/files/CA%20125%20Definitions%20Agreed%20to%20by%20GCIG%20-%20November%202005.pdf).

Patients were sorted into three separate categories based on their CA125 levels over the course of their platinum 1st line treatment; Category A patients began with 'abnormal' readings pre-treatment, that then fell into the normal range during treatment; Category B patients began with abnormal readings that never normalised; and Category C patients began in the normal range. In these definitions, the normal range was chosen to be 0-35 CA125, and the abnormal range 35+, following NICE guidance (https://www.nice.org.uk/guidance/cg122/resources/ovarian-cancer-recognition-and-initial-management-pdf-35109446543557).

These categories then defined the patient-specific CA125 progression threshold. In categories A and C, progression occurred at the 1st CA125 reading that was at least twice the normal limit, or 70, provided that there was a consecutive reading of twice the normal limit recorded at least one week after the initial reading. In category B, the progression criteria were the same, with the exception that the progression limit had to increase to be at least twice the lowest CA125 reading in the treatment line. In many patients, CA125 levels would continue to rise during the early platinum treatments due to the lag between treatment administration and effect. To avoid false early progressions, CA125 readings were only eligible for progression if they occurred after the last treatment in the line. There remained some cases where CA125 readings failed to reach the threshold for progression before the next line of therapy began. In these cases the beginning of the next line was chosen as the date of progression. Using the progression date calculated for each patient, the progression-free survival (PFS) was defined as the number of days from the date of diagnosis to the date of progression.

##### PFS for doxorubicin treatment

The start of the progression-free survival interval in the doxorubicin line was considered as the date of the 1st cycle of treatment. If the patient responded to doxorubicin following the CA125 definitions of response, the progression date was calculated using the same criteria as for platinum, but in this case, progression was permitted to occur during treatment but always after the 3rd cycle of doxorubicin, since there are not many readings post treatment. For nonresponding patients, the date of the 3rd cycle of treatment was used. Response to doxorubicin was identified according to CA125 levels. A response was deemed to have occurred if there was at least a 50% reduction in CA125 levels from a pretreatment sample that was at least twice the upper normal limit. This pretreatment reading must have been taken within 14 days before the first cycle of doxorubicin, and as a result of this requirement, not all patients were able to be evaluated for doxorubicin response. If a 50% reduction was registered, this level must have been maintained for at least 28 days. Any readings taken in those 28 days must have had a CA125 level lower than the previous reading, or have been no more than 10% higher to account for noise. If these requirements were not met before the 1st cycle of the next line of treatment, then the patient was considered as having not responded. In some cases, a patient was treated with another anthracycline such as epirubicin before the first doxorubicin line. In this case, the first anthracycline line was used. After the progression date was calculated for each patient, the PFS was defined as the number of days from the date of the start of the treatment to the date of progression.

##### PFS for paclitaxel treatment

The PFS for paclitaxel was defined as the number of days from the date of the 1st cycle of treatment to the date of progression. The progression date was calculated following the CA125 progression criteria as for platinum, with the exception that progression was permitted to occur during treatment but always after the 3rd cycle of paclitaxel, since there are not many readings post treatment. In those cases where the CA125 readings did not meet the threshold before the next line of therapy began, the date of the next line was chosen as the date of progression.

#### Time to Treatment Failure (TTF)

Due to the absence of CA125 levels in the TCGA clinical data and limited information regarding cancer progression, we used time to treatment failure (TTF) as a proxy measure for PFS.

The TTF data for the TCGA-OV cohort was previously curated[4]. For this cohort, TTF was calculated as the duration from the initiation of one treatment to the initiation of the subsequent treatment, unless there was alternative data indicating disease progression (i.e. metastasis/relapse). In cases where the treatments were deemed adjuvant in nature, treatment lines were manually curated to combine them.

For the other cancer-specific TCGA cohorts, TTF was computed following the same criteria, with the exception that manual curation of treatment lines was not performed and that treatment lines were joined only if they overlapped in terms of the dates of administration.

### Survival analysis

#### Retrospective randomised control study

To evaluate the predictive accuracy of our technology, we conducted a survival analysis that replicated a randomised controlled study design. Initially, we applied our signature-based classifiers to classify patients based on their predicted response. Subsequently, patients belonging to the predicted resistant and predicted sensitive groups were retrospectively assigned, in a manner that mimicked random allocation, to either the experimental arm (receiving single-agent chemotherapy) or the control arm (receiving standard-of-care therapies). Due to sample size requirements, this retrospective randomised controlled study design was only possible to be applied in ovarian and breast TCGA patients treated with taxanes, and ovarian patients treated with doxorubicin (**Extended Data Figure 3**).

For patients assigned to the experimental arm, we filtered out any other treatment regimens and retained only the initial line of single-agent experimental chemotherapy. In the control arm, we included patients who had never received single-agent chemotherapy. Unlike the experimental arm, the control arm allowed for multiple treatment regimens per patient. To mitigate the potential influence of clinical trial drugs or atypical therapies on the survival analysis results, we considered only the five most common treatments or treatment combinations that did not involve the chemotherapy being tested in the experimental arm as standard-of-care treatments, based on the clinical data from the TCGA cohorts. In the context of breast cancers, HER2 inhibitors were omitted from the control arm since this therapeutic approach exhibits superior performance when compared to any other therapies[25–30]. Finally, in order to enable a meaningful comparison between treatment lines, we implemented successive filtering procedures on both the experimental and control arms to ensure that no treatment lines were present in one arm but absent in the other.

Cox proportional hazards models (function *coxph* from the *survival* package in R[31]) were used to compare the TTF of the experimental and control arms in both the predicted resistant and predicted sensitive groups (**Supplementary Figures 44-49**). Cox proportional hazard models were corrected by both age at diagnosis and tumour stage, and further stratified by treatment line. In the case of breast cancers, patients with tumour stage I were filtered out, and survival analyses were additionally stratified based on tumour subtype. In the case of the doxorubicin analysis in ovarian patients, the response to platinum at 1st line was also used as a strata covariate, where patients were considered non-responders if they were moved on to 2nd treatment line within 6 months of the platinum treatment start date.

#### Retrospective study

Cox proportional hazards models were used to compare the PFS between ovarian cancer patients classified as resistant or sensitive included in the OV04 study (**Figure 2**). Cox proportional hazard models were corrected by both age at diagnosis and tumour stage. Age at diagnosis was treated as a continuous variable, whereas tumour stage was stratified into two groups (tumour stages I-III *versus* IV). For the doxorubicin response analysis, 1st line platinum PFS was also added to the model as an interaction term with the response prediction.

We extended the assessment of our technology pan-cancer by performing survival analysis in multiple cancer-specific TCGA cohorts, including ovarian, breast, head and neck, cervical and uterine (**Extended Data Figure 4**). In this case, survival time was measured as TTF rather than PFS. Cox proportional hazard models were also corrected by both age at diagnosis and tumour stage (**Supplementary Figures 50-57**). Age at diagnosis was used as a continuous variable, and tumour stage was grouped into 4 categories corresponding to the main tumour stages. As in the retrospective randomised control study, survival analyses for breast cancers were stratified based on tumour subtype.

#### Annotation of breast tumour subtypes

The TCGA-BRCA cohort underwent annotation for triple-negative status, estrogen receptor (ER) and progesterone receptor (PR) expression, as well as human epidermal growth factor receptor (HER2) amplification.

For the triple-negative status annotation, we used three distinct datasets which incorporated RNA and microRNA expression data, histologic analysis, mutation, copy number, epigenetic, proteomic, and phospho-proteomic information to identify triple-negative status in TCGA-BRCA samples[32–35]. Tumours were classified as triple-negative breast cancers (TNBC) only if all three datasets agreed on this annotation.

Two datasets were also used to separately identify ER, PR, and HER2 status[32,33]. Tumours were defined as positive for ER, PR, or HER2 only if both datasets agreed on their annotation. A patient could be labelled positive for more than one of the ER, PR, and HER2 annotations.

### Concordance between tissue and plasma pairs

We assessed concordance between tissue and plasma samples collected from the same patient by means of absolute copy number profiles (by using the *getDifference* function from our *CNpare* tool in R[36]), signature activities (by computing cosine similarity), and drug response predictions (by observing classification concordance). **Supplementary Figure 58** and **Figure 4b** illustrate differences between matched tissue- and plasma-derived copy number profiles of all patients; while **Extended Data Figure 5** shows signature composition concordance between pairs.
