## Supplementary figures and tables for "Predicting response to cytotoxic chemotherapy"

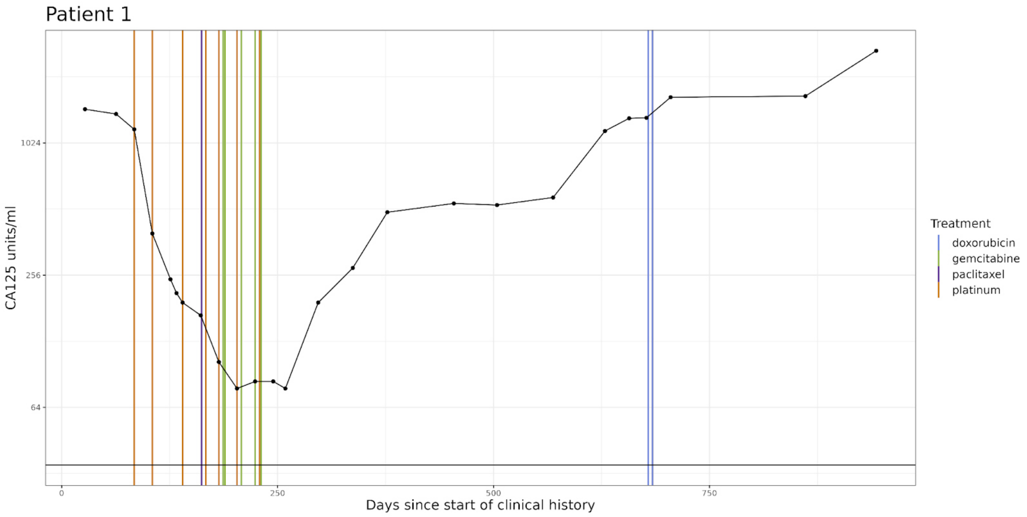


**Supplementary Figure 1.** Clinical history plot for OV04 patient 1. Blood serum CA125 levels are shown over time, with the horizontal bars denoting cycles of treatment. The horizontal bar at 35 units/ml denotes the threshold between ‘normal’ and ‘abnormal’ CA125 readings. In cases where multiple treatments are given on the same day, the treatment date is shifted slightly to show all treatments.

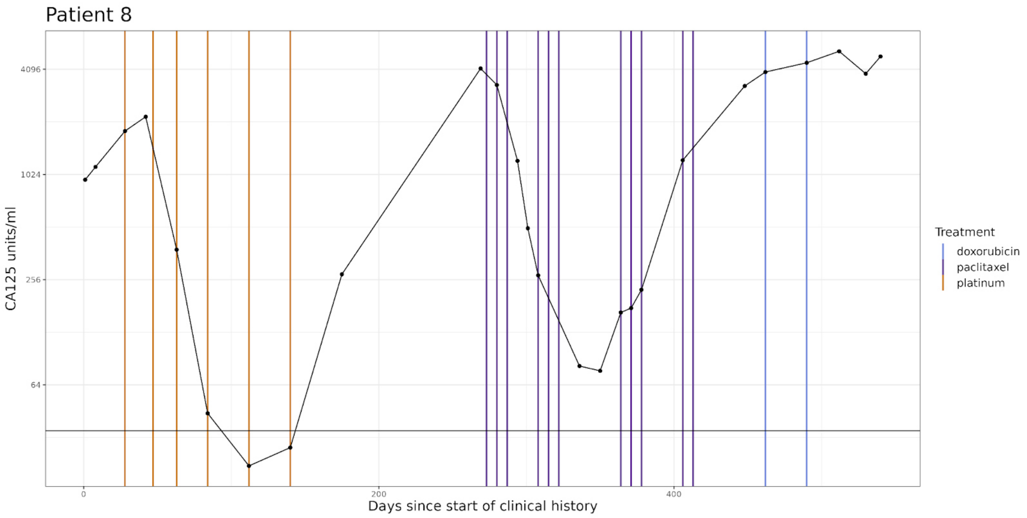


**Supplementary Figure 2.** Clinical history plot for OV04 patient 8. Blood serum CA125 levels are shown over time, with the horizontal bars denoting cycles of treatment. The horizontal bar at 35 units/ml denotes the threshold between ‘normal’ and ‘abnormal’ CA125 readings. In cases where multiple treatments are given on the same day, the treatment date is shifted slightly to show all treatments.

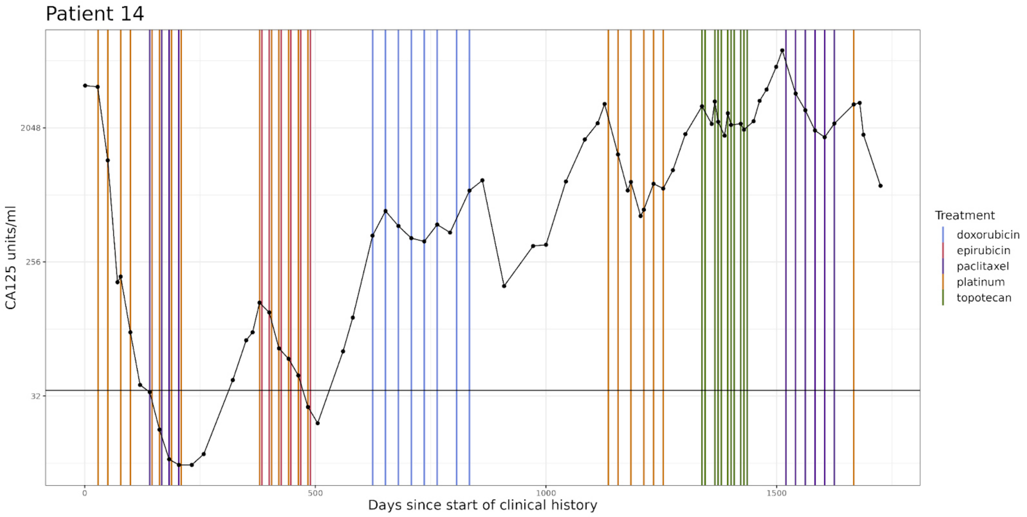


**Supplementary Figure 3.** Clinical history plot for OV04 patient 14. Blood serum CA125 levels are shown over time, with the horizontal bars denoting cycles of treatment. The horizontal bar at 35 units/ml denotes the threshold between 'normal' and 'abnormal' CA125 readings. In cases where multiple treatments are given on the same day, the treatment date is shifted slightly to show all treatments.

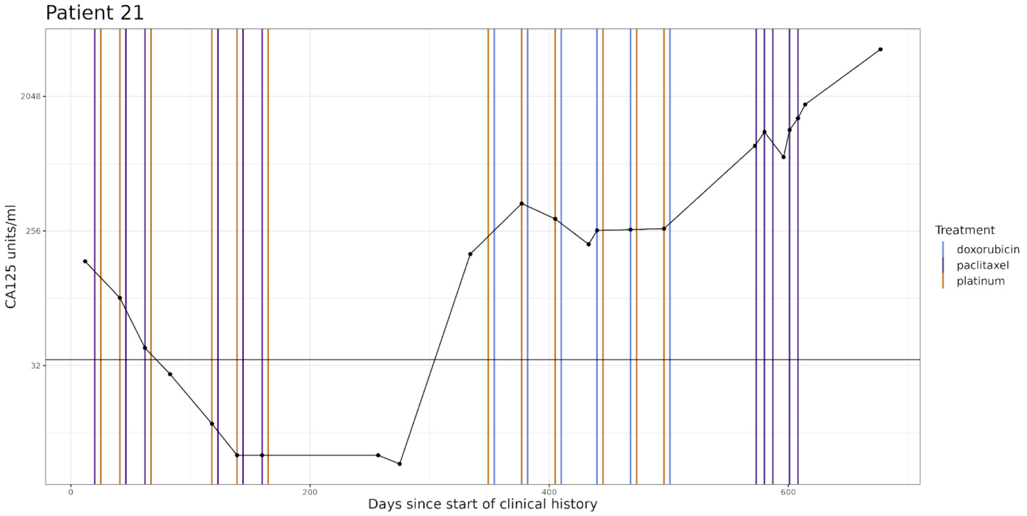


**Supplementary Figure 4.** Clinical history plot for OV04 patient 21. Blood serum CA125 levels are shown over time, with the horizontal bars denoting cycles of treatment. The horizontal bar at 35 units/ml denotes the threshold between 'normal' and 'abnormal' CA125 readings. In cases where multiple treatments are given on the same day, the treatment date is shifted slightly to show all treatments.

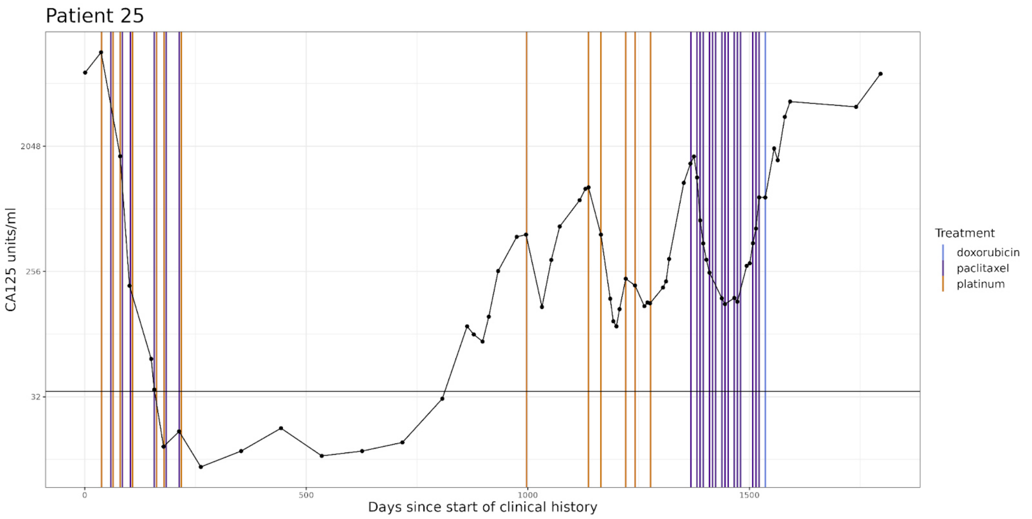


**Supplementary Figure 5.** Clinical history plot for OV04 patient 25. Blood serum CA125 levels are shown over time, with the horizontal bars denoting cycles of treatment. The horizontal bar at 35 units/ml denotes the threshold between 'normal' and 'abnormal' CA125 readings. In cases where multiple treatments are given on the same day, the treatment date is shifted slightly to show all treatments.

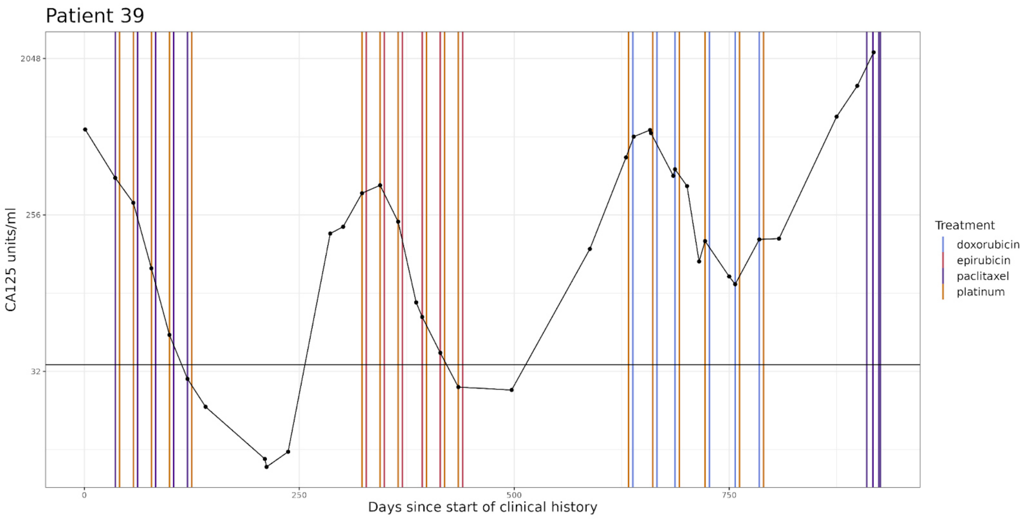


**Supplementary Figure 6.** Clinical history plot for OV04 patient 39. Blood serum CA125 levels are shown over time, with the horizontal bars denoting cycles of treatment. The horizontal bar at 35 units/ml denotes the threshold between 'normal' and 'abnormal' CA125 readings. In cases where multiple treatments are given on the same day, the treatment date is shifted slightly to show all treatments.

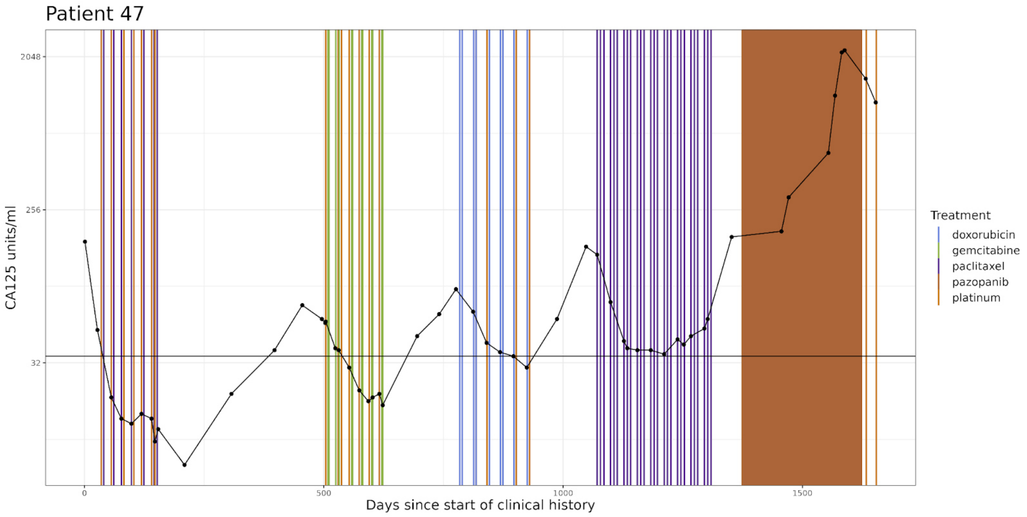


**Supplementary Figure 7.** Clinical history plot for OV04 patient 47. Blood serum CA125 levels are shown over time, with the horizontal bars denoting cycles of treatment. The horizontal bar at 35 units/ml denotes the threshold between ‘normal’ and ‘abnormal’ CA125 readings. In cases where multiple treatments are given on the same day, the treatment date is shifted slightly to show all treatments.

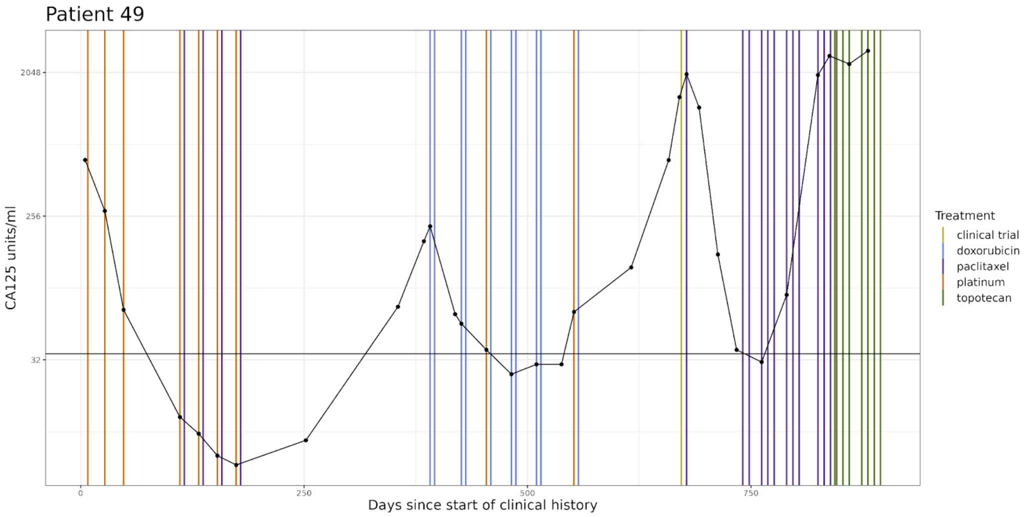


**Supplementary Figure 8.** Clinical history plot for OV04 patient 49. Blood serum CA125 levels are shown over time, with the horizontal bars denoting cycles of treatment. The horizontal bar at 35 units/ml denotes the threshold between ‘normal’ and ‘abnormal’ CA125 readings. In cases where multiple treatments are given on the same day, the treatment date is shifted slightly to show all treatments.

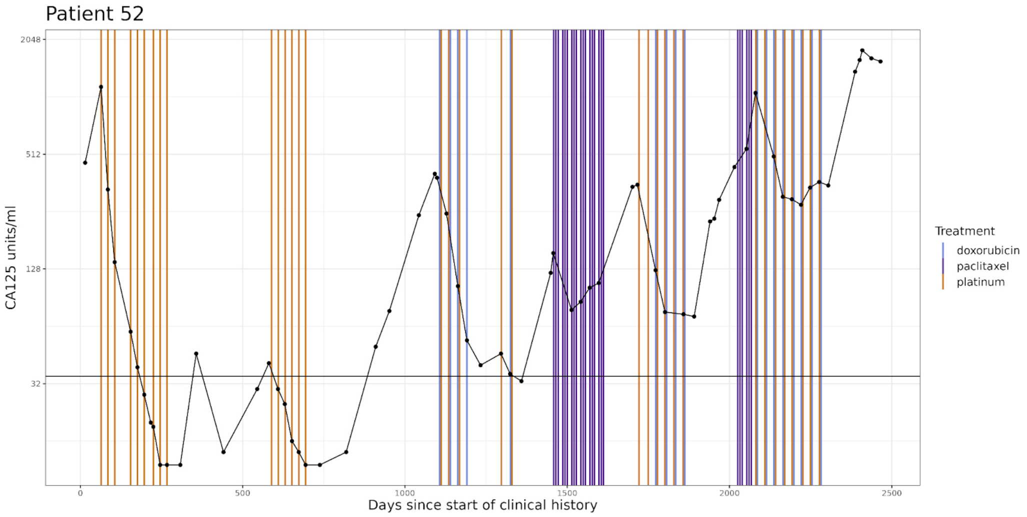


**Supplementary Figure 9.** Clinical history plot for OV04 patient 52. Blood serum CA125 levels are shown over time, with the horizontal bars denoting cycles of treatment. The horizontal bar at 35 units/ml denotes the threshold between ‘normal’ and ‘abnormal’ CA125 readings. In cases where multiple treatments are given on the same day, the treatment date is shifted slightly to show all treatments.

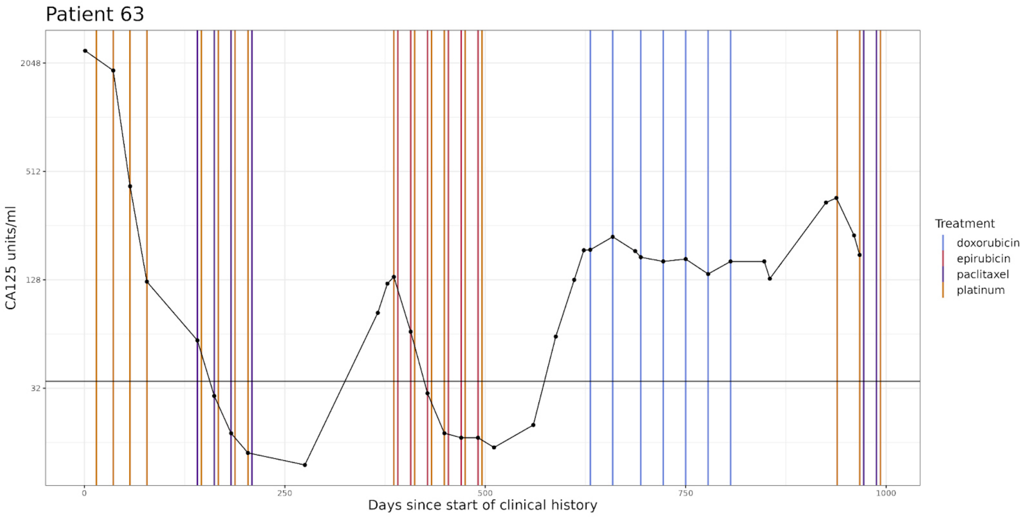


**Supplementary Figure 10.** Clinical history plot for OV04 patient 63. Blood serum CA125 levels are shown over time, with the horizontal bars denoting cycles of treatment. The horizontal bar at 35 units/ml denotes the threshold between ‘normal’ and ‘abnormal’ CA125 readings. In cases where multiple treatments are given on the same day, the treatment date is shifted slightly to show all treatments.

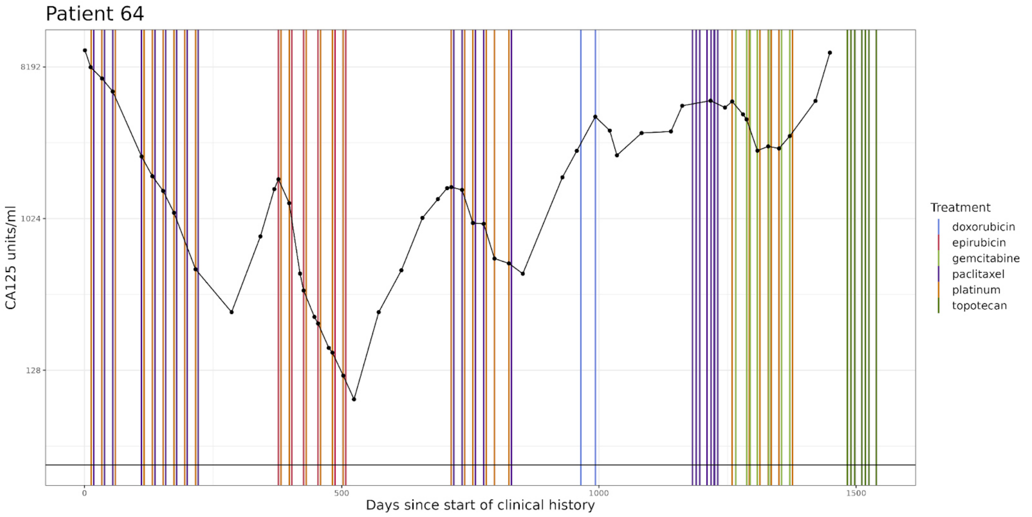


**Supplementary Figure 11.** Clinical history plot for OV04 patient 64. Blood serum CA125 levels are shown over time, with the horizontal bars denoting cycles of treatment. The horizontal bar at 35 units/ml denotes the threshold between ‘normal’ and ‘abnormal’ CA125 readings. In cases where multiple treatments are given on the same day, the treatment date is shifted slightly to show all treatments.

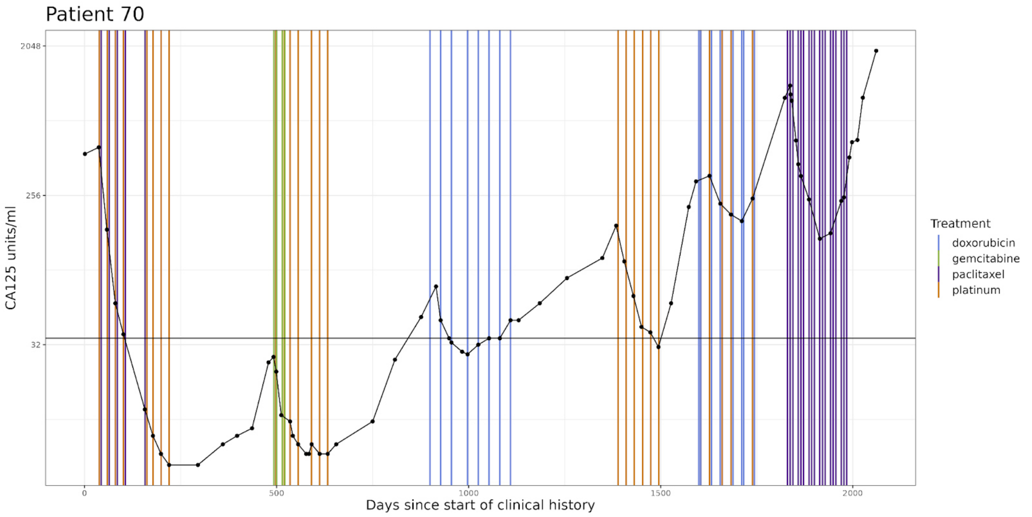


**Supplementary Figure 12.** Clinical history plot for OV04 patient 70. Blood serum CA125 levels are shown over time, with the horizontal bars denoting cycles of treatment. The horizontal bar at 35 units/ml denotes the threshold between ‘normal’ and ‘abnormal’ CA125 readings. In cases where multiple treatments are given on the same day, the treatment date is shifted slightly to show all treatments.

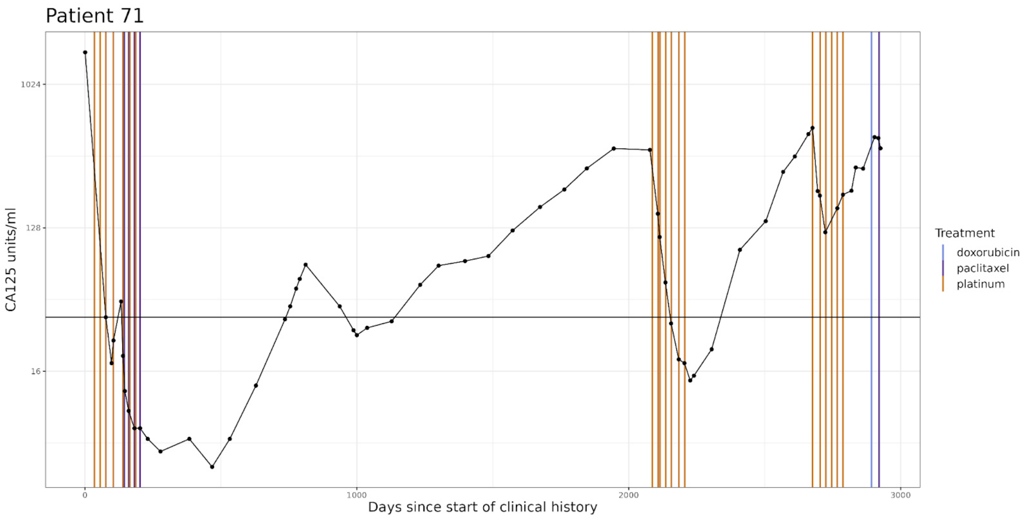


**Supplementary Figure 13.** Clinical history plot for OV04 patient 71. Blood serum CA125 levels are shown over time, with the horizontal bars denoting cycles of treatment. The horizontal bar at 35 units/ml denotes the threshold between ‘normal’ and ‘abnormal’ CA125 readings. In cases where multiple treatments are given on the same day, the treatment date is shifted slightly to show all treatments.

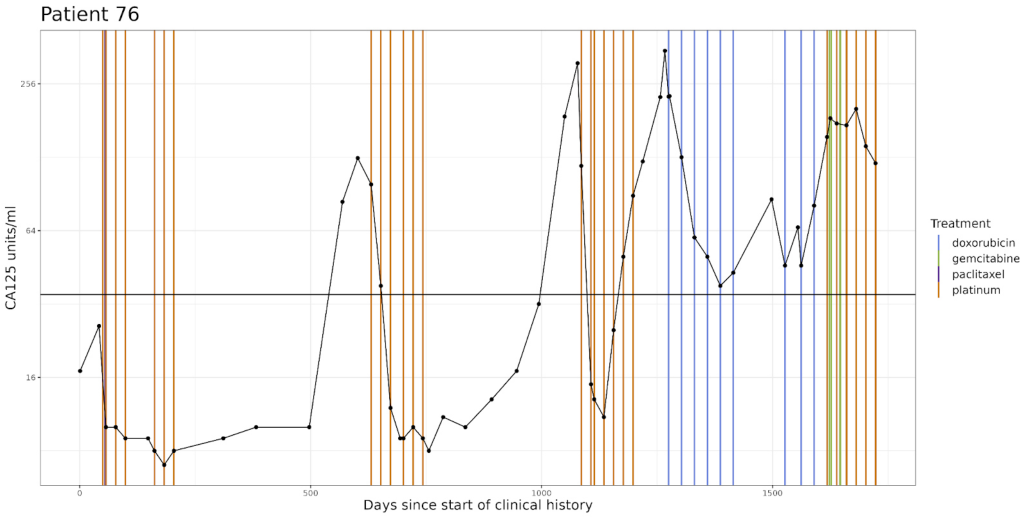


**Supplementary Figure 14.** Clinical history plot for OV04 patient 76. Blood serum CA125 levels are shown over time, with the horizontal bars denoting cycles of treatment. The horizontal bar at 35 units/ml denotes the threshold between ‘normal’ and ‘abnormal’ CA125 readings. In cases where multiple treatments are given on the same day, the treatment date is shifted slightly to show all treatments.

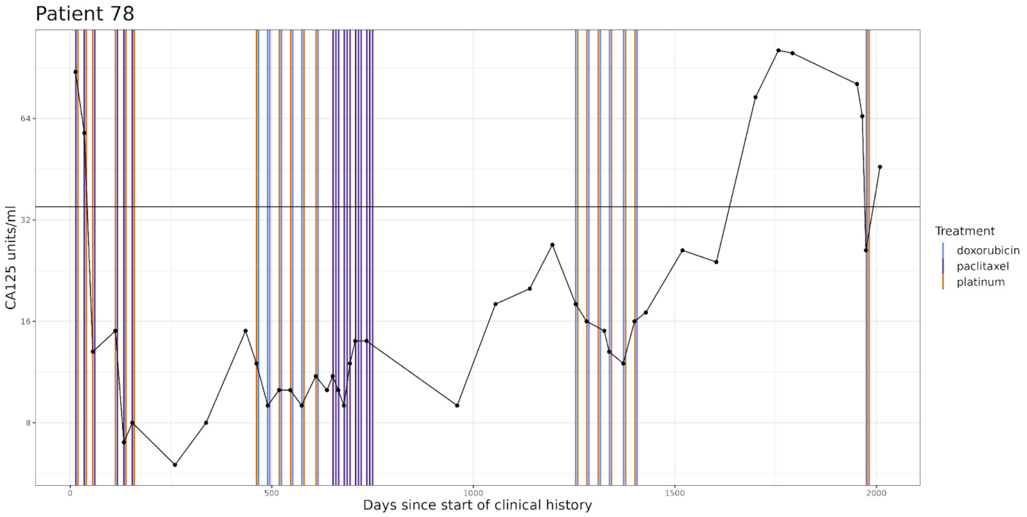


**Supplementary Figure 15.** Clinical history plot for OV04 patient 78. Blood serum CA125 levels are shown over time, with the horizontal bars denoting cycles of treatment. The horizontal bar at 35 units/ml denotes the threshold between ‘normal’ and ‘abnormal’ CA125 readings. In cases where multiple treatments are given on the same day, the treatment date is shifted slightly to show all treatments.

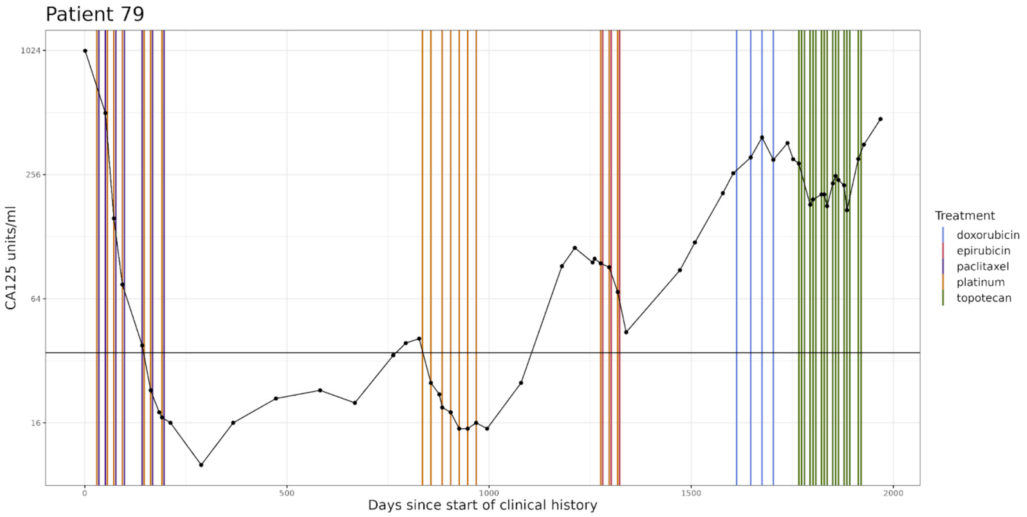


**Supplementary Figure 16.** Clinical history plot for OV04 patient 79. Blood serum CA125 levels are shown over time, with the horizontal bars denoting cycles of treatment. The horizontal bar at 35 units/ml denotes the threshold between ‘normal’ and ‘abnormal’ CA125 readings. In cases where multiple treatments are given on the same day, the treatment date is shifted slightly to show all treatments.

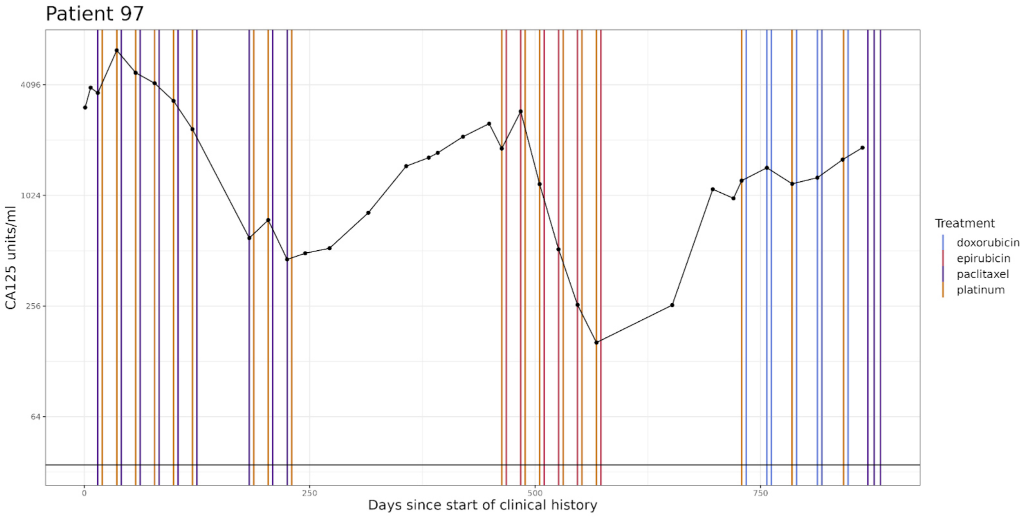


**Supplementary Figure 17.** Clinical history plot for OV04 patient 97. Blood serum CA125 levels are shown over time, with the horizontal bars denoting cycles of treatment. The horizontal bar at 35 units/ml denotes the threshold between ‘normal’ and ‘abnormal’ CA125 readings. In cases where multiple treatments are given on the same day, the treatment date is shifted slightly to show all treatments.

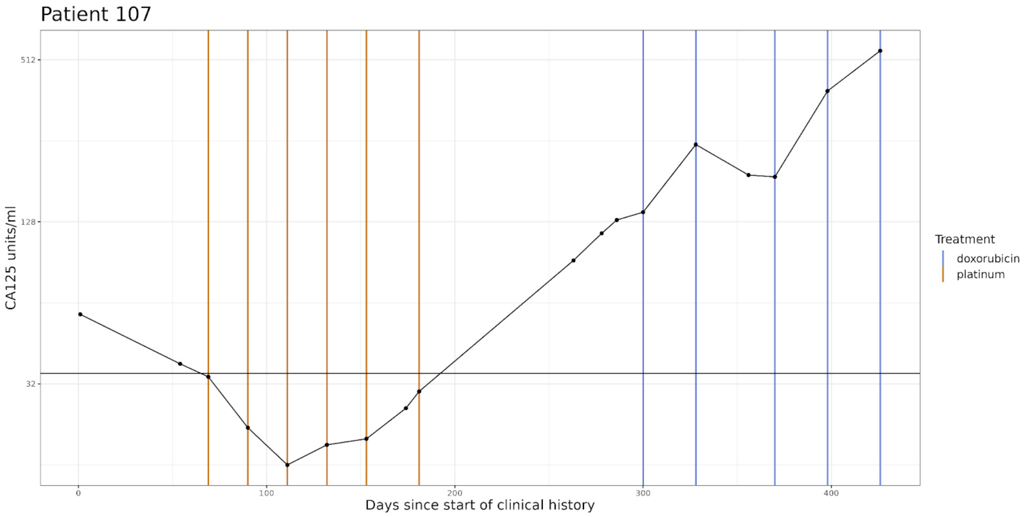


**Supplementary Figure 18.** Clinical history plot for OV04 patient 107. Blood serum CA125 levels are shown over time, with the horizontal bars denoting cycles of treatment. The horizontal bar at 35 units/ml denotes the threshold between ‘normal’ and ‘abnormal’ CA125 readings. In cases where multiple treatments are given on the same day, the treatment date is shifted slightly to show all treatments.

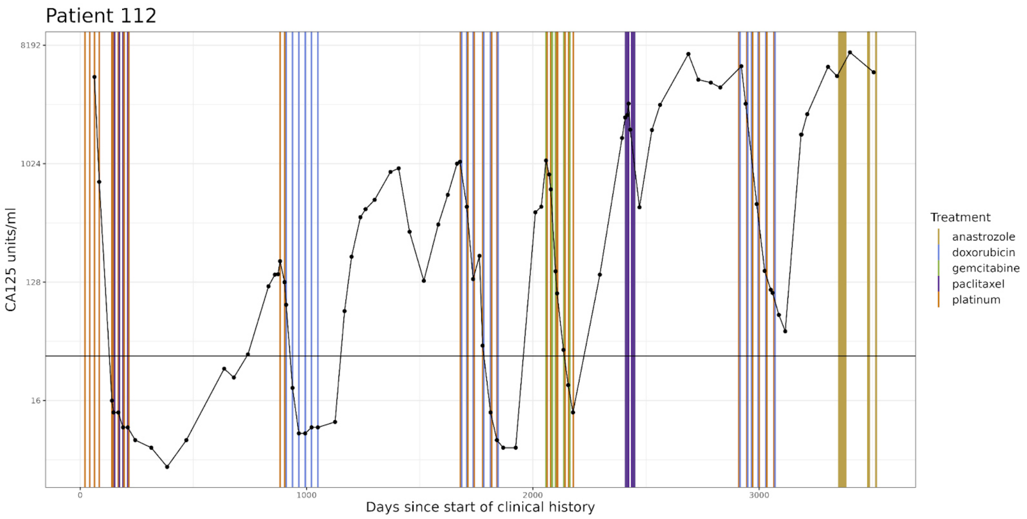


**Supplementary Figure 19.** Clinical history plot for OV04 patient 112. Blood serum CA125 levels are shown over time, with the horizontal bars denoting cycles of treatment. The horizontal bar at 35 units/ml denotes the threshold between ‘normal’ and ‘abnormal’ CA125 readings. In cases where multiple treatments are given on the same day, the treatment date is shifted slightly to show all treatments.

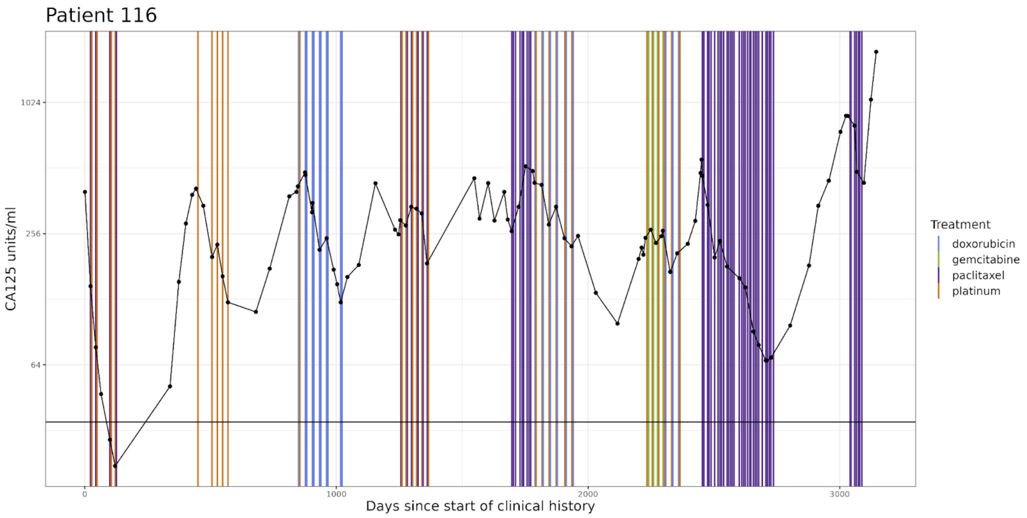


**Supplementary Figure 20.** Clinical history plot for OV04 patient 116. Blood serum CA125 levels are shown over time, with the horizontal bars denoting cycles of treatment. The horizontal bar at 35 units/ml denotes the threshold between ‘normal’ and ‘abnormal’ CA125 readings. In cases where multiple treatments are given on the same day, the treatment date is shifted slightly to show all treatments.

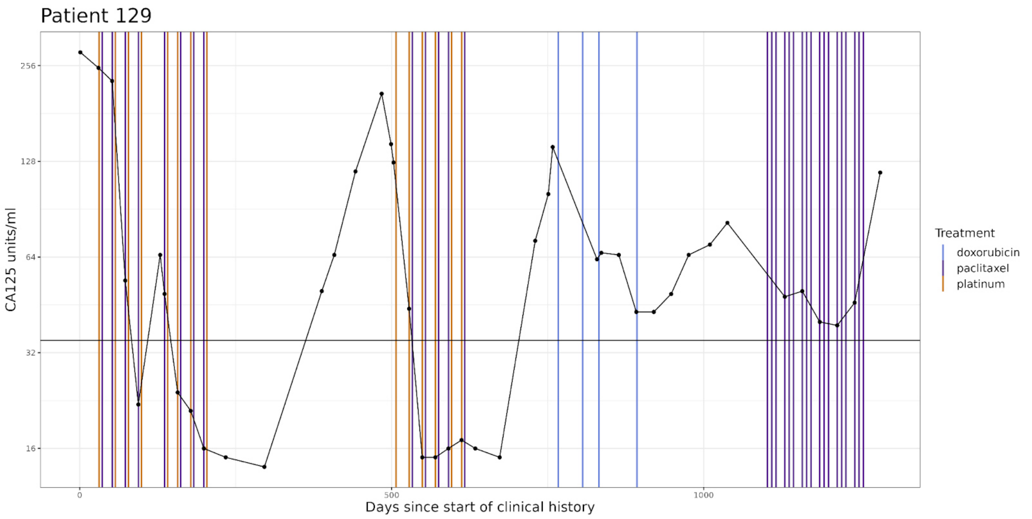


**Supplementary Figure 21.** Clinical history plot for OV04 patient 129. Blood serum CA125 levels are shown over time, with the horizontal bars denoting cycles of treatment. The horizontal bar at 35 units/ml denotes the threshold between 'normal' and 'abnormal' CA125 readings. In cases where multiple treatments are given on the same day, the treatment date is shifted slightly to show all treatments.

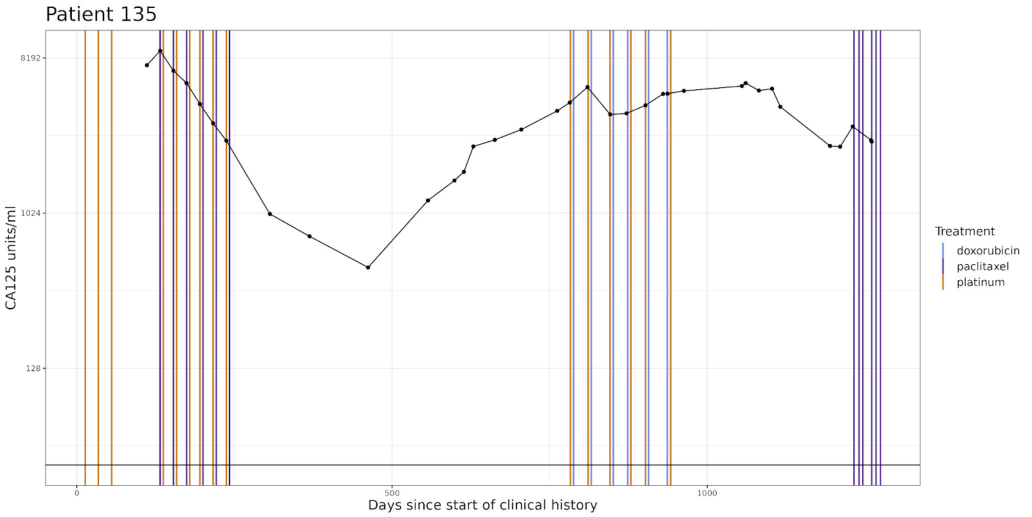


**Supplementary Figure 22.** Clinical history plot for OV04 patient 135. Blood serum CA125 levels are shown over time, with the horizontal bars denoting cycles of treatment. The horizontal bar at 35 units/ml denotes the threshold between 'normal' and 'abnormal' CA125 readings. In cases where multiple treatments are given on the same day, the treatment date is shifted slightly to show all treatments.

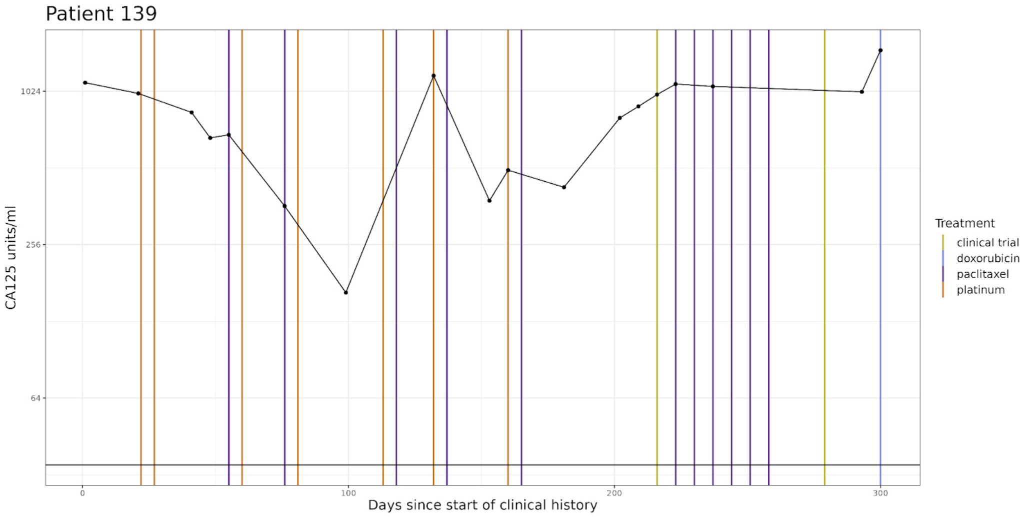


**Supplementary Figure 23.** Clinical history plot for OV04 patient 139. Blood serum CA125 levels are shown over time, with the horizontal bars denoting cycles of treatment. The horizontal bar at 35 units/ml denotes the threshold between ‘normal’ and ‘abnormal’ CA125 readings. In cases where multiple treatments are given on the same day, the treatment date is shifted slightly to show all treatments.

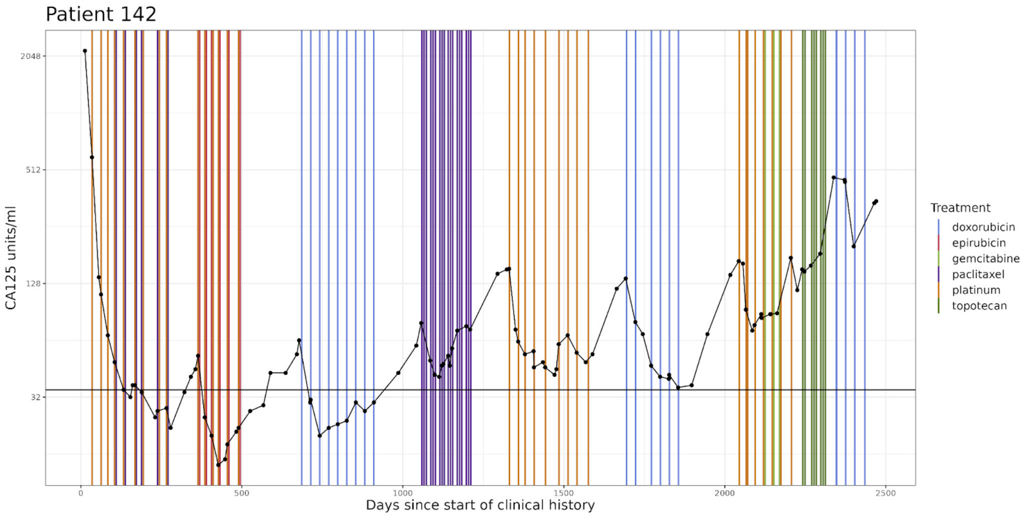


**Supplementary Figure 24.** Clinical history plot for OV04 patient 142. Blood serum CA125 levels are shown over time, with the horizontal bars denoting cycles of treatment. The horizontal bar at 35 units/ml denotes the threshold between ‘normal’ and ‘abnormal’ CA125 readings. In cases where multiple treatments are given on the same day, the treatment date is shifted slightly to show all treatments.

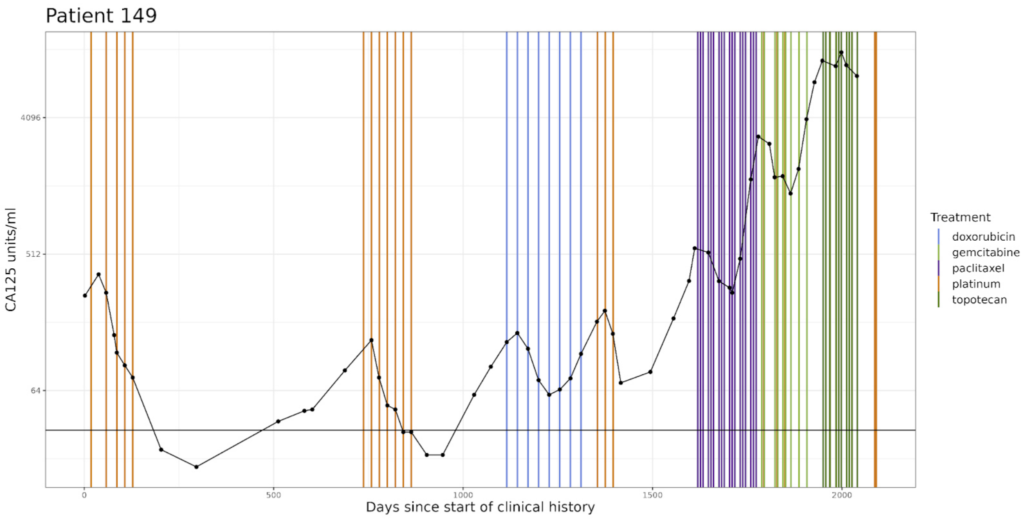


**Supplementary Figure 25.** Clinical history plot for OV04 patient 149. Blood serum CA125 levels are shown over time, with the horizontal bars denoting cycles of treatment. The horizontal bar at 35 units/ml denotes the threshold between ‘normal’ and ‘abnormal’ CA125 readings. In cases where multiple treatments are given on the same day, the treatment date is shifted slightly to show all treatments.

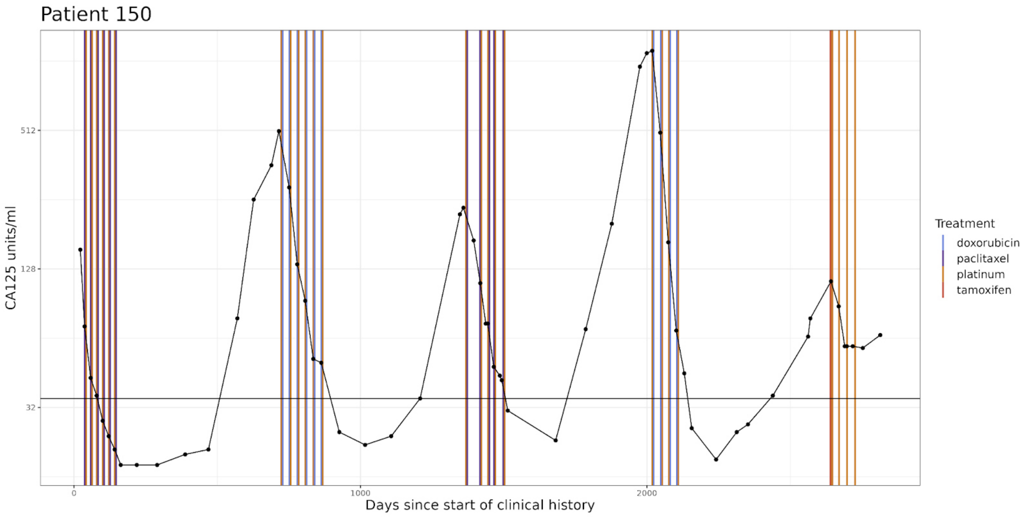


**Supplementary Figure 26.** Clinical history plot for OV04 patient 150. Blood serum CA125 levels are shown over time, with the horizontal bars denoting cycles of treatment. The horizontal bar at 35 units/ml denotes the threshold between ‘normal’ and ‘abnormal’ CA125 readings. In cases where multiple treatments are given on the same day, the treatment date is shifted slightly to show all treatments.

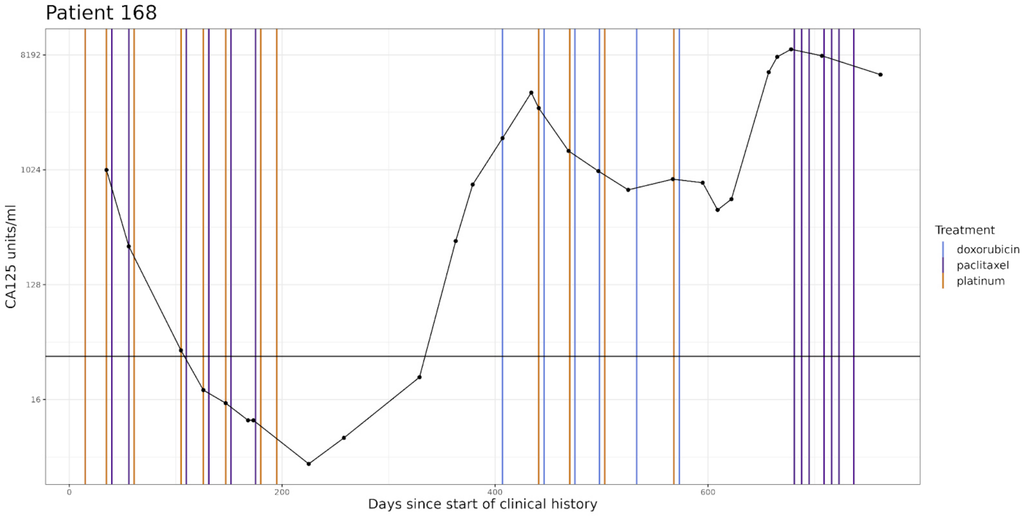


**Supplementary Figure 27.** Clinical history plot for OV04 patient 168. Blood serum CA125 levels are shown over time, with the horizontal bars denoting cycles of treatment. The horizontal bar at 35 units/ml denotes the threshold between ‘normal’ and ‘abnormal’ CA125 readings. In cases where multiple treatments are given on the same day, the treatment date is shifted slightly to show all treatments.

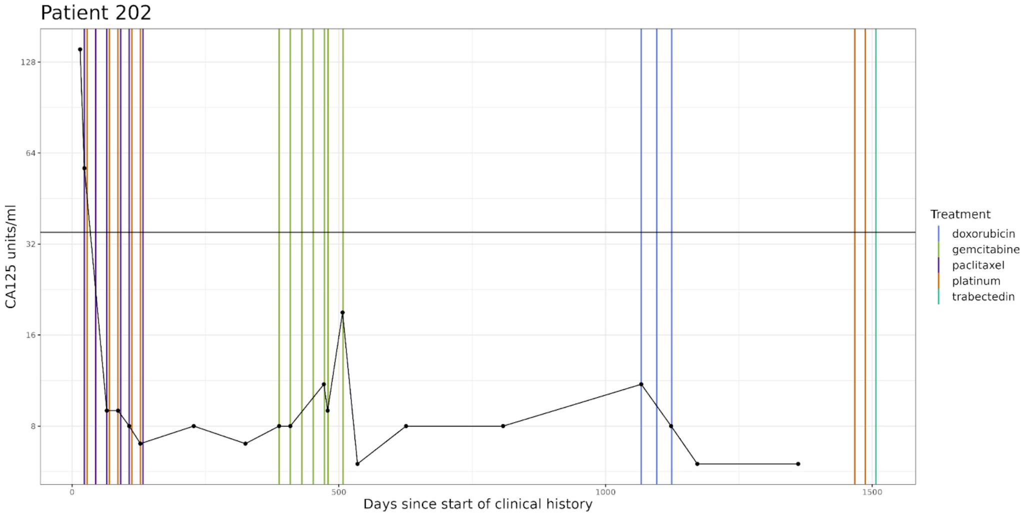


**Supplementary Figure 28.** Clinical history plot for OV04 patient 202. Blood serum CA125 levels are shown over time, with the horizontal bars denoting cycles of treatment. The horizontal bar at 35 units/ml denotes the threshold between ‘normal’ and ‘abnormal’ CA125 readings. In cases where multiple treatments are given on the same day, the treatment date is shifted slightly to show all treatments.

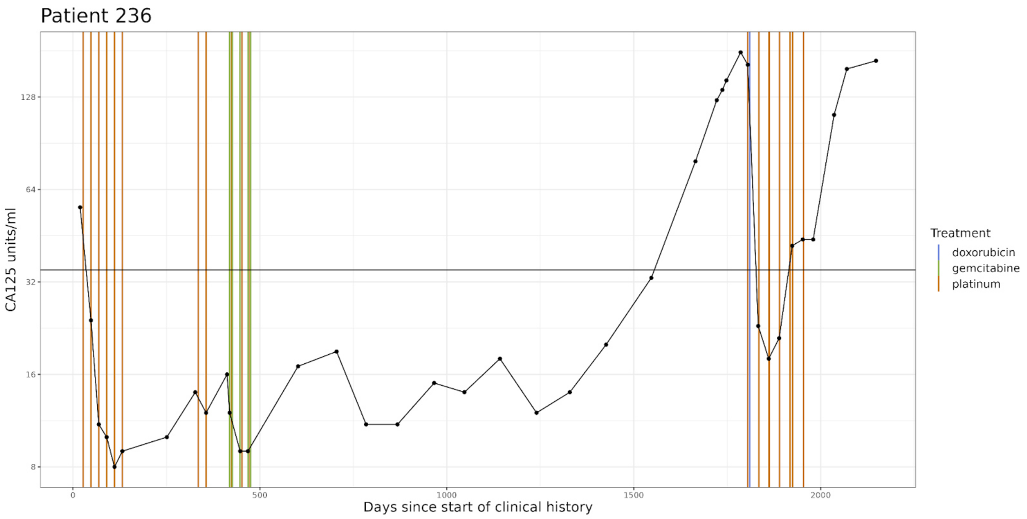


**Supplementary Figure 29.** Clinical history plot for OV04 patient 236. Blood serum CA125 levels are shown over time, with the horizontal bars denoting cycles of treatment. The horizontal bar at 35 units/ml denotes the threshold between ‘normal’ and ‘abnormal’ CA125 readings. In cases where multiple treatments are given on the same day, the treatment date is shifted slightly to show all treatments.

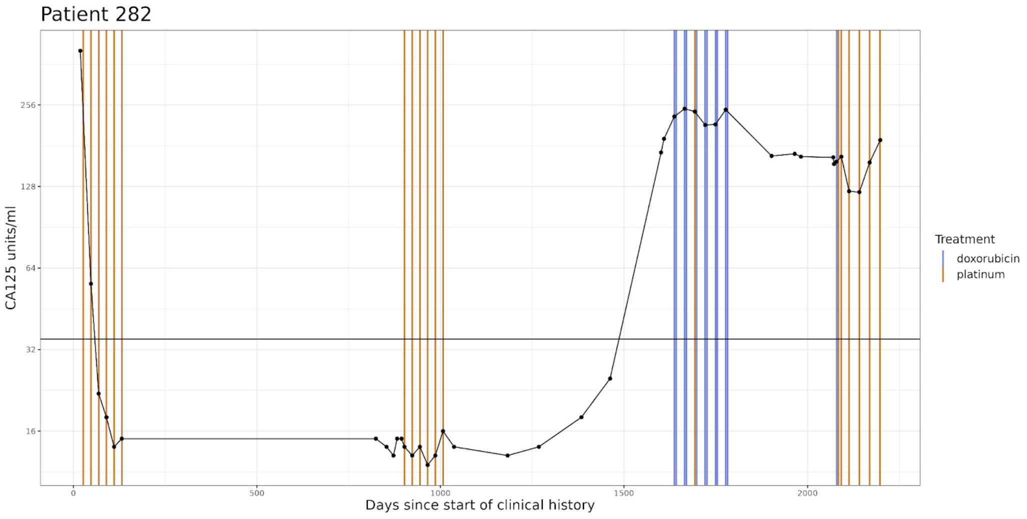


**Supplementary Figure 30.** Clinical history plot for OV04 patient 282. Blood serum CA125 levels are shown over time, with the horizontal bars denoting cycles of treatment. The horizontal bar at 35 units/ml denotes the threshold between ‘normal’ and ‘abnormal’ CA125 readings. In cases where multiple treatments are given on the same day, the treatment date is shifted slightly to show all treatments.

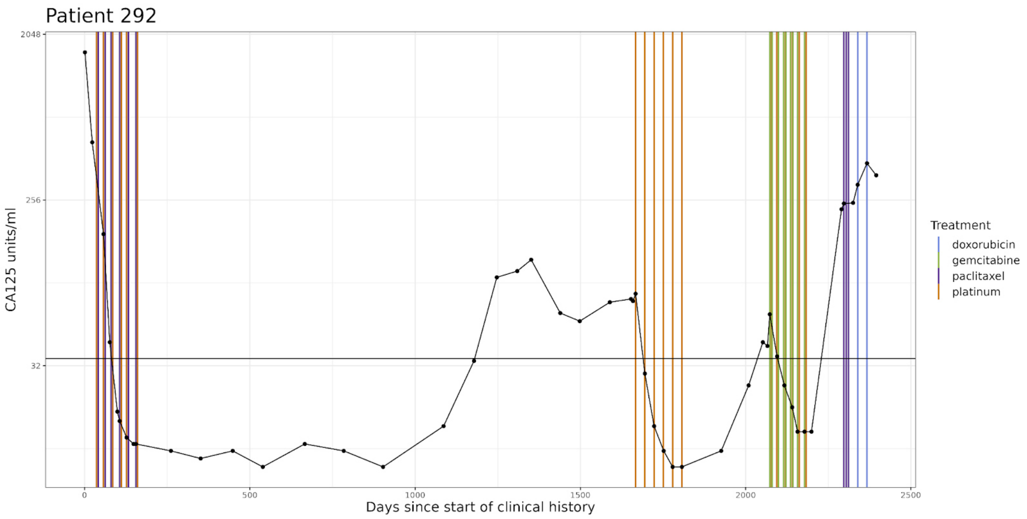


**Supplementary Figure 31.** Clinical history plot for OV04 patient 292. Blood serum CA125 levels are shown over time, with the horizontal bars denoting cycles of treatment. The horizontal bar at 35 units/ml denotes the threshold between ‘normal’ and ‘abnormal’ CA125 readings. In cases where multiple treatments are given on the same day, the treatment date is shifted slightly to show all treatments.

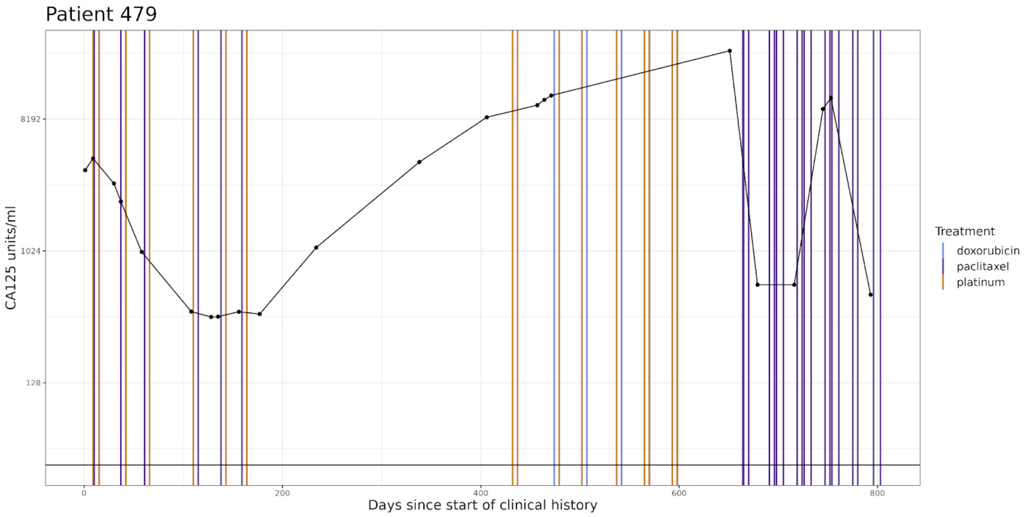


**Supplementary Figure 32.** Clinical history plot for OV04 patient 479. Blood serum CA125 levels are shown over time, with the horizontal bars denoting cycles of treatment. The horizontal bar at 35 units/ml denotes the threshold between ‘normal’ and ‘abnormal’ CA125 readings. In cases where multiple treatments are given on the same day, the treatment date is shifted slightly to show all treatments.

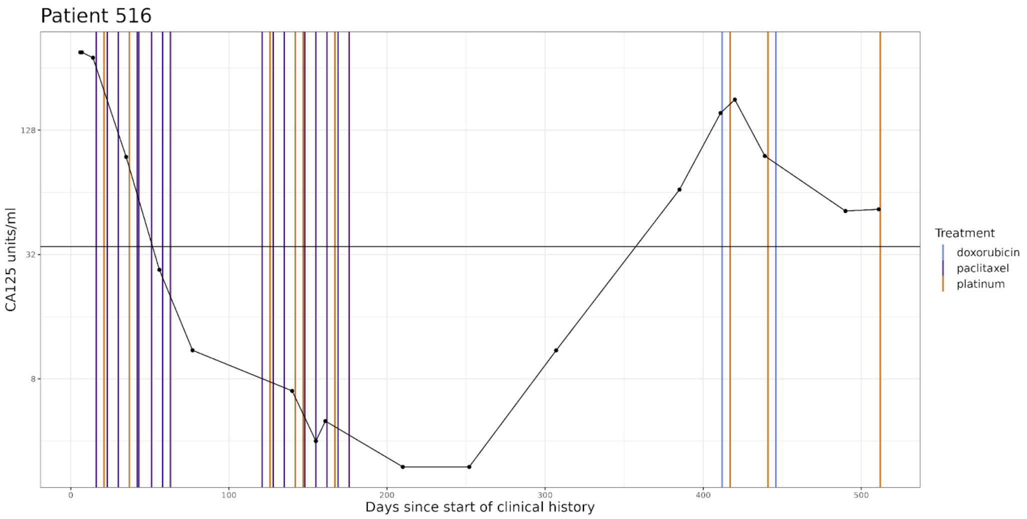


**Supplementary Figure 33.** Clinical history plot for OV04 patient 516. Blood serum CA125 levels are shown over time, with the horizontal bars denoting cycles of treatment. The horizontal bar at 35 units/ml denotes the threshold between 'normal' and 'abnormal' CA125 readings. In cases where multiple treatments are given on the same day, the treatment date is shifted slightly to show all treatments.

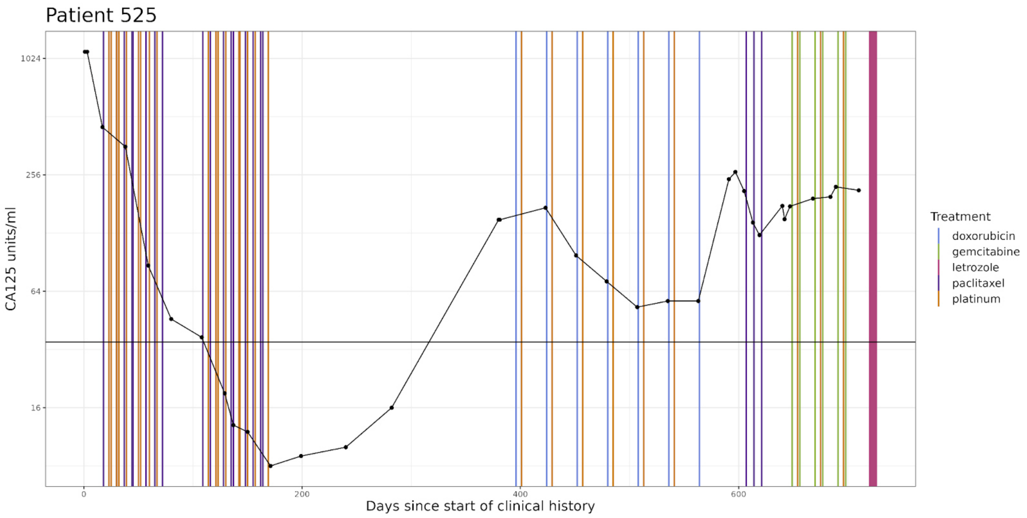


**Supplementary Figure 34.** Clinical history plot for OV04 patient 525. Blood serum CA125 levels are shown over time, with the horizontal bars denoting cycles of treatment. The horizontal bar at 35 units/ml denotes the threshold between 'normal' and 'abnormal' CA125 readings. In cases where multiple treatments are given on the same day, the treatment date is shifted slightly to show all treatments.

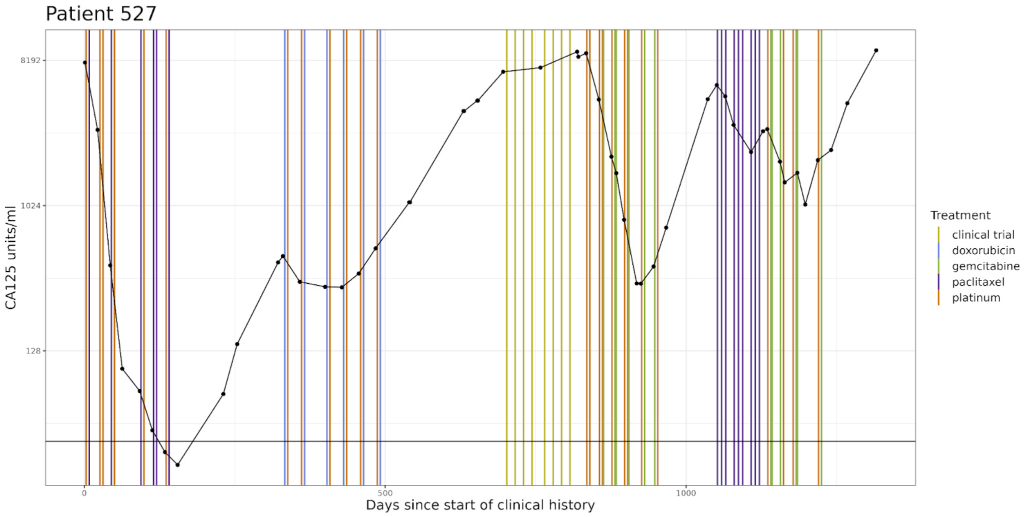


**Supplementary Figure 35.** Clinical history plot for OV04 patient 527. Blood serum CA125 levels are shown over time, with the horizontal bars denoting cycles of treatment. The horizontal bar at 35 units/ml denotes the threshold between ‘normal’ and ‘abnormal’ CA125 readings. In cases where multiple treatments are given on the same day, the treatment date is shifted slightly to show all treatments.

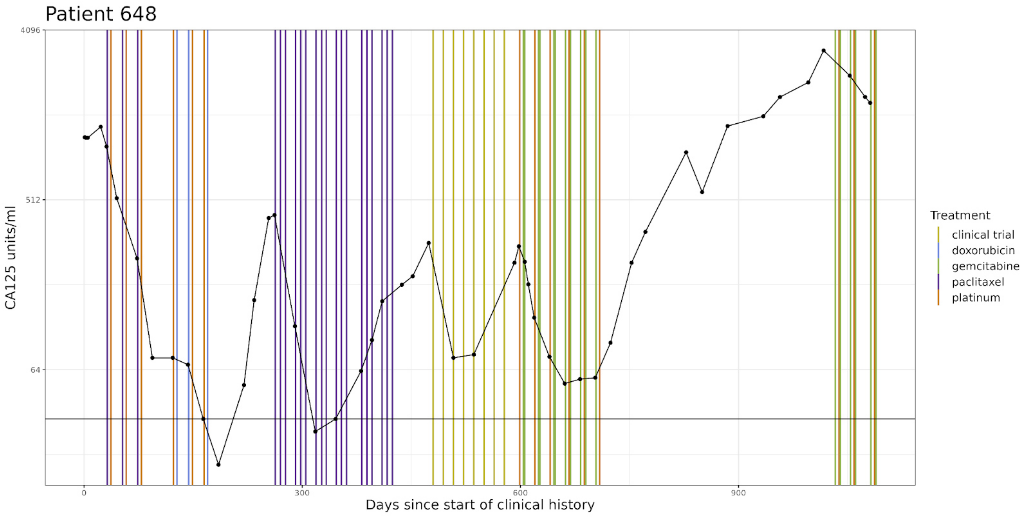


**Supplementary Figure 36.** Clinical history plot for OV04 patient 648. Blood serum CA125 levels are shown over time, with the horizontal bars denoting cycles of treatment. The horizontal bar at 35 units/ml denotes the threshold between ‘normal’ and ‘abnormal’ CA125 readings. In cases where multiple treatments are given on the same day, the treatment date is shifted slightly to show all treatments.

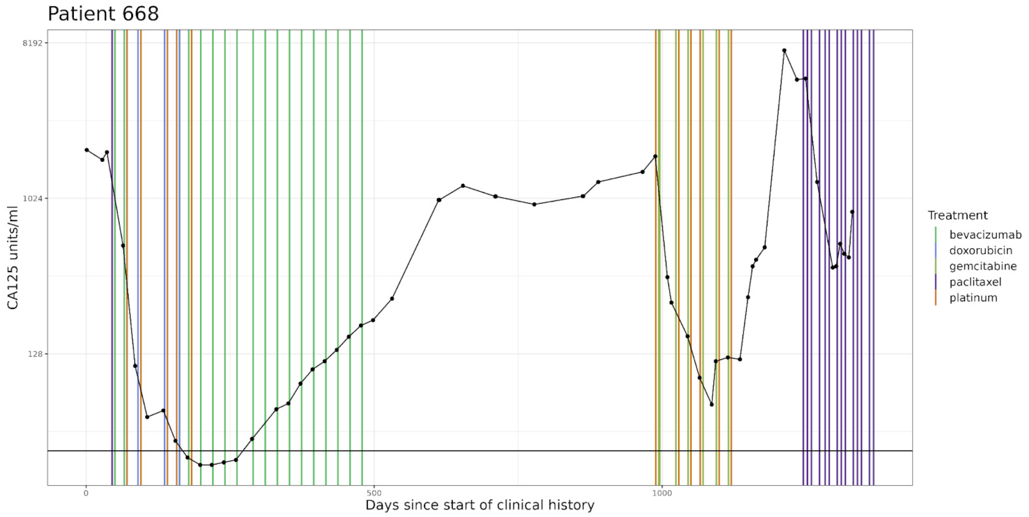


**Supplementary Figure 37.** Clinical history plot for OV04 patient 668. Blood serum CA125 levels are shown over time, with the horizontal bars denoting cycles of treatment. The horizontal bar at 35 units/ml denotes the threshold between ‘normal’ and ‘abnormal’ CA125 readings. In cases where multiple treatments are given on the same day, the treatment date is shifted slightly to show all treatments.

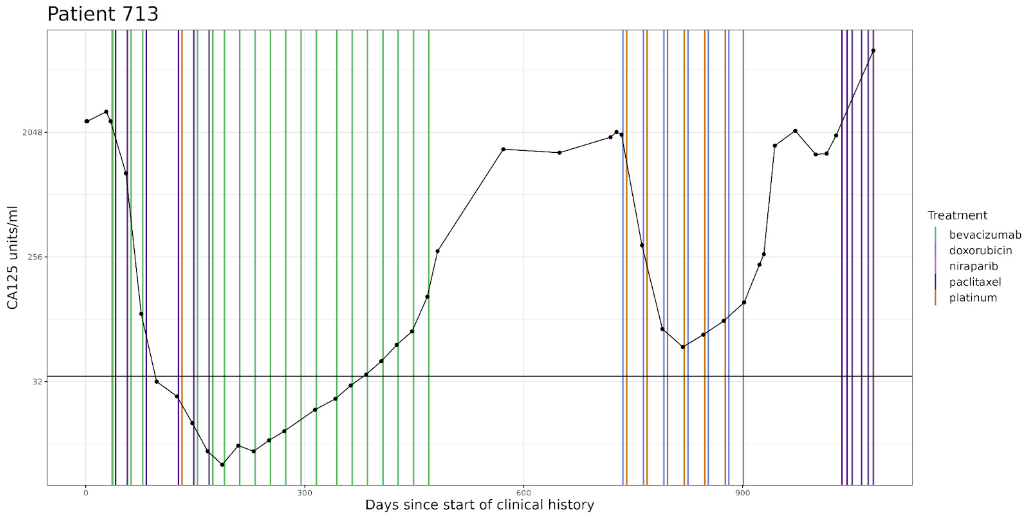


**Supplementary Figure 38.** Clinical history plot for OV04 patient 713. Blood serum CA125 levels are shown over time, with the horizontal bars denoting cycles of treatment. The horizontal bar at 35 units/ml denotes the threshold between ‘normal’ and ‘abnormal’ CA125 readings. In cases where multiple treatments are given on the same day, the treatment date is shifted slightly to show all treatments.

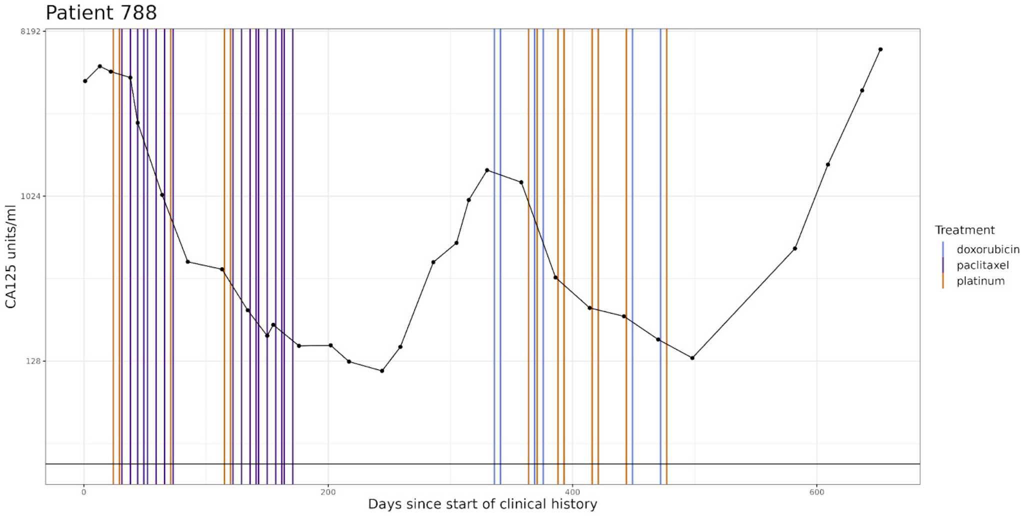


**Supplementary Figure 39.** Clinical history plot for OV04 patient 788. Blood serum CA125 levels are shown over time, with the horizontal bars denoting cycles of treatment. The horizontal bar at 35 units/ml denotes the threshold between ‘normal’ and ‘abnormal’ CA125 readings. In cases where multiple treatments are given on the same day, the treatment date is shifted slightly to show all treatments.

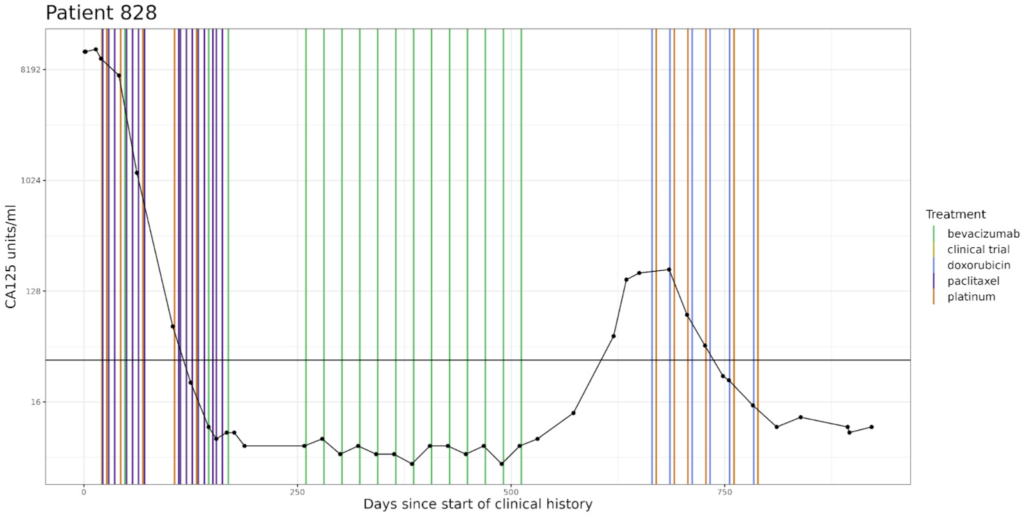


**Supplementary Figure 40.** Clinical history plot for OV04 patient 828. Blood serum CA125 levels are shown over time, with the horizontal bars denoting cycles of treatment. The horizontal bar at 35 units/ml denotes the threshold between ‘normal’ and ‘abnormal’ CA125 readings. In cases where multiple treatments are given on the same day, the treatment date is shifted slightly to show all treatments.

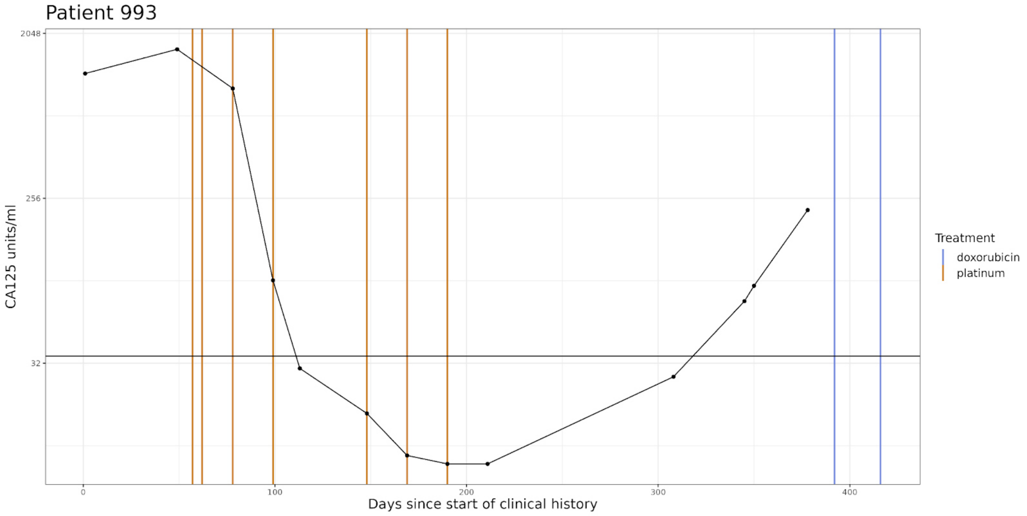


**Supplementary Figure 41.** Clinical history plot for OV04 patient 993. Blood serum CA125 levels are shown over time, with the horizontal bars denoting cycles of treatment. The horizontal bar at 35 units/ml denotes the threshold between 'normal' and 'abnormal' CA125 readings. In cases where multiple treatments are given on the same day, the treatment date is shifted slightly to show all treatments.

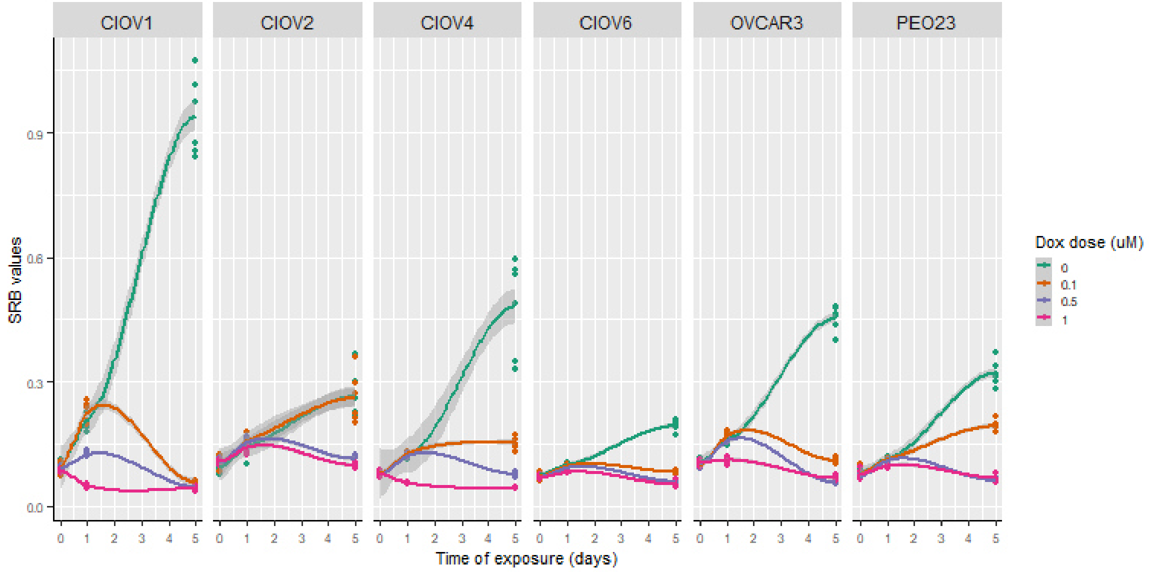


**Supplementary Figure 42. Cell growth rates for ovarian cancer lines under doxorubicin treatment for 5 days.** Sulforhodamine B (SRB) colorimetric assay used for quantifying cell number in culture.

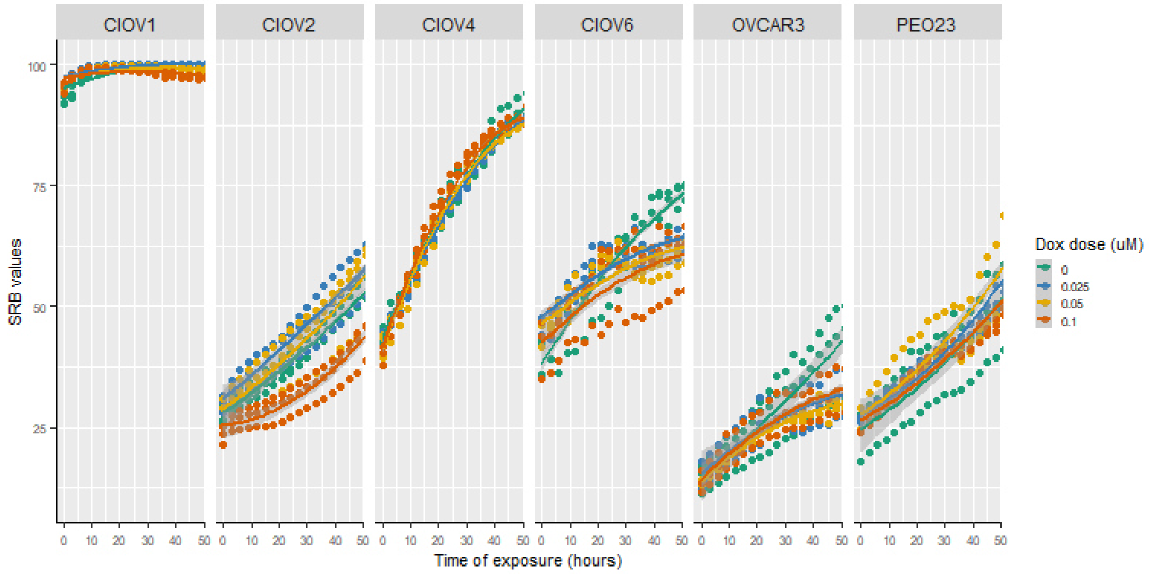


**Supplementary Figure 43. Cell growth rates for ovarian cancer lines under low doses of doxorubicin for 48 hours.** Sulforhodamine B (SRB) colorimetric assay used for quantifying cell number in culture.

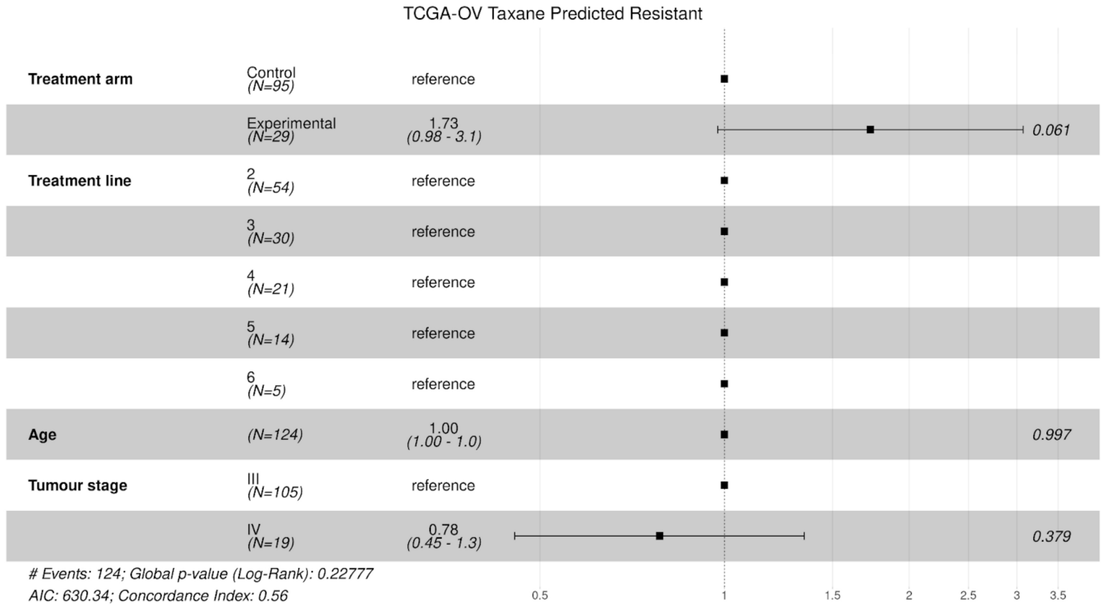


**Supplementary Figure 44.** Cox proportional hazards model for patients predicted as resistant to taxane in TCGA-OV, expanded from Figure 3b.

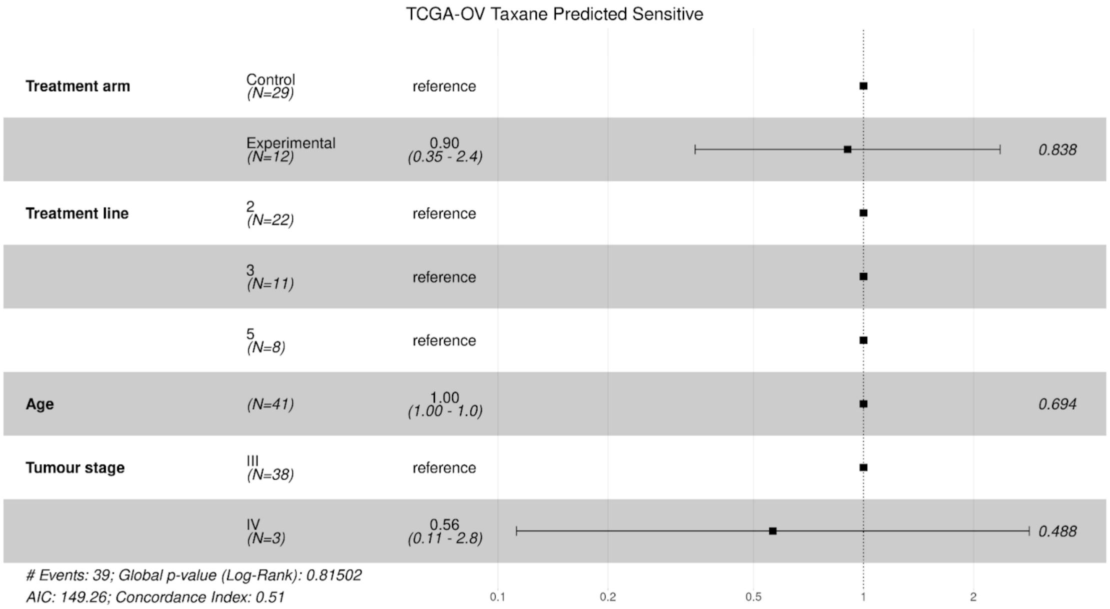


**Supplementary Figure 45.** Cox proportional hazards model for patients predicted as sensitive to taxane in TCGA-OV, expanded from Figure 3b.

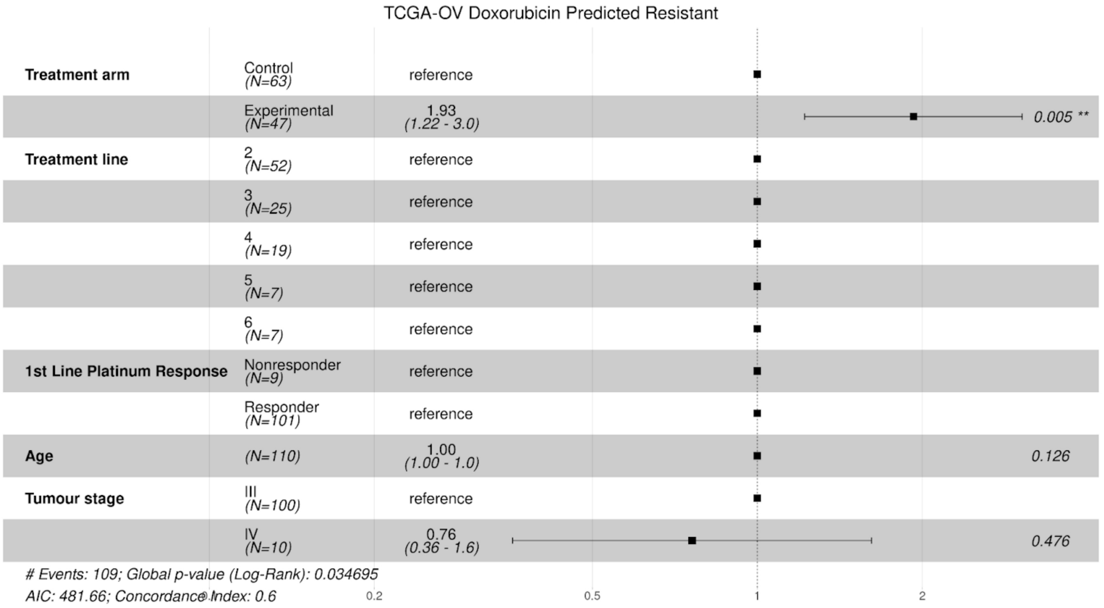


**Supplementary Figure 46.** Cox proportional hazards model for patients predicted as resistant to doxorubicin in TCGA-OV, expanded from Figure 3b.

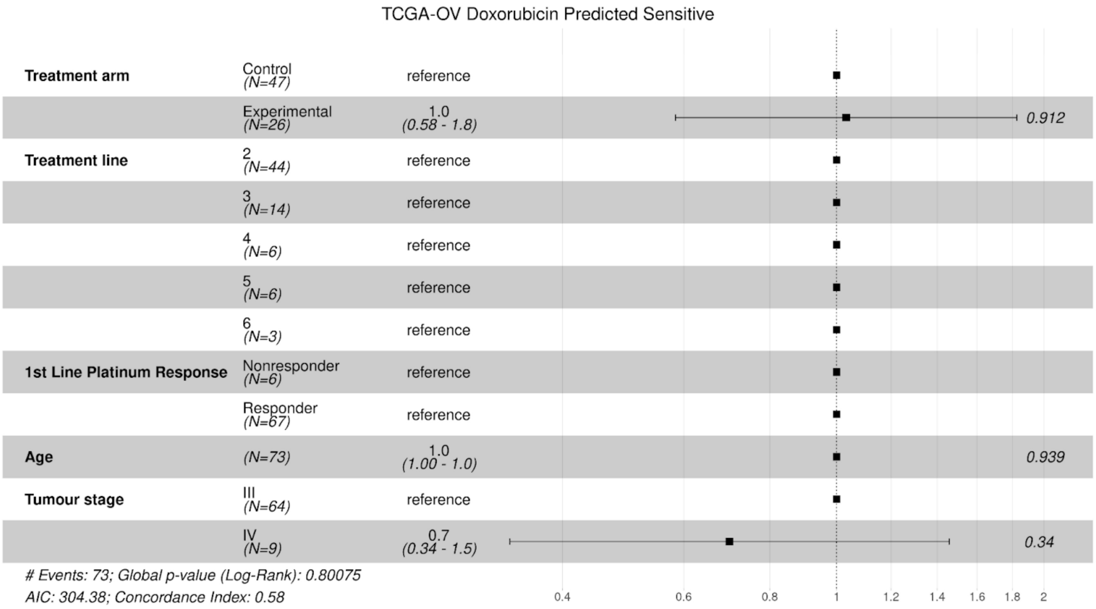


**Supplementary Figure 47.** Cox proportional hazards model for patients predicted as sensitive to doxorubicin in TCGA-OV, expanded from Figure 3b.

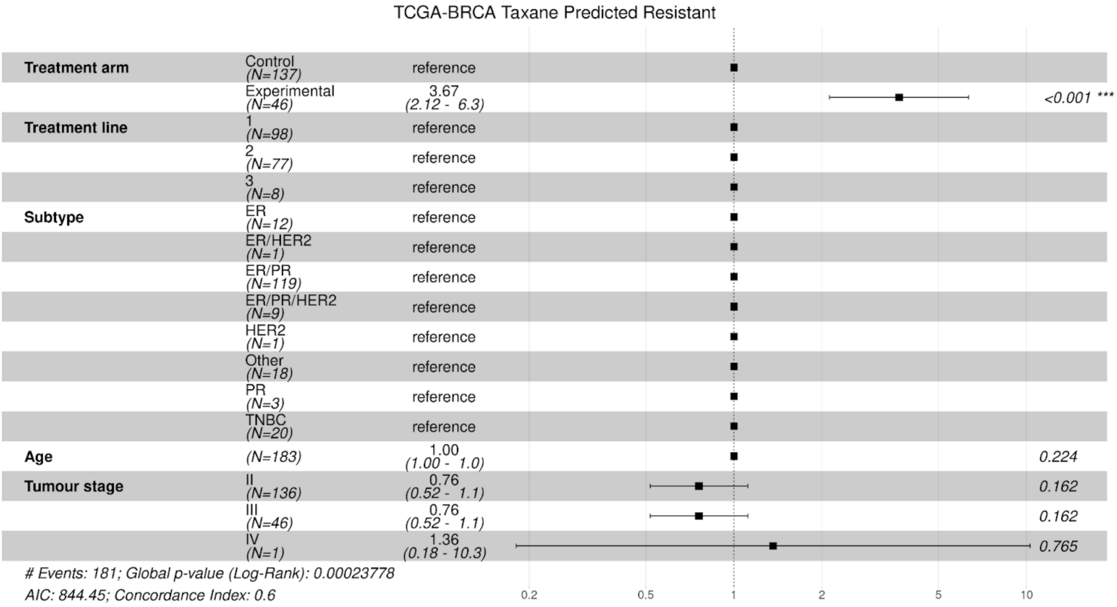


**Supplementary Figure 48.** Cox proportional hazards model for patients predicted as resistant to taxane in TCGA-BRCA, expanded from Figure 3b.

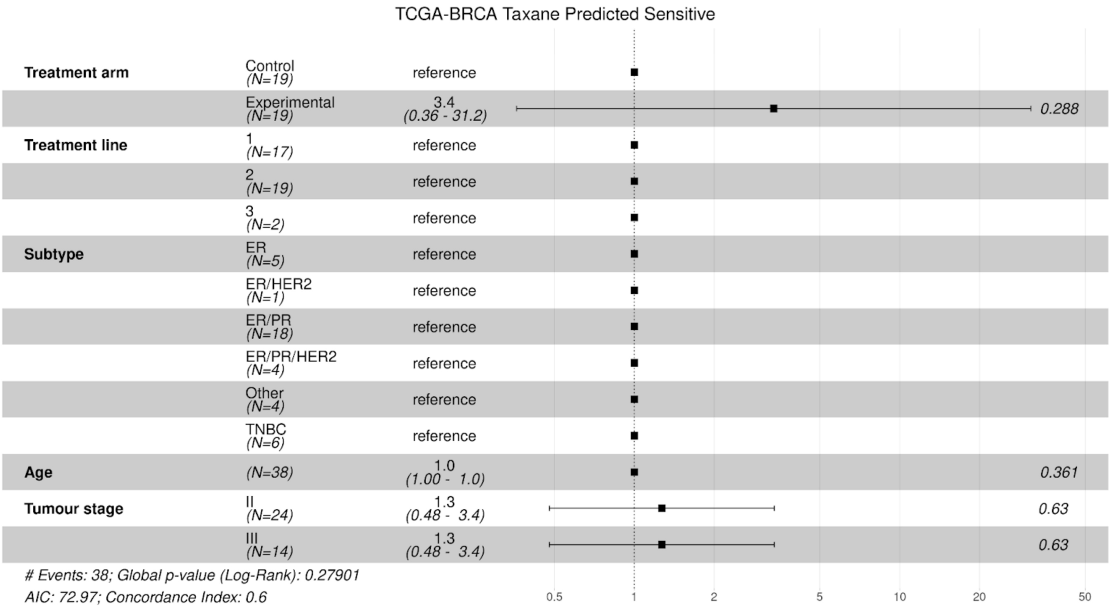


**Supplementary Figure 49.** Cox proportional hazards model for patients predicted as sensitive to taxane in TCGA-OV, expanded from Figure 3b.

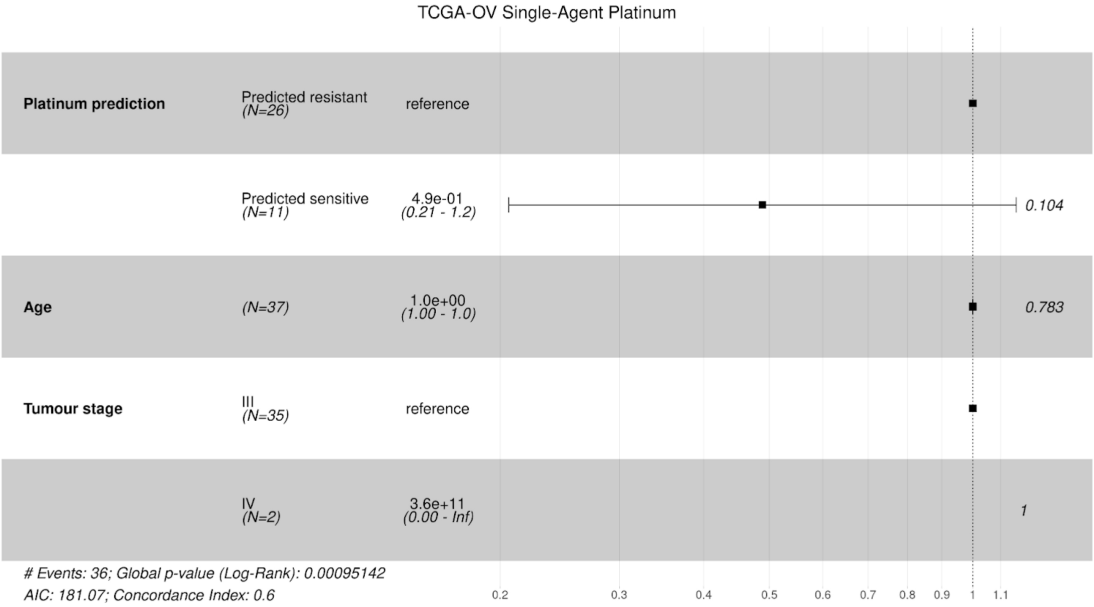


**Supplementary Figure 50.** Cox proportional hazards model for patients treated with single-agent platinum in TCGA-OV, expanded from Figure 3d.

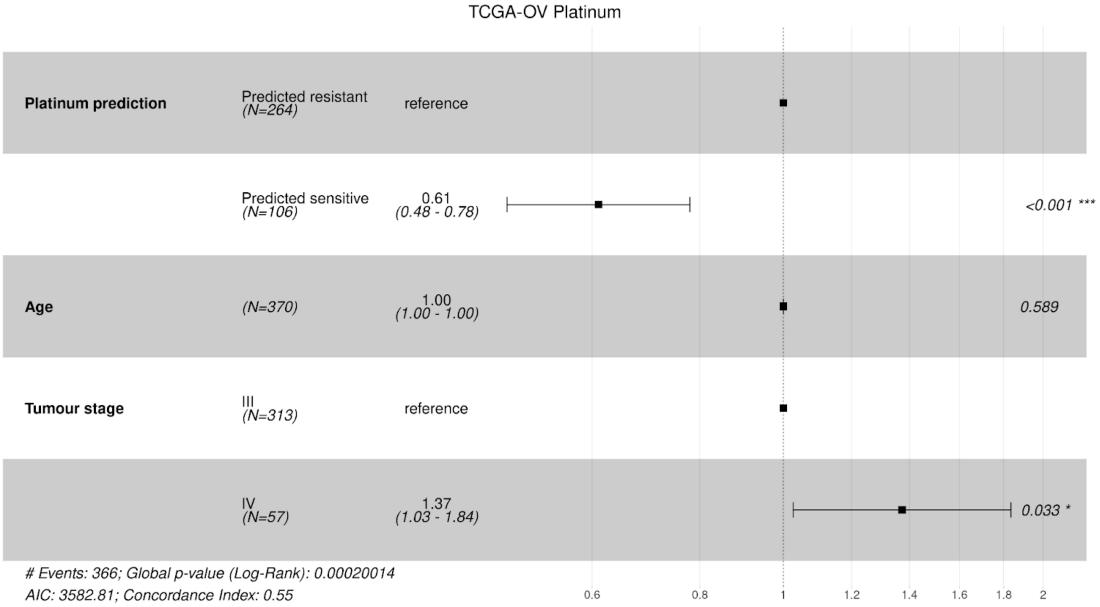


**Supplementary Figure 51.** Cox proportional hazards model for patients treated with platinum in TCGA-OV, expanded from Figure 3d.

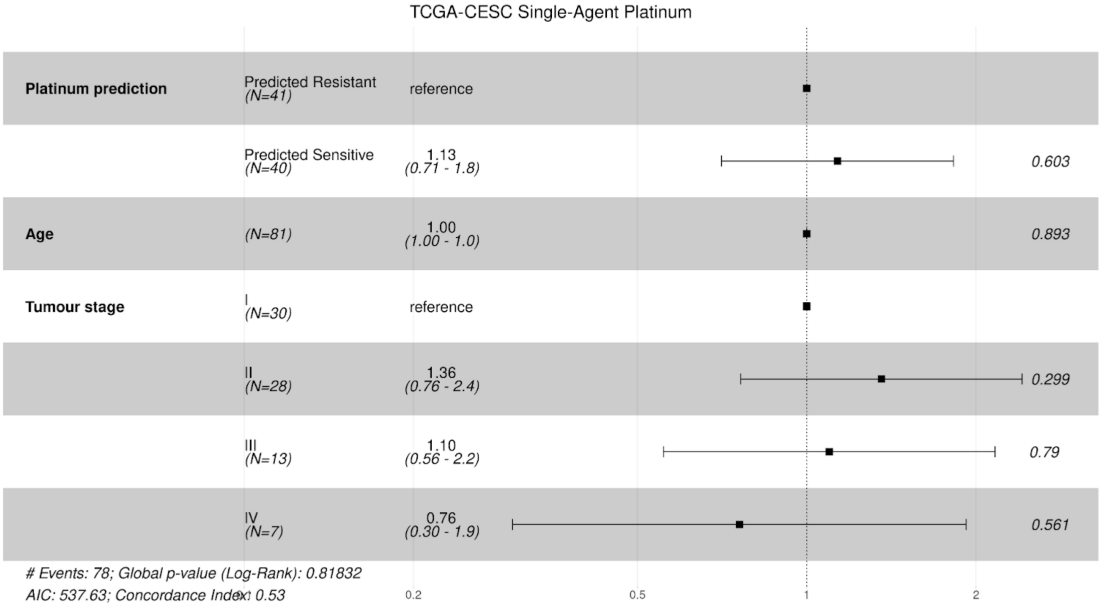


**Supplementary Figure 52.** Cox proportional hazards model for patients treated with single-agent platinum in TCGA-CESC, expanded from Figure 3d.

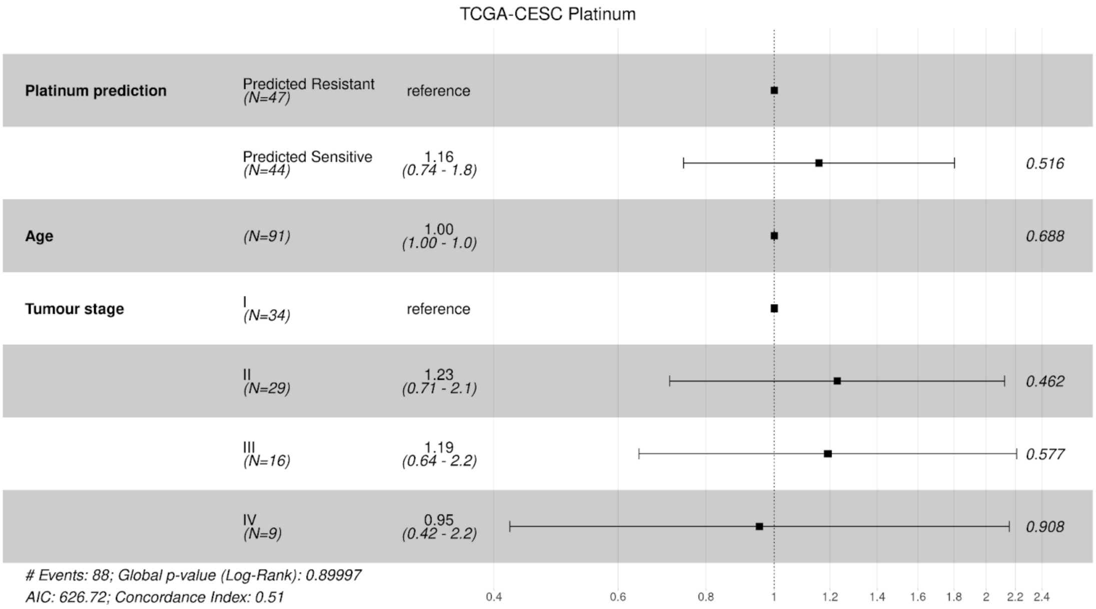


**Supplementary Figure 53.** Cox proportional hazards model for patients treated with platinum in TCGA-CESC, expanded from Figure 3d.

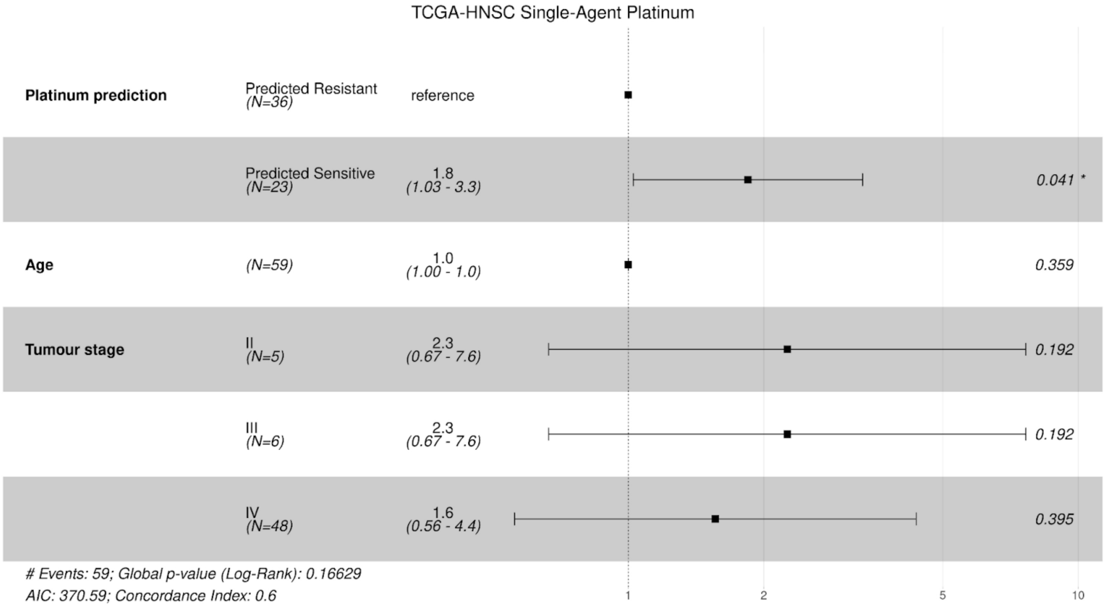


**Supplementary Figure 54.** Cox proportional hazards model for patients treated with single-agent platinum in TCGA-HNSC, expanded from Figure 3d.

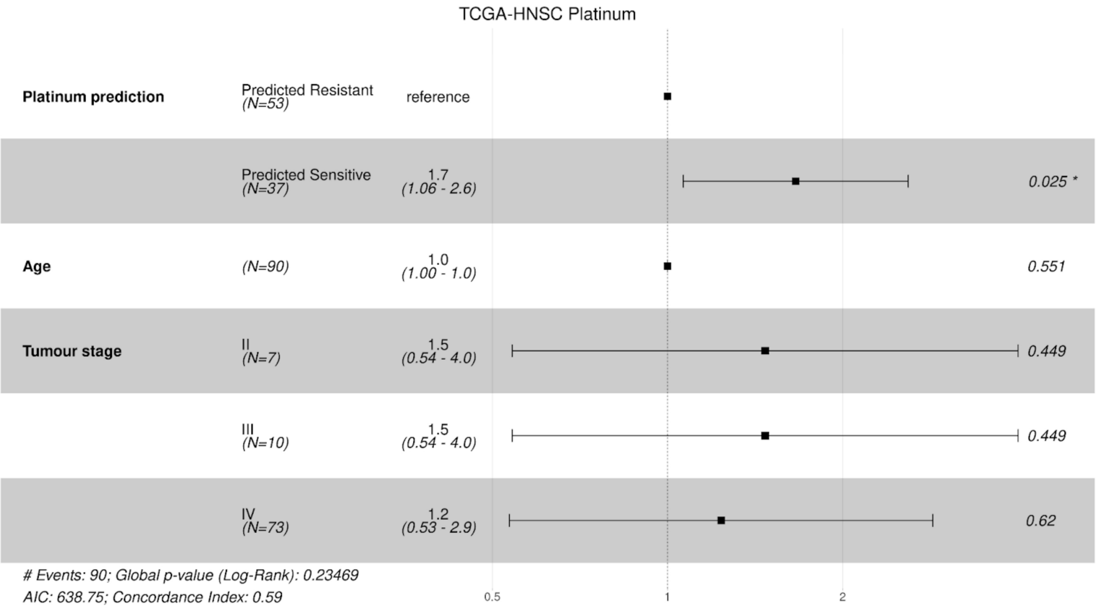


**Supplementary Figure 55.** Cox proportional hazards model for patients treated with platinum in TCGA-HNSC, expanded from Figure 3d.

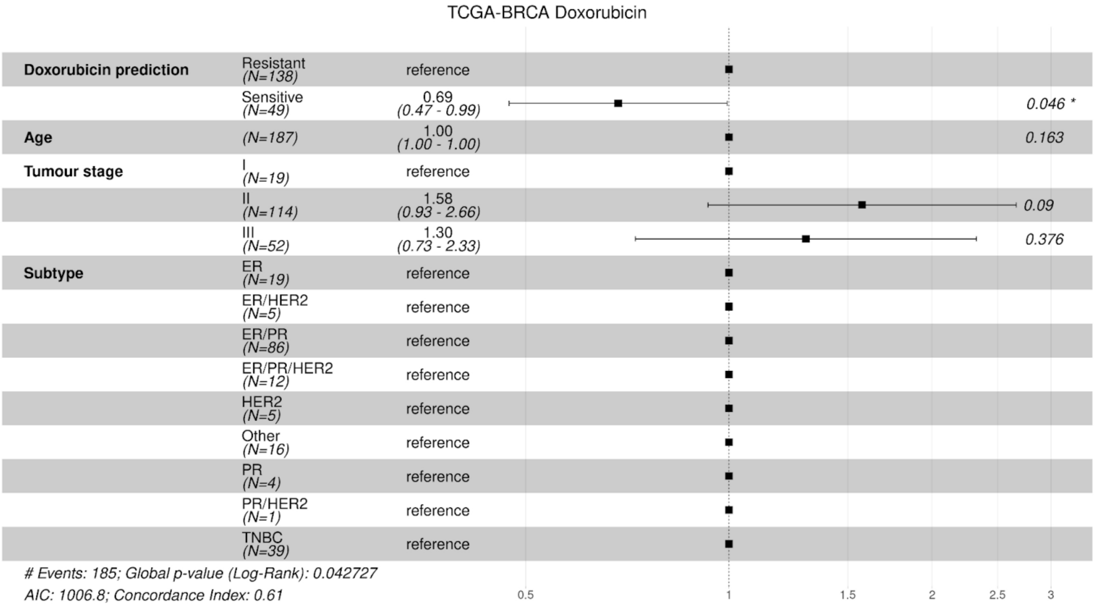


**Supplementary Figure 56.** Cox proportional hazards model for patients treated with doxorubicin in TCGA-BRCA, expanded from Figure 3d.

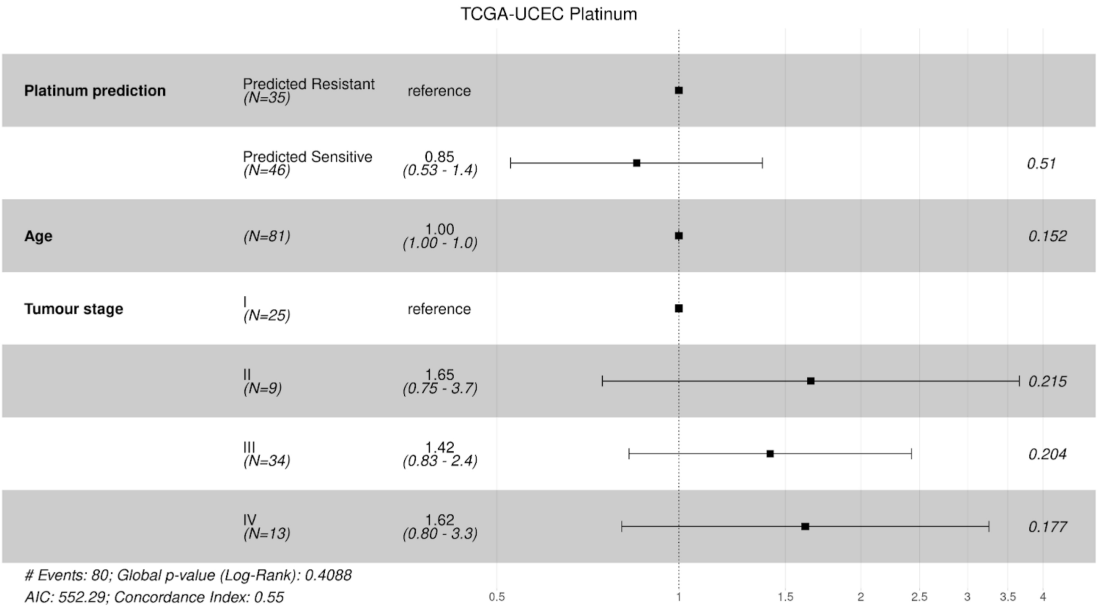


**Supplementary Figure 57.** Cox proportional hazards model for patients treated with platinum in TCGA-UCEC, expanded from Figure 3d.

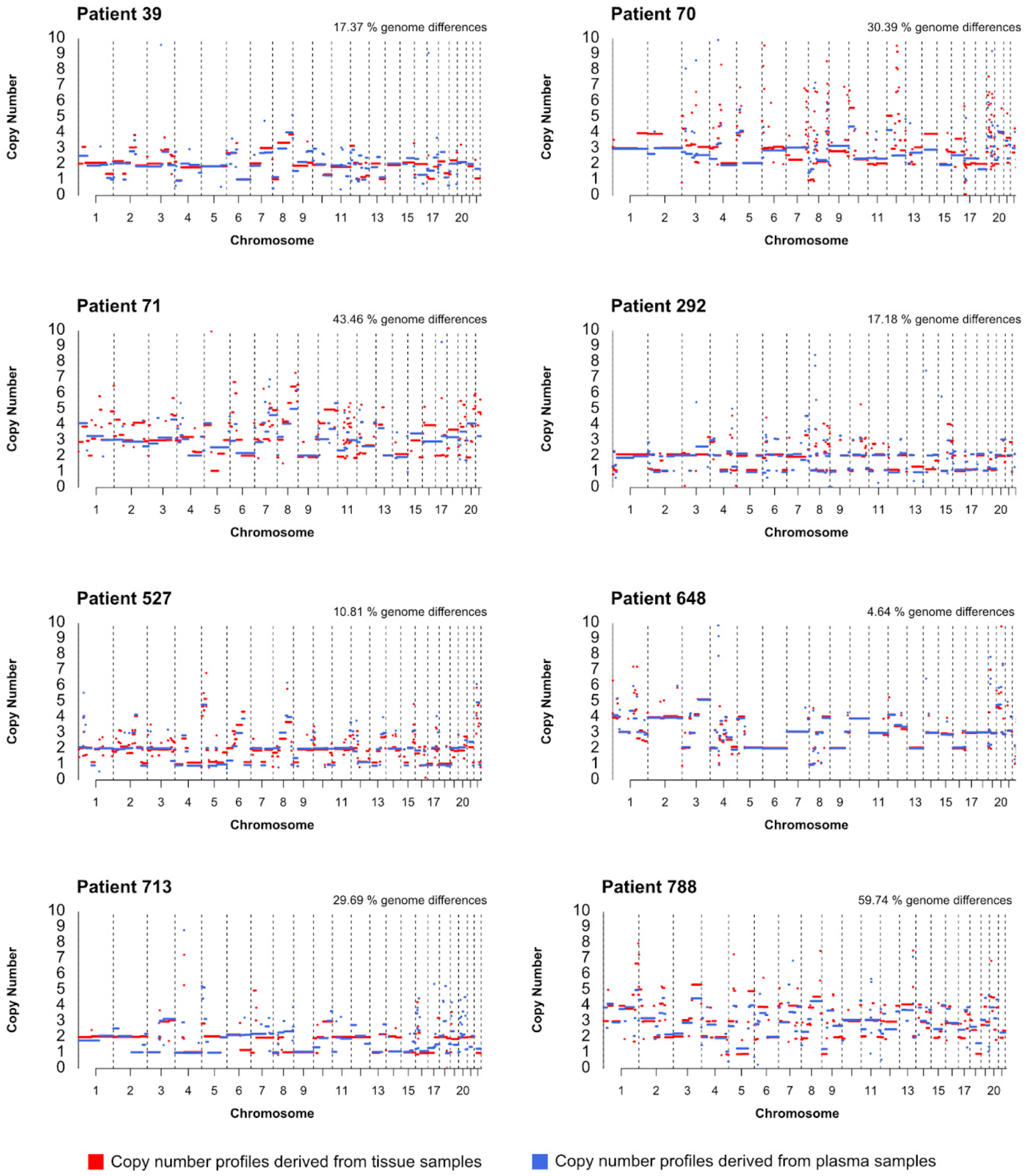


**Supplementary Figure 58. Comparison of copy number profiles derived from tissue or liquid biopsy.** Matched tissue- and plasma-derived copy number profiles from the patient 139 are shown in Figure 4b. The quantification of the extent of genome differences between pairs was calculated using the *CNpare* tool in R.

### Supplementary Tables

**Supplementary Table 1.** Clinical characteristics of the OV04 cohort

| | | **OV04** | |
|---|---|---|---|
| | | **Tissue (n=41)** | **Plasma (n=9)** |
| **Age at Diagnosis** | | | |
| | **<50** | 1 | 1 |
| | **50-60** | 9 | 2 |
| | **60-70** | 22 | 6 |
| | **70-80** | 8 | 0 |
| | **>80** | 1 | 0 |
| **Tumour Stage** | | | |
| | **Unknown** | 2 | 1 |
| | **I** | 2 | 0 |
| | **II** | 2 | 1 |
| | **III** | 20 | 2 |
| | **IV** | 15 | 5 |
| ***BRCA* Mutated** | | | |
| | ***BRCA1*** | 0 | 0 |
| | ***BRCA2*** | 0 | 0 |

**Supplementary Table 2.** Linear regression model for correlating response to paclitaxel (AUC) with activity levels of CX3 and CX5 signatures in 537 cell lines.

| Predictors | Estimates | CI | p-value |
|---|---|---|---|
| (Intercept) | 0.46 | [0.42, 0.50] | <0.001 |
| CX3 | -0.18 | [-0.33, -0.02] | 0.026 |
| CX5 | -0.40 | [-0.79, 0.00] | 0.051 |
| CX3 * CX5 | 0.82 | [-0.39, 2.04] | 0.184 |
| Observations | 291 | | |
| $R^2$ adjusted | 0.034 | | |

**Supplementary Table 3.** Platinum sensitivity status for organoid and spheroid samples

| **Patient ID** | **Sample type** | **Sample ID** | **Platinum sensitivity** |
|---|---|---|---|
| 75 | organoid | 23868org | sensitive |
| 297 | organoid | 54276org | resistant |
| 366 | organoid | 32077org | sensitive |
| 409 | organoid | 54288org | resistant |
| 413 | organoid | 54327org | sensitive |
| 466 | organoid | 118976org | sensitive |
| 571 | organoid | 151773org | sensitive |
| 627 | organoid | 119127org | resistant |
| 788 | organoid | 119178org | resistant |
| 920 | organoid | 151723org | resistant |
| 333 | spheroid | 118947 | sensitive |
| 338 | spheroid | 54356 | sensitive |
| 364 | spheroid | 32072 | resistant |
| 409 | spheroid | 54289 | resistant |
| 413 | spheroid | 54327 | sensitive |
| 466 | spheroid | 118976 | sensitive |
| 525 | spheroid | 80720 | sensitive |
| 626 | spheroid | 119016 | resistant |

| | | | |
|---|---|---|---|
| 648 | spheroid | 80601 | resistant |
| 669 | spheroid | 80630 | sensitive |
| 687 | spheroid | 118902 | resistant |
| 788 | spheroid | 119178 | resistant |
| 800 | spheroid | 119120 | resistant |
| 839 | spheroid | 119025 | sensitive |
| 875 | spheroid | 119136 | sensitive |

**Supplementary Table 4.** R packages used throughout the project.

| Name | Version | Citation | Name | Version | Citation |
|---|---|---|---|---|---|
| annotate | 1.72.0 | [1] | iterators | 1.0.14 | [2] |
| AnnotationDbi | 1.56.2 | [3] | lattice | 0.20.45 | [4] |
| Biobase | 2.54.0 | [5] | locfit | 1.5.9.8 | [6] |
| BiocGenerics | 0.40.0 | [5] | lsa | 0.73.3 | [7] |
| BiocManager | 1.30.19 | [5] | magrittr | 2.0.3 | [8] |
| broom | 1.0.3 | [9] | MASS | 7.3.58.1 | [10] |
| CINSignature Quantification | 1.1.1 | Repo link | MatrixGenerics | 1.6.0 | [11] |
| CNpare | 0.99.0 | [12] | matrixStats | 0.63.0 | [13] |
| ComplexHeatmap | 2.10.0 | [14] | mclust | 6.0.0 | [15] |
| data.table | 1.14.8 | [16] | org.Hs.eg.db | 3.14.0 | [17] |
| DESeq2 | 1.34.0 | [18] | QDNAseq | 1.30.0 | [19] |
| doMC | 1.3.8 | [20] | QDNAseqmod | 1.21.0 | Repo link |
| doParallel | 1.0.17 | [21] | readr | 2.1.4 | [22] |
| dplyr | 1.1.0 | [23] | readxl | 1.4.2 | [24] |
| drc | 3.0.1 | [25] | remotes | 2.4.2 | [26] |
| fgsea | 1.20.0 | [27] | reshape2 | 1.4.4 | [28] |

| | | | | | |
|---|---|---|---|---|---|
| flexmix | 2.3.18 | [29] | S4Vectors | 0.32.4 | [30] |
| foreach | 1.5.2 | [20] | SnowballC | 0.7.1 | [31] |
| genefilter | 1.76.0 | [32] | stats4 | 4.1.2 | [33] |
| geneplotter | 1.72.0 | [34] | Summarized Experiment | 1.24.0 | [35] |
| GenomeInfoDb | 1.30.1 | [36] | survival | 3.5-5 | [37] |
| GenomicData Commons | 1.18.0 | [38] | survminer | 0.4.9 | [39] |
| GenomicRanges | 1.46.1 | [40] | TCGAbiolinks | 2.22.4 | [41] |
| geomnet | 0.3.1 | Repo link | this.path | 1.2.0 | [42] |
| ggalluvial | 0.12.5 | [43] | tidyr | 1.3.0 | [44] |
| ggplot2 | 3.4.1 | [45] | tidyverse | 1.3.2 | [46] |
| ggpubr | 0.6.0 | [47] | XML | 3.99.0.13 | [48] |
| ggthemes | 4.2.4 | [49] | xml2 | 1.3.3 | [50] |
| grid | 4.1.2 | [51] | YAPSA | 1.20.1 | [52] |
| IRanges | 2.28.0 | [40] | | | |

**Supplementary Table 5.** Sources of data used in this study

| **Data** | **Source** |
|---|---|
| OV04 clinical and raw data | Generated in this project |
| OV04 copy number profiles | Generated in this project |
| Ovarian cancer cell lines | Derived in-house (CIOV1, CIOV2,CIOV4, CIOV6), American Type Culture Collection (OVCAR3), gifted by the Langdon Lab (PEO23) |
| Cell line, organoid, and spheroid doxorubicin dose-response data | Generated in this project |
| TCGA copy number profiles | Drews et al 2022. Files hosted on http://github.com/VanLoo-lab/ascat |

| | |
|---|---|
| TCGA clinical data | Genomic Data Commons Data Portal: https://portal.gdc.cancer.gov/ |
| Transcriptomics data for the TCGA ovarian cancer samples | Genomic Data Commons Data Portal: https://portal.gdc.cancer.gov/ |
| TTF data for TCGA ovarian cancer samples | [53] |
| Triple negative status for TCGA breast cancer samples | [54], accessed via https://github.com/TransBioInfoLab/TNBC_analysis |
| Triple negative status for TCGA breast cancer samples | [55] |
| Triple negative status for TCGA breast cancer samples | [56] |
| ER/PR/HER2 status for TCGA breast cancer samples | [57] |
| ER/PR/HER2 status for TCGA breast cancer samples | [56] |
| Hallmark pathways for differential expression analysis (h.all.v7.1.symbols.gmt) | Human MSigDB Collections: https://www.gsea-msigdb.org/gsea/msigdb/index.jsp |
